# Dilated cardiomyopathy-associated skeletal muscle actin (ACTA1) mutation R256H disrupts actin structure and function and causes cardiomyocyte hypocontractility

**DOI:** 10.1101/2024.03.10.583979

**Authors:** Ankit Garg, Silvia Jansen, Rui Zhang, Kory J. Lavine, Michael J. Greenberg

## Abstract

Skeletal muscle actin (ACTA1) mutations are a prevalent cause of skeletal myopathies consistent with ACTA1’s high expression in skeletal muscle. Rare *de novo* mutations in ACTA1 associated with combined cardiac and skeletal myopathies have been reported, but ACTA1 represents only ∼20% of the total actin pool in cardiomyocytes, making its role in cardiomyopathy controversial. Here we demonstrate how a mutation in an actin isoform expressed at low levels in cardiomyocytes can cause cardiomyopathy by focusing on a unique ACTA1 mutation, R256H. We previously identified this mutation in multiple family members with dilated cardiomyopathy (DCM), who had reduced systolic function without clinical skeletal myopathy. Using a battery of multiscale biophysical tools, we show that R256H has potent functional effects on ACTA1 function at the molecular scale and in human cardiomyocytes. Importantly, we demonstrate that R256H acts in a dominant manner, where the incorporation of small amounts of mutant protein into thin filaments is sufficient to disrupt molecular contractility, and that this effect is dependent on the <u>presence</u> of troponin and tropomyosin. To understand the structural basis of this change in regulation, we resolved a structure of R256H filaments using Cryo-EM, and we see alterations in actin’s structure that have the potential to disrupt interactions with tropomyosin. Finally, we show that *ACTA1^R256H/+^* human induced pluripotent stem cell cardiomyocytes demonstrate reduced contractility and sarcomeric disorganization. Taken together, we demonstrate that R256H has multiple effects on ACTA1 function that are sufficient to cause reduced contractility and establish a likely causative relationship between ACTA1 R256H and clinical cardiomyopathy.

**Significance Statement:** Skeletal muscle actin mutations are well-known to cause skeletal myopathies, but their role in cardiomyopathies have been controversial as skeletal muscle actin is only expressed at modest levels in the heart. Here, we demonstrate that a skeletal muscle actin mutation potently causes multiple defects in actin function at the atomic and molecular scales, and it functions in a dominant fashion, leading to cardiomyocyte contractile defects. Our results establish how skeletal muscle actin mutations may cause cardiomyocyte dysfunction and lay the foundation for future studies of the role of skeletal muscle actin in cardiomyopathy.

## Introduction

Dilated cardiomyopathy (DCM) is a leading cause of heart failure and left ventricular assist device/cardiac transplantation (1). DCM can be both secondary to another pathologic process, but it also has numerous genetics causes, including mutations in sarcomeric proteins (2). DCM mutations associated with sarcomeric proteins have poorer outcomes to guideline directed interventions (3) which suggests mechanistic differences in the pathogenesis of these patients’ DCM. As our appreciation for the mechanistic heterogeneity of DCM grows, it becomes imperative to utilize multiscale analysis to study unique phenotypes so we may better utilize available therapies and identify new targets (4).

While some genes have been unambiguously associated with DCM including cardiac myosin and titin, there is ambiguity about the roles of other genes in DCM pathogenesis (5). In part, this ambiguity stems from the small number of patients with these mutations and a lack of understanding of the connections between variants and the disease phenotype. One such gene with a controversial role in DCM is skeletal muscle actin, ACTA1. Actin is one of the most conserved and highly expressed proteins in eukaryotes. Six isoforms of actin are expressed in humans, and two isoforms, ACTA1 and ACTC1 (referred to as “cardiac actin”), are specific to sarcomeres (6). Actin is a principal component of the sarcomere thin filament, where it binds to troponin and tropomyosin, which regulate the calcium-dependent interactions between myosin and the thin filament (4). Mutations perturbing contraction can cause skeletal myopathies and cardiomyopathies (7). While ACTC1 is expressed exclusively in the adult heart, ACTA1 is expressed at high levels in skeletal muscle and an order of magnitude less in cardiac muscle where ACTC1 expression dominates (6). Given this specificity, multiple mutations in ACTA1 have been associated with skeletal myopathies while ACTC1 mutations have been associated with cardiomyopathies. With the low expression of ACTA1 in the heart and high expression in skeletal muscle, some have speculated that ACTA1 mutations would have no impact on cardiac contractility (8); however, rare cases of DCM with combined skeletal myopathy have since been identified in patients bearing ACTA1 mutations (9–12). To date, no studies have established cardiac pathogenicity of these ACTA1 mutations nor the underlying mechanism.

We previously described the unique R256H (also referred to as R254H when using the post-translationally cleaved actin numbering convention; henceforth referred to as “post-cleavage” numbering) mutation in ACTA1 which was associated with DCM in *ACTA1^R256H/+^* patients but uniquely lacked a clinical skeletal myopathy typically seen with ACTA1 mutants (13). Unlike previous ACTA1-associated DCM mutations which were *de novo* mutations affecting single patients, this report demonstrated an ACTA1 DCM mutation in multiple patients within a family. There was a subsequent report of a similar ACTA1 mutation, G253S, that also appeared to be associated with DCM without skeletal myopathy (14). Though these mutations provided evidence for ACTA1 having a role in cardiomyopathy, it was unknown how a low expressed actin isoform in the heart could affect cardiac contractility.

Here, we used multiscale computational and experimental tools spanning from atoms to cells to understand the mechanism of the ATCA1 R256H mutation. Our results demonstrate that R256H perturbs multiple actin functions, including a potent effect on contractility, and they provide new insights into how mutations in the skeletal muscle actin isoform can cause cardiomyopathy.

## Results

### Molecular dynamics simulations show that R256H alters the structural dynamics of monomeric ACTA1

To gain insights into the atomic-level changes induced by the R256H mutation, we used molecular dynamics simulations. The structure of the ACTA1 monomer has been extensively modeled (15) and used in molecular dynamics simulations (16). For our starting structure, we utilized the monomeric structure of ACTA1 bound to ADP and calcium (PDB 1J6Z) (17). All our simulations were performed in explicit water. We ran five separate 1 μs simulations for the WT ACTA1. For the R256H mutant, a histidine is introduced into the structure, and at physiological pH, His residues can exist in 3 separate states: “HSD” neutrally Nδ protonated, “HSE” neutrally Nε protonated, and “HSP” doubly protonated His. Therefore, we also ran five separate 1 μs simulations for simulations of each of these possibilities (R256HSD, R256HSE, and R256HSP).

To quantify the differences between structures, we calculated the root mean square displacement (RMSD) of the structures relative to the average WT structure. The RMSDs for each protonation state of the mutant were significantly different from WT ACTA1 (**Figures 1A and 1B**), demonstrating that the mutation affects the mean structure of ACTA1, regardless of the protonation state. To better understand where these changes might arise, we calculated the RMSD per residue compared to the average WT structure. Residues with the greatest change in RMSD appeared to concentrate to subdomains 2 and 4 (**Figure 1C**), suggesting that R256H induces allosteric changes that affects both these subdomains. Interestingly, in many of our simulations, we observed the loss of ADP. Since subdomains 2 and 4 are close to each other in the ADP bound state, we examined whether the mutant affects the time to the loss of nucleotide. We found that time to ADP dissociation was significantly faster for the R256HSE and R256HSP ACTA1 compared to WT ACTA1, and R256HSD trended towards faster ADP dissociation (**Figures 1D and 1E**). Taken together, our simulations predict that R256H changes the structural dynamics of ACTA1, potentially affecting nucleotide binding.

**Figure 1.**
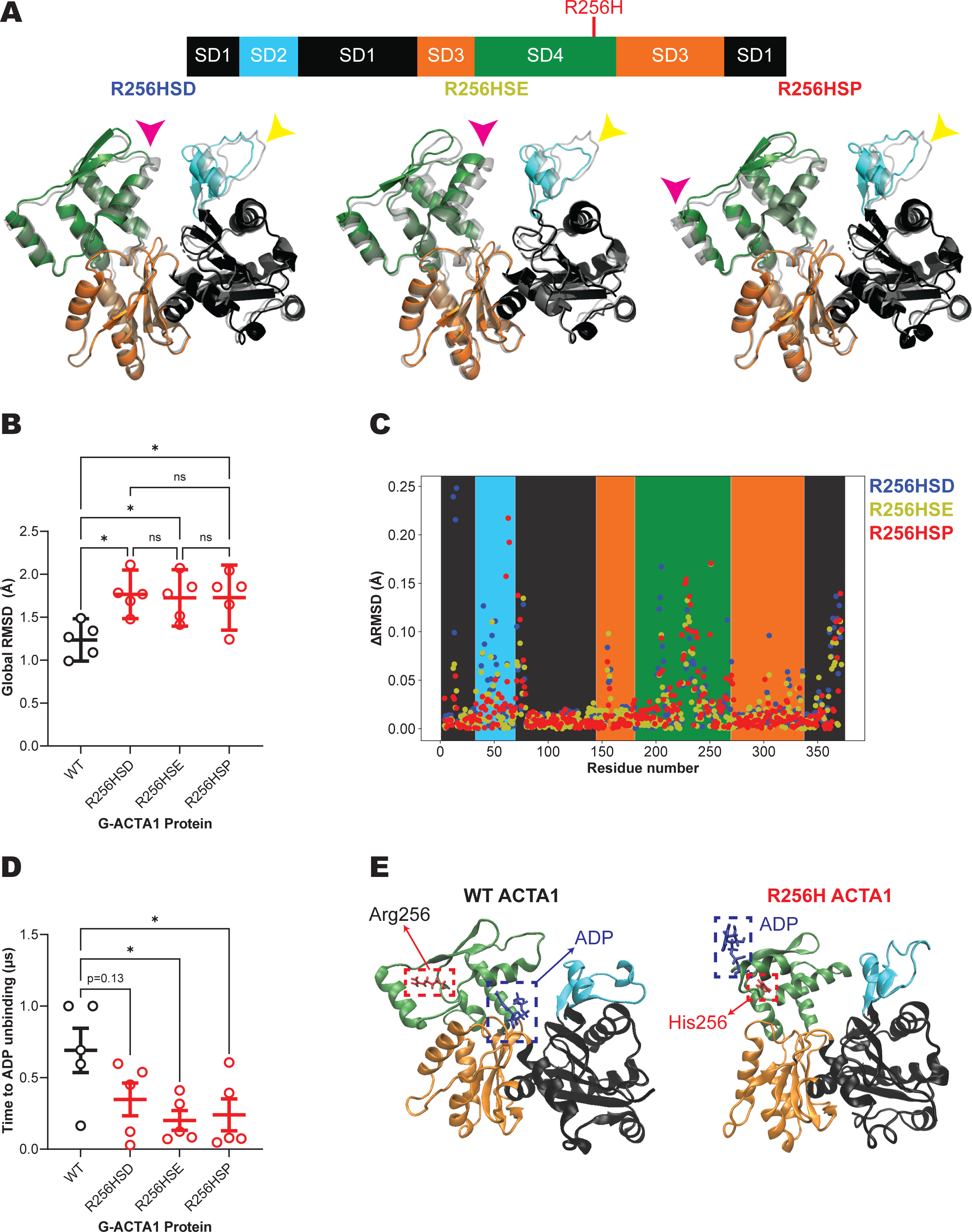
Simulations of G-ACTA1 R256H predict differences in structure and decreased ADP binding stability. **A)** (Top) Cartoon representing the domain architecture of ACTA1 with individual subdomains (“SD”) colored to correspond to colors seen throughout the figure. The position of the R256H mutation is indicated in red. (Bottom) Superimposed average structures of R256HSD, R256HSE, and R256HSP ACTA1 (solid colors) on the average structure of WT ACTA1 (transparent grey color) obtained from five 1 μs simulations of each condition. Note the differences in subdomain 4 (magenta arrow) and subdomain 2 (yellow arrow) in R256H ACTA1. **B)** The global RMSD of mutant ACTA1 is different from WT ACTA1 indicating an overall difference in structure. RMSD is calculated relative to the average WT structure from five 1 μs simulations. For example, “WT” shows the RMSD difference between each WT simulation averaged structure compared to the averaged structure of all WT simulations. * = p < 0.05 as calculated by ANOVA with multiple comparison testing followed by Tukey correction. **C)** Scatter plot of mean difference in RMSD between R256H ACTA1 and WT ACTA1. Error bars are not shown for clarity. Data are not significant on a per-residue basis after performing two-way ANOVA with multiple comparisons followed by Dunnett correction. **D)** ACTA1 R256H binding time to ADP is shorter than ACTA1 WT. Binding time is defined as the amount of time that the ADP molecule lies within 20 Å of the center of mass of ACTA1. Data are obtained from the same simulations as above. * = p < 0.05 as calculated by ANOVA with multiple comparison testing followed by Dunnett correction. **E)** Representative single frame captures from molecular dynamics simulations of the ACTA1 1J6Z starting structure in the WT (left) and R256H mutated (right) states. Shown in the red dashed box is the 256 residue and in the blue dashed box is the ADP nucleotide. Note the loss of ADP from the binding pocket in R256H. All error bars represent standard error of the mean (SEM).

### R256H reduces actin stability and impairs actin polymerization

Our molecular dynamics simulations predict that R256H affects the structure of the actin monomer, leading to destabilization of nucleotide binding. It is well established that nucleotide binding is important to G-actin stability and polymerization (18). Therefore, we set out to test the effects of R256H ACTA1 on thermal stability and actin polymerization. Critically, testing our hypotheses, requires R256H ACTA1 protein; however, recombinant ACTA1 has been difficult to purify in sufficient quantities and purity. We adapted an SF9/baculovirus system that has been successfully used to produce other α-actins including mutant actins (19–22). Previous SF9 ACTA1 purification strategies either overexpressed untagged ACTA1 which could co-polymerize with endogenous SF9 actin, resulting in decreased purity, or they utilized a non-polymerizable tag whose proteolytic cleavage left multiple non-native amino acids (23). We employed an expression strategy utilizing a chymotrypsin-cleavable C-terminal tag composed of thymosin β4 (TMSB4) to block polymerization in tandem with a multi-histidine chain that has been successfully used to purify functional wildtype and mutant ACTA2 (smooth muscle actin) (24), which is similar to ACTA1 (**Supplementary Figure 1**). We successfully applied this system to generate both WT and R256H ACTA1 recombinant proteins with yields ranging from 0.4 to 0.9 mg per 250 mL of SF9 culture.

To test whether globular (i.e. monomeric) ACTA1 R256H had altered thermal stability, we used a protein thermal shift assay. We found that the melting temperature of R256H ACTA1 (46.1°C) was significantly lower than the melting temperature of WT ACTA1 (58.9°C) (**Figure 2A**), consistent with our predictions from the simulations.

**Figure 2.**
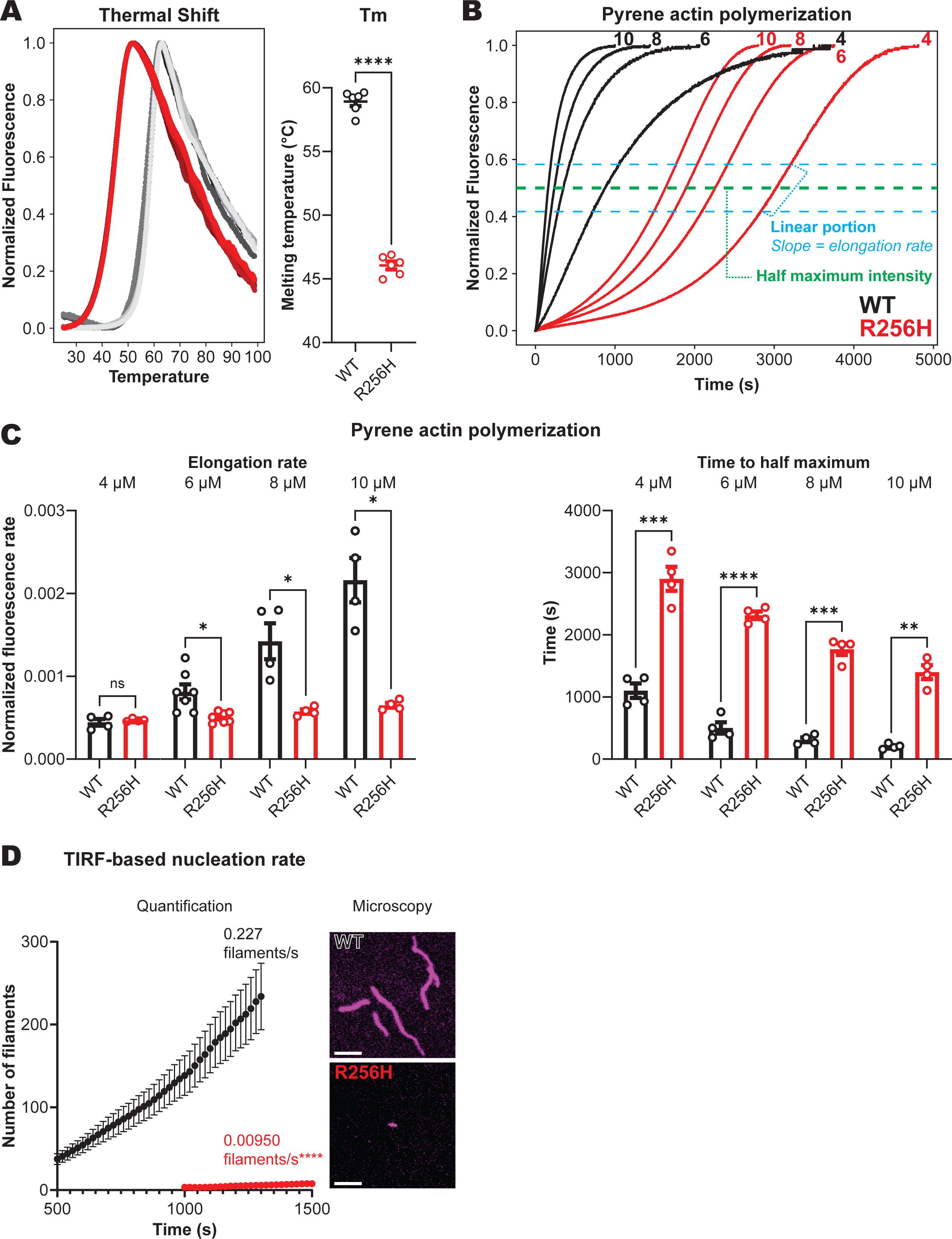
R256H ACTA1 has decreased thermal stability and disrupted polymerization. **A)** (Left graph) Normalized fluorescence generated from melting WT and R256H proteins using Thermo Fisher Scientific Protein Thermal Shift™ assay. As protein denatures, the dye binds previously inaccessible hydrophobic regions resulting in increased fluorescent signal. Six traces are present for each sample, but not all 6 traces are visible due to high similarity between data. (Right graph) Calculated melting temperature. The experiment was performed in triplicate from two independently purified sets of ACTA1 WT and R256H protein (n=6 for each sample). **** = p < 0.0001 by two-tailed T-test. **B)** Representative traces of pyrene polymerization assays performed at various actin monomer concentrations for WT and R256H ACTA1 (each number above each curve denotes concentration in μM). Fluorescent signal increases as actin polymerizes. Note the substantial difference in the kinetics of polymerization between WT and R256H. These are quantified in the proceeding panels as differences in **C)** elongation rate and time to half maximum intensity. These values were derived from the regions shown in cyan shown in “B”. *N*=4-7 and the individual traces used to derive the data in “C” are shown in **Supplementary Figure 2**. * = p < 0.05, ** = p < 0.01 as calculated by two-tailed T-test with Welch’s correction for unequal variance. **D)** R256H ACTA1 has decreased nucleation rate. Number of filaments detected and quantified using Fiji plugin, FilamentDetector. **** p < 0.0001 by simple linear regression analysis in GraphPad Prism. Image insets are examples of fluorescent filaments at 1000 seconds. Note that images were post-processed for visualization purposes but raw images were used for analysis. Full focal field can be found in **Supplementary Figure 3**. Scale bar represents 5 μm. All error bars represent SEM.

Next, we measured the kinetics of polymerization using pyrene-labeled actin which shows increased fluorescence when actin is integrated into a filament (25). The time course of polymerization has been extensively studied, and follows a typical sigmoidal relationship, consisting of an initial slow growth lag phase whose duration is related to actin nucleation, a rapid linear growth phase largely driven by filament elongation, and then a steady state phase (26). Consistent with previous reports, the WT actin clearly follows this relationship, and, in line with nucleation being the rate-limiting step, the lag phase becomes shorter at higher actin concentrations (**Figure 2B**). Compared to WT, R256H ACTA1 demonstrated altered polymerization kinetics. Its lag phase, which reflects the time to nucleation, is prolonged even at the highest concentrations of actin tested. Moreover, the elongation rate, calculated as the slope of the linear phase, was slower for R256H ACTA1 than WT ACTA1 over the entire concentration range tested (**Figure 2C**, left graph). With these two effects combined, the R256H ACTA1 has an increased time to half-maximum intensity (**Figure 2C**, right graph). To directly test whether the mutation affects actin nucleation as well as elongation, we employed total internal reflection (TIRF) microscopy to visualize growing actin filaments incorporating fluorescent actin monomers in real time. Consistent with the prolonged lag phase seen in our pyrene polymerization assay and reduced polymerization kinetics (**Figure 2B**), R256H ACTA1 had a significantly decreased rate of filament formation (**Figure 2D**) and showed significantly slower polymerization (**Supplementary Figure 3B**).

Our polymerization assays in Figure 2 utilized either WT or R256H ACTA1; however, all of the DCM patients observed were heterozygous for the mutation and therefore express both WT and mutant ACTA1 (13). To test whether R256H ACTA1 has a dominant negative effect on polymerization (i.e., the mutant protein acts as a poison protein and prevents polymerization), we repeated our polymerization experiments with mixing of both proteins. If dominant negative effects of R256H ACTA1 were present, we would expect that adding 3 μM R256H to 3 μM WT protein would inhibit polymerization compared to 3 μM WT alone. However, we saw that adding 3 μM R256H to 3 μM WT protein did not inhibit the polymerization kinetics compared to 3 μM WT (**Supplementary Figure 2E**). Moreover, we saw via TIRF microscopy that mixing WT and R256H ACTA1 resulted in an intermediate filament length and nucleation rate (**Supplementary Figure 3**). This demonstrates that mutant protein is being integrated into filaments without preventing polymerization, and that the mutant protein is not acting through a dominant negative mechanism that poisons polymerization. Rather, the mutant slows the kinetics of polymerization, both by itself and in the presence of WT protein. Taken together, our results demonstrate that, consistent with consequences of less nucleotide binding predicted by molecular dynamics simulations, the mutant protein affects both the stability and polymerization of actin.

### R256H ACTA1 filaments have disrupted actomyosin interactions in the presence of thin filament regulatory proteins

Our polymerization experiments demonstrated that both WT and R256H actin can form filaments, and therefore, we set out to examine how the mutation affects the function of actin filaments interacting with binding partners. The force and power of muscle contraction is driven by the myosin molecular motor pulling on regulated thin filaments consisting of actin and the regulatory proteins troponin and tropomyosin (4). Given that *ACTA1^R256H/+^* patients have reduced cardiac contractility, we hypothesized that R256H ACTA1 filaments would alter myosin-based interactions with regulated thin filaments. To test our hypothesis, we used an *in vitro* motility assay in which fluorescently-labeled actin/thin filaments are translocated over a bed of myosin in the presence of ATP and varying concentrations of calcium (27). For these experiments, we used porcine cardiac myosin (MYH7), which has biochemical and biophysical characteristics that are indistinguishable from human cardiac myosin (28–30). First, we examined whether the mutant protein affected the speed of actin translocation by myosin in the absence of regulatory proteins. We found that there was no difference in speed between WT and R256H ACTA1 filaments, demonstrating that myosin is capable of interacting with both WT or R256H ACTA1 filaments, and that the mutant protein does not affect the rate of myosin-based translocation (**Figure 3A**, left column pair).

**Figure 3.**
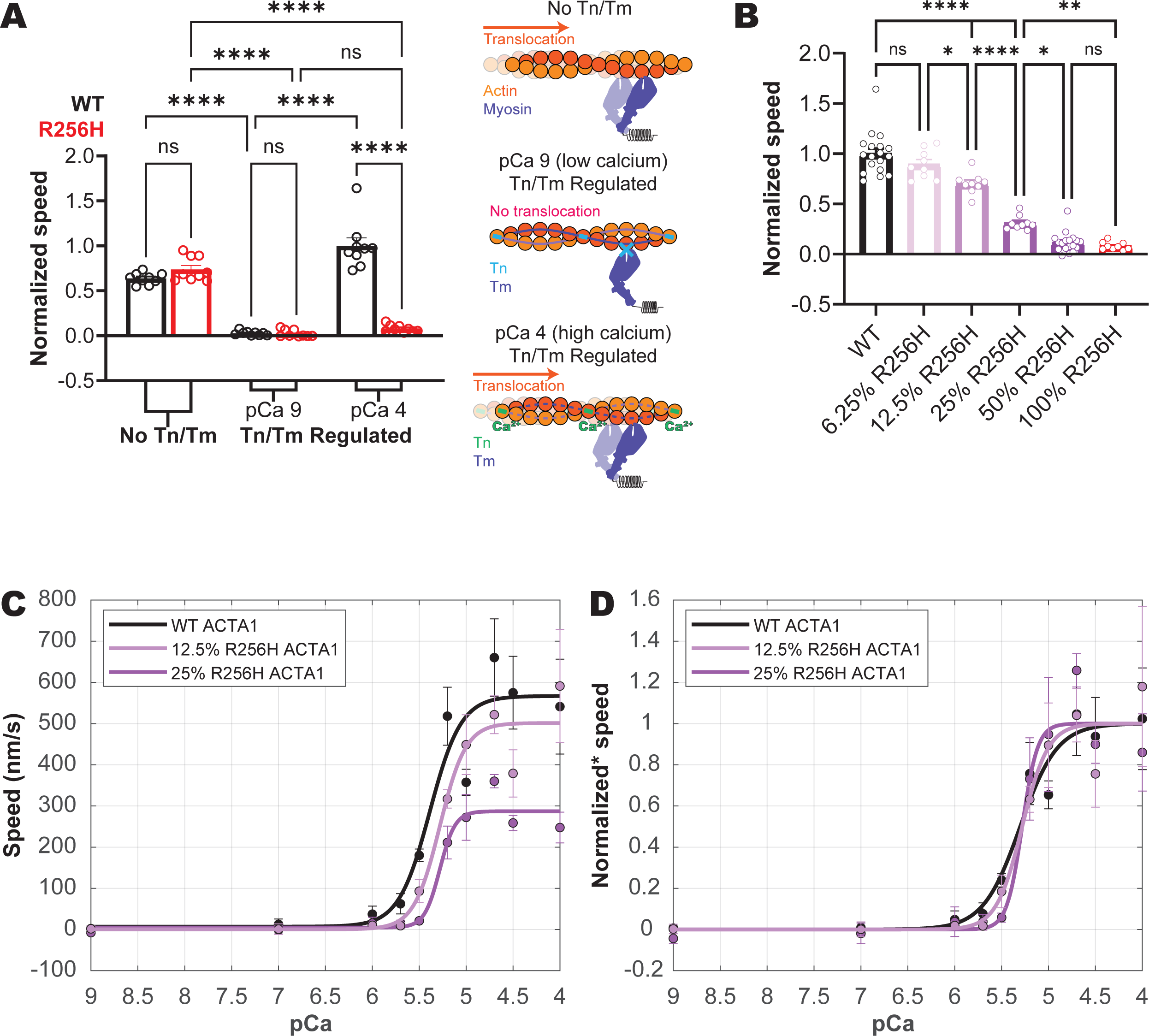
ACTA1 R256H filaments have inhibited translocation by myosin only in the presence of Tn/Tm without affecting calcium sensitivity. **A)** *In vitro* motility of either 100% WT or 100% R256H ACTA1 filaments with porcine cardiac myosin. Cartoons to the right of graphs are shown for conceptual visualization of the experiment. (**Left column pair**) In the absence of Tn/Tm, myosin freely translocates actin only limited by its rate of binding/unbinding. There is no difference in speed between WT and R256H ACTA1 filaments. (**Middle column pair**) In the presence of Tn/Tm and under low calcium conditions, both WT and R256H ACTA1 filament speeds are arrested. This means that R256H ACTA1 filaments are capable of regulation by Tn/Tm. (**Right column pair**) In the presence of Tn/Tm and high calcium conditions, WT ACTA1 filament speeds are restored, but R256H ACTA1 filament speeds remain inhibited. *N*=9 for all conditions representing average speed of all filaments in 3 focal fields quantified from 3 separate motility experiments. **B)** Regulated *in vitro* motility experiments at pCa 4 (high calcium) of filaments containing various concentrations of ACTA1 R256H. As concentration of ACTA1 R256H increases, the speed of the filament decreases. Note that 100% R256H data is reproduced from Figure 3A. 100% WT from Figure 3A is also reproduced but combined with additional replicates. *N*=18 for WT and 50% R256H while *N*=9 for all other conditions from 3 to 6 motility experiments. * = p < 0.05, ** = p < 0.01, *** = p < 0.001, and **** p < 0.0001 as calculated by one-way ANOVA followed by multiple comparisons of every condition with either Šidák correction (3A) or Tukey correction (3B) as recommended by GraphPad Prism. Normalized speed means min/max normalization across all conditions. **C)** pCa-speed response curves for 100% WT, 12.5% R256H ACTA1, and 25% R256H ACTA1 filaments and fit to the Hill equation. Each point represents *N*=6 from two motility experiments. Absolute speeds are shown to demonstrate differences in maximum speed which are calculated as WT 567 nm/s (95% CI 437 to 698 nm/s), 12.5% R256H 501 nm/s (95% CI 408 to 595 nm/s), 25% R256H 287 nm/s (95% CI 408 to 595 nm/s). These differences are better demonstrated in the experiment in Figure 3B. **D)** Calcium sensitivity is unchanged for mutant-containing filaments. WT, 12.5% R256H, and 25% R256H pCa_50_ are 5.302 (95% CI 5.118 to 5.485), 5.280 (95% 5.099 to 5.462), and 5.281 (95% CI 5.110 to 5.451). Cooperativity has similarly overlapping 95% CI: WT cooperativity is 5.053 (95% CI 1.241 to 8.865), 12.5% R256H cooperativity is 7.225 (95% CI -0.1912 to 14.64), and 25% R256H cooperativity is 12.18 (95% CI -7.143 to 31.51). Lines fit to the Hill equation have adjusted r^2^ of 0.89 for WT, 0.93 for 12.5% R256H, and 0.93 for 25% R256H. “Normalized*” indicates min-max normalization within each condition as opposed to across all data sets as for other normalized data in figure.

Next, we examined whether the myosin-based translocation of actin is affected by the presence of the regulatory proteins troponin (Tn) and tropomyosin (Tm) (henceforth, collectively referred to as Tn/Tm). Tm and Tn bind to the actin filament and they gate the calcium-dependent interactions between myosin and regulated thin filaments. We performed *in vitro* motility assays in the presence of cardiac Tn/Tm for both WT and R256H ACTA1 filaments. Both WT and R256H ACTA1 filaments were non-motile in the presence of Tn/Tm under low calcium conditions (pCa 9, **Figure 3A**, middle column pair) suggesting that Tn/Tm successfully inhibit actin-myosin interactions for both WT and mutant proteins at low calcium. Having confirmed Tn/Tm regulation of R256H ACTA1 at low calcium, we next wanted to test if the inhibition of the Tn/Tm complex could be relieved under high calcium (pCa 4) conditions. Consistent with previous studies, WT ACTA1 thin filaments showed robust movement at pCa 4 (**Figure 3A**, right column pair, black outlined column representing WT ACTA1). Surprisingly, we found that R256H ACTA1 filaments remained non-motile at pCa 4, and this speed was not different from pCa 9 (**Figure 3A**, right column pair, red outlined column representing R256H ACTA1). This is not due to an inability of myosin to interact with R256H actin, since these filaments move robustly in the absence of regulatory proteins (**Figure 3A**). Taken together, our results reveal that R256H thin filaments show inhibited motile activity, only in the presence of regulatory proteins. This result was unexpected, since a similar effect has not previously been observed for cardiomyopathy mutants.

All known patients with the R256H mutations are heterozygous for the mutation, and therefore express a mix of WT and mutant R256H. Moreover, ACTA1 protein is estimated to make up about 20% of the adult heart actin (ACTC1 is the major isoform in the heart) (31). As such, the mutant protein is likely expressed as a minority (∼10%) of actin in the heart along with a mixture of other actin isoforms. Therefore, we assessed the dose-dependence of the percent R256H ACTA1 on thin filament motility in the presence of Tn/Tm at pCa 4. To create filaments of various R256H ACTA1 concentrations, we mixed different concentrations of WT and R256H ACTA1 monomers which we subsequently polymerized (**Figure 3B**). We found that 12.5% R256H ACTA1 was sufficient to reduce the speed of actin filaments in the presence of Tn/Tm at pCa 4.

While we see a clear reduction in the speed of thin filament translocation with increasing amounts of mutant protein at fully-activating conditions (pCa 4), the heart generally works under sub-saturating conditions (pCa 5-7). Therefore, we measured the full speed-calcium relationship for several mixtures containing mutant protein. The relationship between speed and pCa can be fitted with the Hill equation to derive maximum speed, minimum speed, calcium concentration at 50% maximum speed (pCa_50_), and cooperativity of activation. Because even 50% R256H ACTA1 results in nearly all filaments being non-motile (**Figure 3B**), we generated speed-pCa curves using 25% and 12.5% R256H ACTA1 so there were enough motile filaments to quantify (**Figures 3C and 3D**). We found again that while the maximum speed under fully activating conditions decreased with increasing percentage of R256H ACTA1, the pCa_50_ was unaffected by increasing amounts of mutant protein suggesting that the calcium sensitivity of activation is not decreased. In summary, we discovered that a small percentage of R256H ACTA1 is sufficient to inhibit biochemical contractility only in the presence of troponin and tropomyosin.

### R256H ACTA1 filament has altered positioning of the tropomyosin-interacting residue K240

Given the unexpected magnitude of inhibition of myosin-driven motility of R256H filaments only in the presence of Tn/Tm, we hypothesized that a structural change in the R256H filament alters interactions with regulatory proteins. High resolution structures of the actin filament have revealed multiple charged residues in actin that specifically interact with tropomyosin (32–34). Given the changes in structure seen in the conformation of the actin monomer in the molecular dynamics simulations, we hypothesized that the mutation could affect residues in the actin filament involved in thin filament regulation.

To test this hypothesis, we used cryo-electron microscopy (cryo-EM) to determine the structures of both WT and R256H actin filaments bound to phalloidin. We used phalloidin to ensure stability of the mutant actin, since the mutant actin shows polymerization defects. It should be noted that phalloidin does not alter the natural conformations of the F-actin filament (35) . Thus, we proceeded with resolving our WT and R256H filaments with phalloidin. We achieved 3.4 Å resolution for the WT and 3.3 Å resolution for the R256H mutant. At this resolution we could directly visualize the position of amino acid side chains. We found that the WT and R256H ACTA1 electron density maps were nearly identical (**Figure 4A**, “Combined”). Our refined WT and R256H ACTA1 structures aligned well with the published PDB 6T1Y structure (35) with global RMSDs of 0.532 and 0.553 Å, respectively. Despite the similarity between WT and R256H ACTA1 structures, there was a significant difference between the WT and mutant in the positioning of K240 (**Figure 4B**). K240 (K238 by post-cleavage numbering) is located near R256, and it plays a key role in interacting with tropomyosin in the myosin-bound state (**Supplementary Figure 4**).

**Figure 4.**
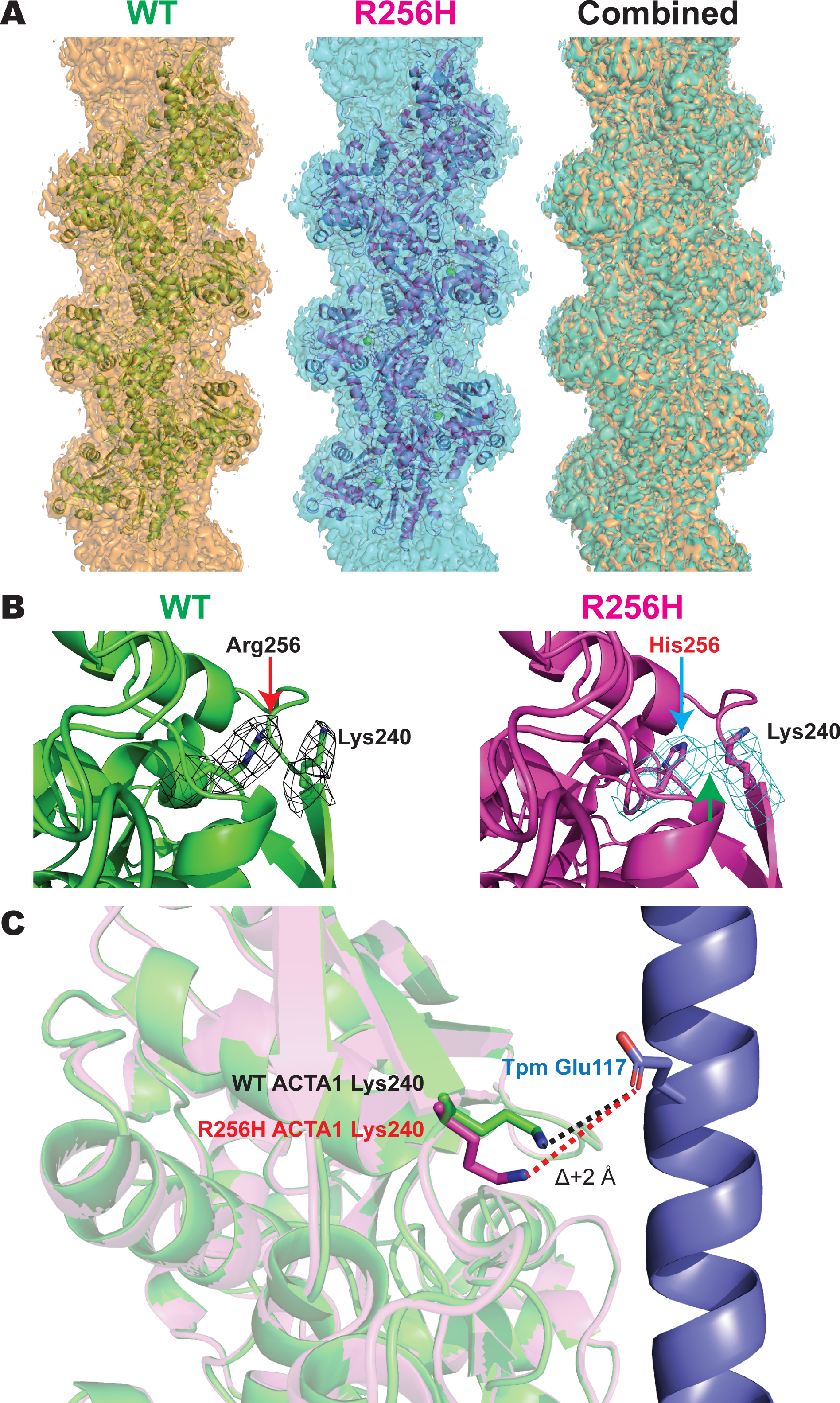
R256H ACTA1 alters position of ACTA1 K240 which may disrupt interaction with Tpm E117. **A)** WT and R256H ACTA1 have similar structures. Shown are cryo-EM densities at the 3σ contour level and superimposed atomic models rendered as cartoons for WT and R256H ACTA1. The cryo-EM densities of WT and R256H ACTA1 have substantial overlap as seen in the “Combined” figure. **B)** Focused view of WT and R256H ACTA1 256 residue and K240 residue which are rendered as sticks while the remainder of the peptide backbone is rendered as a cartoon. The wire mesh represents the 5σ contour level of electron density surrounding each respective residue. Note how R256H ACTA1 K240 moves closer in proximity to His256 such that there is overlapping electron density between the two residues (green arrow) which is not present in WT ACTA1. **C)** The change in Lys240 in R256H ACTA1 is predicted to cause an increase in distance between ACTA1 K240 and Tpm E117. Shown is the approximate position of tropomyosin E117 after aligning our atomic model to 8EFI which contains a high-resolution structure of F-actin bound by myosin in rigor in the presence of Tpm. We show only the Tpm strand that interacts with ACTA1 K240 for clarity.

We aligned our structures to a published structure of actin bound with tropomyosin and myosin (PDB 8EFI) (36). There are no changes near the myosin binding interface, consistent with the unchanged myosin *in vitro* motility (**Supplementary Figure 4B**); however, it is apparent that the change in position of K240 for the R256H filament results in K240 Nζ being about 2 Å further away from the ε-1 oxygen of E117 on tropomyosin which likely disrupts this interaction between actin and tropomyosin (**Figure 4C**). The potential loss of the interaction between ACTA1 K240 and tropomyosin E117 in the myosin-bound state of actin provides a potential link with the observed inhibition of myosin-induced R256H ACTA1 motility in the presence of tropomyosin.

### *ACTA1^R256H/+^* induced cardiomyocytes generate less force without disrupting calcium transients and demonstrate sarcomere disorganization

While our biochemical and structural studies demonstrate that R256H ACTA1 affects contractility at the molecular scale, it is not clear whether these changes translate to altered contractility in cardiomyocytes. Therefore, we generated isogenic human induced pluripotent stem cells (hPSCs) heterozygous for the ACTA1 R256H mutation using CRISPR/Cas9 (37). The parent wild type cells were shown by whole-exome sequencing to be free of any known genetic variants associated with familial cardiomyopathies (38). Two separate *ACTA1^R256H/+^* lines (referred to as “R256H clone 1” and “R256H clone 2” in supplementary figures) were created and independently compared to the parent wildtype isogenic line (referred to as “WT”). We differentiated our hPSCs to cardiomyocytes (hPSC-CMs) at least 28 days prior to performing experiments using well-established protocols (39, 40). Data were acquired from at least two separate differentiations.

Altered contractility is a hallmark of DCM, and therefore we investigated whether the R256H mutation is sufficient to alter contraction. To measure cardiomyocyte contractility, we used traction force microscopy (TFM) by plating hPSC-CMs onto 10 kPa hydrogels containing embedded fluorescent beads. Cells were adhered to micropatterned extracellular matrix in a 7:1 aspect ratio as we have done previously (38). The 10 kPa hydrogel mimics physiological cardiac stiffness, and the 7:1 aspect ratio has been shown to improve sarcomere alignment, and both factors improve maturation (41). We found that *ACTA1^R256H/+^* hPSC-CMs produced less force than WT cells, which we believe is the first time an ACTA1 mutation has been experimentally demonstrated to alter contractility in human cardiomyocytes (**Figures 5A and 5B**).

**Figure 5.**
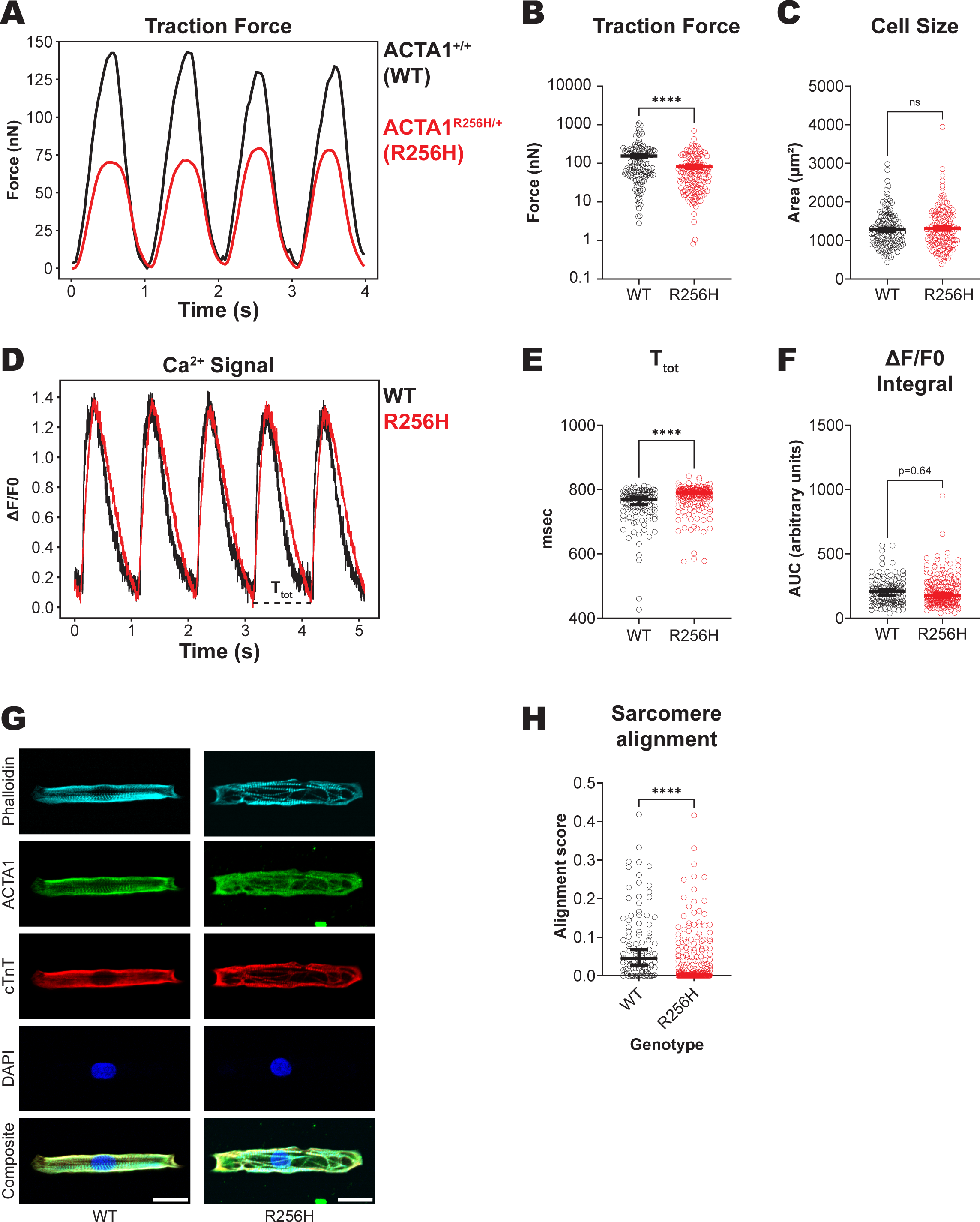
*ACTA1^R256H/+^* hPSC-CMs exhibit reduced force without a change in calcium flux but have disordered sarcomeres. **A)** Representative force tracing of a single WT and *ACTA1^R256H/+^* hPSC-CM (referred to as “R256H”) demonstrating the reduced force exhibited by mutant cells as determined by TFM. **B)** Summary of quantified force (WT mean force 155.2 nN vs R256H mean force 82.28 nN) and **C)** area (WT mean 1282 μm^2^ vs R256H mean area 1310 μm^2^) for individual hPSC-CMs. **** = p < 0.0001 as calculated by two-tailed non-parametric Mann-Whitney t-test. WT *N=*139 and R256H *N=*149 from at least 3 separate differentiations. **D)** Representative Ca^2+^ traces using FLUOFORTE from cells paced at 1 Hz. Dashed lines denote the time values quantified for **E)** the total time of calcium transient (T_tot_) with WT mean of 750.1 ms and R256H mean of 777.1 ms. **F)** Calcium flux (integral of ΔF/F0 signal) is not different. WT *N=*120 and R256H *N=*226 from at least 2 separate differentiations. **** = p < 0.0001 as calculated by non-parametric Mann-Whitney and parametric t-tests for “E” and “F” respectively. “G” p-value calculated by parametric t-test. Note that points for “E” and “G” are spread horizontally to try to minimize overlap. **G)** Examples of patterned hiPSC-CMs stained for different structures. ACTA1-specific antibody is validated in **Supplementary Figure 1C**. Phalloidin binds all polymerized actin (F-actin). Scale bar = 25 μm. **H)** Sarcomere alignment scores processed from phalloidin images using SotaTool. WT mean alignment score of 0.080 and R256H mean alignment score 0.032. A higher score corresponds to more alignment of Z-discs within the cell. WT *N=*99 and R256H *N=*258 from at least 2 separate differentiations. **** = p < 0.0001 calculated by non-parametric Mann-Whitney t-test. All error bars represent SEM though some are not visible due to the high density of points. Mutant cell data obtained from two separate clones, and the separated data can be found in **Supplementary Figure 5**.

While reduced contractility in the mutant cells is consistent with the observed molecular defects, there are multiple other downstream mechanisms that could also contribute the reduced contractility seen in the mutant cells, including alterations in the calcium transient, sarcomeric organization, cell size, and/or changes in the structure/function of the thin filaments. Therefore, we tested these possibilities. First, we manually traced cell edges, and we did not detect a change in size between *ACTA1^R256H/+^*and WT cells (**Figure 5C**). Next, we measured the calcium transients in mutant cells, since reduced calcium transients could contribute reduced contractility. We found that overall calcium transient duration was increased (**Figures 5D and 5E**), which would be expected to increase, not decrease contractility. However, the net calcium flux, calculated as the integral of the calcium transient from ΔF/F0, was not significantly different between the WT and the mutant, indicating the overall amount calcium moving into the cytoplasm is similar between both WT and R256H (**Figure 5F**). These data suggest that alterations in calcium handling do not explain the ACTA1 R256H cardiomyocyte hypocontractility phenotype. Finally, we used immunofluorescence to measure the alignment of sarcomeres in the cells, since alignment of the sarcomere is important for optimized force production, (**Figure 5G**). First, we identified successfully differentiated cells based on staining with cardiac troponin T (cTnT), a specific marker of cardiomyocytes (6). We also verified the presence and localization of ACTA1 to the sarcomere using an ACTA1 specific antibody (validated in **Supplementary Figure 1C**) and dual-staining with phalloidin which is specific for F-actin. We clearly observe the integration of ACTA1 in the sarcomere. Moreover, we observed that *ACTA1^R256H/+^* hPSC-CMs appear to have sarcomeric arrays that were less aligned with the longitudinal cell axis compared to WT. We quantified the alignment using SarcOmere Texture Analysis algorithm (SotaTool) which mathematically scores sarcomere alignment on a scale between 0 to 1, with higher alignment corresponding to a higher score (42). Consistent with our observations, WT cells had a significantly higher alignment score than *ACTA1^R256H/+^* hPSC-CMs (**Figure 5I**). Taken together, our results show that the mutant has reduced force production at the cellular level, consistent with our molecular phenotype. We did not detect changes in the calcium transient or cell size that could potentially contribute to this phenotype, but we did see changes in sarcomeric organization which could also contribute to decreased force.

## Discussion

Due to ACTA1’s high levels of expression in skeletal muscle, mutations in it have been shown to cause skeletal myopathies, and hundreds of missense mutations have been associated with disease. Recently, ACTA1 mutations were identified in patients with DCM; however, ACTA1 only makes up ∼20% of the actin in the heart (31), and it has been unclear whether or how these mutations impact cardiac contractility. Here, we performed the first multiscale study of a cardiomyopathy-associated ACTA1 mutation, R256H, which revealed fundamental connections between structure, molecular dysfunction, and altered cardiomyocyte contractility. We used molecular dynamics simulations to show that R256H mutant actin monomers favor a different average structure compared to the WT, and that the mutation likely affects nucleotide binding. To explore the functional implications of these predictions, we successfully purified recombinant WT and mutant ACTA1. We find that R256H ACTA1 has lower thermal stability and slower polymerization kinetics. Moreover, R256H potently disrupted actin-myosin translocation speeds only in the presence of actin regulatory proteins. We could observe the inhibitory effects of the mutant protein at likely physiological levels of mutant protein. We used cryo-EM to determine a high-resolution structure of R256H ACTA1, and we found an altered residue that likely disrupts an interaction with tropomyosin that is important for thin filament activation. Finally, we introduced the ACTA1 R256H mutation in hPSC-CMs and show that this disrupts force production and sarcomeric architecture. Taken together, our work demonstrates how an ACTA1 mutation can reduce cardiomyocyte contractility across multiple scales of organization, even at low levels of expression.

### Elucidating the primary driver of R256H ACTA1 hypocontractility

To date, several mutations in ACTA1 associated with skeletal myopathies have been characterized, and these mutations have shown a wide spectrum of defects in protein stability, polymerization, and actomyosin interactions (43–45). While the exact levels of ACTA1 mutant protein have not been measured in patient hearts, it is informative to do a back-of-the-envelope calculations. ACTA1 makes up ∼20% of the actin in the heart, and the patients were heterozygous for the mutation. Since R256H ACTA1 appears to cause DCM in an autosomal dominant manner (13), this suggests that ∼10% mutant protein in the heart is sufficient to cause disease. As such, mutations in ACTA1 associated with cardiomyopathies likely have large impacts on protein function, and we would expect to see impacts on function even at low expression levels. Consistent with our estimations, we found that R256H ACTA1 causes several dose-dependent changes in actin function at the molecular scale including changes in thin filament regulation and polymerization. It is also worth noting that ACTA1 expression increases in DCM, which likely increases R256H ACTA1 expression above 10% of total actin and establishes a feed forward loop of worsening cardiomyopathy (31).

Our data clearly show that molecular contractility is inhibited for R256H in the presence of the actin regulatory proteins troponin and tropomyosin, even at very low concentrations (∼10%) of mutant protein (**Figure 3**). This suggests that even small amounts of R256H expressed in the heart could affect contractility. To our knowledge, no other cardiomyopathy-associated thin filament mutation shows complete arrest of thin filament movement only in the presence of the Tn/Tm regulatory complex. Interestingly, other mutations in the same region of actin show a similar decrease in actin translocation speeds by myosin in the presence of tropomyosin. The D294V ACTA1 mutation, associated with severe congenital skeletal muscle fiber type disproportion, had a similarly dramatic effect of completely abrogating actin filament translocation with the *in vitro* motility assay regulated by tropomyosin (45). Additionally, the R258H ACTA2 (smooth muscle actin) mutation associated with thoracic aortic aneurysms also shows decreased actomyosin translocation speeds in the presence of tropomyosin (24). Taken together, we believe that the reduced speed of thin filaments by myosin in the presence of regulatory Tn/Tm is a primary driver of hypocontractility.

Interestingly, we only observe inhibition of motility in the presence of the regulatory proteins, suggesting that the mutation does not directly affect the kinetics of actin’s interaction with myosin. The regulation of actin and myosin binding by Tn/Tm has a well-established structural basis (33). Thus, we hypothesized that the R256H mutant would have an associated structural change in the actin filament that could affect regulation. Using cryo-EM, we solved a high-resolution structure and built atomic model of ACTA1 R256H (**Figure 4**). The R256H mutation causes K240 to move closer to H256, evidenced by an increase in electron density between these residues. We believe that the loss of the positively charged arginine when substituting with histidine allows K240 to move closer to residue 256. Importantly, it has previously been shown that K240 forms a specific interaction with the E117 residue on tropomyosin in the myosin-bound state (**Figure 4C**) (36). Thus, we speculate that the loss of this interaction decreases the favorability of the tropomyosin open state, potentially contributing to our observed decreases in contractility. Consistent with this notion, the E117K mutation in tropomyosin, causes congenital skeletal myopathy, and biochemical studies show that this mutation causes reduced thin filament activation associated with hypocontractility (46, 47).

In addition to altering thin filament regulation, our data clearly demonstrate that R256H can alter the stability of actin monomers and alter polymerization. Our molecular dynamics simulations predict that R256H could affect nucleotide binding (**Figure 1D**). Importantly, nucleotide binding has been shown to be critical for the stability of actin monomers and polymerization kinetics (18). Accordingly, R256H shows decreased thermal stability (**Figure 2A**). Unstable protein can either lead to decreased protein levels, indicating haploinsufficiency, or misfolding and aggregation which may contribute to pathogenicity (48). Haploinsufficiency of ACTA1 has not been shown to cause disease based on the lack of cardiac phenotype of *Acta1^+/^*^-^ mice (49). However, it should be noted that these mice do not have reduced *Acta1* transcripts, so it is possible that reducing *Acta1* at the mRNA and protein levels may still produce a haploinsufficiency phenotype that has yet to be uncovered. While unstable protein can lead to protein aggregation, potentially contributing to pathogenicity (48), we did not observe protein aggregates in our *ACTA1^R256H/+^* hPSC-CMs making this possibility less likely. Thus, it is difficult to directly connect decreased R256H stability to hypocontractility. Moreover, loss of nucleotide has been shown to dramatically decrease actin polymerization kinetics (18). Consistent with this notion, we find that R256H polymerization kinetics are severely reduced, especially its nucleation rate (Figure 2). However, R256H does not inhibit the polymerization kinetics of WT ACTA1 when mixed together (**Supplementary Figure 3**). Having ∼10% polymerization-defective actin seems unlikely to affect the overall thin filament length of the cardiomyocyte, though further work will be needed to test this hypothesis.

### hPSC-CMs with R256H ACTA1 are hypocontractile with disrupted sarcomeric organization

The hallmark of DCM is reduced ejection fraction, which is a hypocontractile state. We and others have shown that hPSC-CMs harboring a DCM-causing mutation in a sarcomere protein exhibit reduced force or stress production (38, 50–56). Consistent with this notion, *ACTA1^R256H/+^* hPSC-CMs show reduced force production without significant changes in calcium handling (**Figures 5A-F**). To our knowledge, this is the first direct evidence that ACTA1 mutations, which are usually associated with skeletal myopathies, can affect human cardiomyocyte contractility.

We and others previously found that DCM-associated mutations in hPSC-CMs cause increased cell size, but this was not the case for *ACTA1^R256H/+^*hPSC-CMs (38, 56). However, consistent with our studies with cardiac troponin T, we did find cells had sarcomere misalignment (**Figures 5G and 5H**). Defects in ACTA1 polymerization have been linked to skeletal myopathies and are thought to be partly driven by changes in thin filament length (57). Changes in thin filament length in cardiac tissues has also been linked to cardiomyopathy, which was seen with deletion of the cardiac-enriched actin nucleation factor, Leiomodin 2 (Lmod2) (58, 59). It is possible that the polymerization defects seen with R256H could contribute to the sarcomeric disarray seen here.

### ACTA1 mutations in cardiomyopathy and skeletal myopathy

Our data collectively shows biochemical and cellular evidence of cardiomyocyte hypocontractility in the presence of R256H ACTA1. Another important question is whether R256H ACTA1 impacts skeletal muscle function. Of the 3 patients with the R256H mutation, none had gross clinical features of skeletal myopathy (13). This was a surprising finding given the much higher expression of ACTA1 in skeletal muscle compared to the heart (6). The lack of a skeletal muscle phenotype is even more unexpected taken together with our *in vitro* motility data that shows 12.5% R256H ACTA1 is sufficient to disrupt actin-myosin translocation speeds in the presence of Tn/Tm (**Figure 3B**). In *ACTA1^R256H/+^* skeletal muscle, mutant actin could be as high as 50% which in our *in vitro* motility assay results in arrested filament movement.

We speculate about two possibilities how a skeletal muscle phenotype may be absent. Firstly, our data shows that R256H ACTA1 has profoundly reduced polymerization kinetics, especially its nucleation rate. It is possible that less R256H ACTA1 is incorporated into skeletal muscle than cardiac thin filaments due to different factors governing polymerization in skeletal and heart muscles. There are some thin filament proteins that are uniquely expressed in either skeletal or cardiac muscle. Leiomodins (LMODs) are important nucleation factors in striated muscle and exist in 3 isoforms: LMOD1-3 (60). LMOD2 is predominantly expressed in the heart and may allow for more incorporation of R256H ACTA1 into cardiac thin filaments compared to skeletal muscle thin filaments. Notably, a published crystal structure of LMOD2 with actin shows interactions of leiomodin with actin subdomain 2 (61), and our molecular dynamics simulations of the R256H G-actin shows changes in this region (**Figure 1C**). Similarly, nebulin plays a critical role in regulating skeletal muscle thin filaments and several ACTA1 residues in subdomain 4, where R256H is located, are predicted to form contacts with nebulin based on the recently resolved cryo-electron tomography (cryo-ET) structure of myofibrils with nebulin (62). Finally, cardiac and skeletal muscles express different tropomyosin isoforms in different abundance (63). Different tropomyosin isoforms affect actin polymerization differently providing a possible explanation to differential incorporation of WT and R256H ACTA1 in cardiac and skeletal muscles (64). Second, our data shows decreased thermal stability for R256H ACTA1. Alternatively, it is possible R256H ACTA1 has greater stability in cardiac muscle leading to more mutant protein in cardiac filaments. Regardless, future studies to probe more deeply into the differences in the cardiac and skeletal muscle phenotypes of R256H ACTA1, skeletal muscle models of R256H ACTA1 are needed.

There are several other mutations in ACTA1 that have been linked with DCM (9–12) with skeletal myopathy. However, there was a recent report of an ACTA1 G253S mutation that may, like R256H, cause DCM without a clinically apparent skeletal myopathy (14). We believe it will be important to apply the multiscale framework demonstrated here to further explore these ACTA1 mutations to see if these mutations have similarly potent effects on actin function like R256H.

### Conclusions

We define, for the first time, the likely mechanistic basis for the low-expression ACTA1 isoform causing cardiomyocyte hypocontractility. We demonstrate that the R256H ACTA1 mutation induces hypocontractility by potently disrupting actomyosin translocation in the presence of the thin filament regulatory proteins troponin and tropomyosin. Importantly, we show that ∼10% R256H ACTA1 protein is sufficient to cause molecular hypocontractility. We link our molecular hypocontractility to a structural change in R256H ACTA1 filaments that alters the interaction between ACTA1 K240 and tropomyosin E117. We find additional defects in R256H ACTA1 monomer stability and polymerization kinetics which may further contribute to pathology. We also show, for the first time, that an ACTA1 mutation can cause cardiomyocyte hypocontractility. Taken together, our work demonstrates that ACTA1 mutations have a role in cardiomyopathy and lays the foundation for future studies into ACTA1 mutations associated with cardiomyopathy.

## Materials and Methods

### Molecular dynamics

Simulations were performed similar to previous ACTC1 simulations (16). Details can be found in the supplemental methods.

### Recombinant ACTA1 purification

We adapted a SF9/baculovirus system that has been successfully used to produce other muscle actins including mutant actins (19–22). Details can be found in the supplemental methods.

### Thermal shift assay

All thermal shift experiments were performed using the Protein Thermal Shift Dye Kit (Thermo Fisher Scientific) and following the standard commercial protocol with an Applied Biosystems QuantStudio 3 Real-Time PCR system. Protein was diluted to a final concentration of 2 µM prior to performing melt curves. Custom Python scripts were written to derive T_m_ from raw data. All data and analysis scripts will be available for download at the time of publication.

### Pyrene-actin polymerization fluorescence assay

Rabbit ACTA1 was pyrene labeled, and experiments were performed as previously described. (65, 66). Details can be found in the supplemental methods.

### TIRF analysis of fluorescent labeled ACTA1

A subset of rabbit ACTA1 was Alexa Fluor 647 labeled and another subset of rabbit ACTA1 was biotinylated as previously described (66). Details can be found in the supplemental methods.

### *In vitro* motility assay

Porcine cardiac myosin and actin purification, and preparation of recombinant human cardiac tropomyosin and troponin were performed as we have described previously (38). Details can be found in the supplemental methods.

### Cryo-EM

We utilized methods similar to prior (67–69). Details can be found in the supplemental methods.

### Structure alignment and interpolation for interaction with tropomyosin

We aligned our final WT and R256H ACTA1 models to the published PDB 8EFI model of human cardiac actin-tropomyosin-myosin complex in rigor form with PyMol. We calculated interatomic distances using built-in tools in PyMol.

### hPSC line derivation, maintenance, and hPSC-CM differentiation

*ACTA1^R256H/+^* hPSCs were derived by the Genome Engineering and iPSC Center (GEiC) at Washington University in St. Louis, cultured, and differentiated as detailed before (38). Details can be found in the supplemental methods.

### Traction force microscopy

Traction force microscopy was performed as previously described (38, 56, 70). Details can be found in the supplemental methods.

### Measurement and analysis of calcium transients in live cells

We performed calcium transient imaging as previously except with a different calcium indicator dye and used a different method for analysis with CalTrrak (70, 71). Details can be found in the supplemental methods.

### Immunofluorescence staining and measurement of sarcomere alignment

Immunostaining was performed similar to prior (38), but SotaTool was used to calculate sarcomere alignment (42). Details can be found in the supplemental methods.

## Acknowledgments

We thank Lina Greenberg for her technical help with hPSCs and reagent preparation for the *in vitro* motility assay. We thank Teresa Naismith and DeHaven McCray for their respective help with actin purification and TIRF microscopy fluorescent actin polymerization experiments. We thank Patricia Fagnant and Kathleen Trybus for discussion regarding recombinant actin purification. We thank Katherine Basore and Brock Summers at the Washington University Center for Cellular Imaging (WUCCI) for preparing the cryo-EM samples and collecting the data which was funded through the Department of Biochemistry and Molecular Biophysics Seed Grant (to A.G., R.Z., and M.J.G.). We would like to acknowledge the Genome Engineering and iPSC Center (GEiC) at Washington University in St. Louis for creation of our hPSC line which was partly funded by the Washington University School of Medicine in St. Louis Institute of Clinical and Translational Science Just-In-Time Core Usage Funding Program (JIT744 to A.G. and M.J.G.). Finally, we thank Anjali Owens for clinical discussions regarding the ACTA1 R256H mutation. This work was supported by the National Institutes of Health (T32 HL007081 to A.G.; R01 GM138448 to S.J.; R01 GM138854 to R.Z.; R01 HL141086 to M.J.G.; R01 HL138466, R01HL139714, R01 HL151078, and R35 HL161185 to K.J.L); the Leducq Foundation Network (20CVD02 to K.J.L.); the Burroughs Wellcome Fund (1014782 to K.J.L.); the Children’s Discovery Institute of Washington University and St. Louis Children’s Hospital (CH-II-2015-462, CH-II-2017-628 to K.J.L.; PM-LI-2019-829 to M.J.G and K.J.L.); the WUCCI (CDI-CORE-2015-505 to M.J.G.).; the Foundation of Barnes-Jewish Hospital (8038-88 to K.J.L.)

## Notes

### Competing Interest Statement

AG reports no disclosures. SJ reports no disclosures. RZ reports no disclosures. KJL reports consulting agreements from Implicit Biosciences, Cytokinetics, Medtronic, Kiniksa Pharmaceuticals, and SUN Pharmaceuticals. MJG reports no disclosures.

