## Supplemental Materials for "Dilated cardiomyopathy-associated skeletal muscle actin (ACTA1) mutation R256H disrupts actin structure and function and causes cardiomyocyte hypocontractility"

**This PDF file includes:**

SI Methods

Supplementary Figures 1-5

SI References

**Other supporting materials for this manuscript include the following:**

Datasets S1 to S2

SI Methods

**Molecular dynamics.** Simulations were performed similar to previous ACTC1 simulations (1). Specifically, we performed simulations with NAMD 3.0 alpha (2) and the CHARMM36 forcefield (3-20). NAMD was developed by the Theoretical and Computational Biophysics Group in the Beckman Institute for Advanced Science and Technology at the University of Illinois at Urbana-Champaign. All simulations and analyses were performed on either a Dell Precision 5820 Tower with Intel Xeon W-2445 8 core/16 thread CPU, NVIDIA RTX A4000 16 GB GPU, DDR4 64 GB RAM or a custom built AMD Ryzen 9 5950X 16 core/32 thread CPU, EVGA NVIDIA GeForce RTX 3090Ti 24 GB GPU, DDR4 128 GB RAM running the Ubuntu operating system. For all simulations, we used the monomeric structure of ACTA1 bound to ADP and calcium obtained by X-ray crystallography, PDB 1J6Z (21). This structure was chosen for its high resolution, completeness, and the lack of use of actin complexing reagents like gelsolin. The structure contained no intrastrand gaps, however residues 3-5 (“DED”) at the N-terminus and 375-377 (“KCF”) at the C-terminus were not resolved in the available structure and homology modeling was not utilized to fill in these disordered regions. ATP bound ACTA1 structures like PDB 3HBT (22), had missing residues creating two non-contiguous peptide chains, or used a hydrolyzing-resistant ATP analog, phosphoaminophosphonic acid-adenylate ester, like PDB 1NWK (23) which could bias our simulation in unknown ways. We generated protein structural files compatible with the CHARMM36 forcefield using CHARMM-GUI (24-27). We also used this interface to introduce the R256H mutation in the HSD, HSE, and HSP configurations. Each structure was fully solvated with the TIP3P water model (28) and autoionized to balance charge using VMD (29). Solutes used in autoionization were given a minimum distance of 15 Å to the edge of the water box. All simulations used periodic boundary conditions and used Langevin dynamics and Langevin piston method to maintain a constant temperature of 310K and pressure of 1 atm. We used a cutoff of 12 Å for van der Waals interactions and short-range electrostatics while the particle-mesh Ewald method was used for long-range electrostatics (30). We performed a 500 picosecond energy minimization and equilibration with a fixed peptide backbone. From this structure, we performed 5 separate 1 μs simulations. Data were analyzed using the MDAnalysis Python package (31, 32) with additional algorithms (33, 34). PyMOL was used to generate images (35-37). All data files with scripts used for analysis will be available for download at the time of publication.

**Recombinant ACTA1 purification.** We adapted a SF9/baculovirus system that has been successfully used to produce other muscle actins including mutant actins (38-41). We did not pursue a bacterial expression system since it has been shown that they cannot properly fold actin due to the absence of a eukaryotic chaperone (42). We briefly tested the *Pichia pastoris* yeast system which contains a stably integrated actin-specific N-acetyltransferase Naa80 and SETD3 methylase (which methylates His73) and has been shown to yield large quantities of non-muscle actin (ACTB1, ACTG1, ACTG2) (43). Our initial attempts at using the *Pichia pastoris* system yielded insoluble ACTA1, so we did not pursue this system further. Therefore, we continued with our SF9/baculovirus system in close resemblance to a prior publication studying ACTA2 (44).

We cloned mouse Acta1 (NP_001258970.1), which is 100% identical to human ACTA1 protein, from mouse heart cDNA and into the pFastBac1 (part of the Bac-to-Bac system by Thermo Fisher Scientific) with a flexible C-terminal linker encoding ASRGGSGGSSGGSSA containing a chymotrypsin cleavage site predicted to cut flush with the terminal phenylalanine of the ACTA1 protein. Next, we sequentially added to the C-terminus human thymosin-ß4 (NP_066932.1) and an 8X histidine tag using synthetic gBlocks from Integrated DNA Technologies (IDT). ACTC1 (NP_001393411.1; only used as a control in Supplementary Figure 1) with the same C-terminal tag was created synthetically with codon optimization and using gBlocks from IDT. The ACTA1 R256H mutation was created using the QuickChange Lightning kit from Agilent. All inserts were confirmed by sequencing. Bacmid DNA and baculovirus were created per original manufacturer protocol for the Bac-to-Bac system by Thermo Fisher Scientific.

For protein expression, 250 mL of SF9 4-10 x 10^6^ cells/mL infected with P1 (amplified only once) virus was harvested within 72 hours of infection with a goal viability of > 70%. Cells were pelleted at 5,000xg for 15 minutes at 4°C. The supernatant was discarded, and the pellet was flash frozen in liquid nitrogen to initiate cell lysis. Frozen cells were stores at -80°C until use. On the day of use, pellets were allowed to thaw at room temperature and then stored on ice. Next, lysis was performed using “Lysis Buffer” containing 20 mM Tris-HCl pH 7.5, 300 mM NaCl, 10 mM Imidazole, 2 mM MgCl_2_, 1 mM ATP, 1 mM DTT, 0.5% IGEPAL, 1mM PMSF, 0.01 mg/mL aprotinin, and 0.01 mg/mL leupeptin. All protease inhibitors were added within 30 minutes of mechanical lysis. For mechanical lysis, cell pellets were subjected to dounce homogenization followed by sonication on ice. Cell lysate was clarified by spinning at 35,000xg in an SS-34 rotor for 1 hour at 4°C. The supernatant was applied to a 1 mL column volume of HisPur (Thermo Fisher Scientific) equilibrated with “Wash Buffer” containing 20 mM Tris-HCl pH 7.5, 300 mM NaCl, 20 mM Imidazole, 5 mM CaCl_2_, 0.15 mM ATP, and 1 mM DTT. After passing over the column once, flow through was applied again to the column. The column was then washed with 45 column volumes of “Wash Buffer” followed by 25 column volumes of “G-actin” buffer containing 20 mM Tris-HCl pH 8, 0.2 mM CaCl_2_, 0.2 mM ATP, and 0.5 mM DTT. Then, the column was capped and 6 mL of 10 µg/mL chymotrypsin from Sigma-Aldrich in “G-actin” buffer was added to the column which was rocked overnight at 4°C. The following day, the original 6 mL of eluate was collected followed by more elution by an additional 6 mL of “G-actin” buffer. Chymotrypsin was quenched by adding PMSF to a final concentration of 1 mM. The 12 mL of eluate was spin concentrated approximately 24-fold using an Amicon Ultra-15 Centrifugal Filter with a 3 kDa molecular weight cut off. Next, G-actin polymerization was induced by spiking in EGTA to an approximate concentration of 5 mM to chelate Ca^2+^ with simultaneous addition of MgCl_2_ to an approximate concentration of 2 mM followed by adding KCl to an approximate concentration of 100 mM and ATP to an approximate concentration of 0.5 mM. Polymerizing G-actin was left overnight at 4°C. The following day, F-actin was pelleted by centrifugation at 516,000xg for 1 hour at 4°C in a Beckman Coulter TLA 120.2 rotor. Supernatant was discarded and the pellets were resuspended in G-actin buffer (except with 10 mM of DTT instead of 1 mM). The resuspension was then dialyzed in G-actin buffer, to depolymerize the actin, 4 times over 3 days using a Slide-A-Lyzer dialysis cassette (Thermo Fisher Scientific) with a 10 kDa molecular weight cut-off. The final G-actin buffer was spun at 516,000xg for 30 minutes at 4°C in a Beckman Coulter TLA 120.2 rotor to remove any insoluble material. All experiments were done with fresh protein used within 3 weeks of purification. Protein concentration was calculated by measuring absorbance at 290 nm and using the 38.5 µM/OD_290_ conversion.

**Pyrene-actin polymerization fluorescence assay.** Rabbit ACTA1 was pyrene labeled and experiments were performed as previously described. (45, 46). Briefly, actin oligomers were removed by spinning at 90,000xg for 1 h at 4°C and taking only the top 80% of the supernatant immediately after spinning. Each reaction was performed with 5% pyrene-labeled rabbit ACTA1, recombinant WT ACTA1, and recombinant R256H ACTA1 which were mixed in G-actin buffer prior to initiation of polymerization. Initiation was done by adding 3 µL of 20X initiation mix (200 mM imidazole pH 7.0, 1 M KCl, 20 mM MgCl_2_, and 20 mM EGTA) in 60 µL reactions. Polymerization was measured over time using 364 nm excitation and 416 nm emission in an Edinburgh Instruments FS5 fluorometer at 25°C. Data were processed and analyzed using custom written Python scripts. Elongation rate was calculated by performing simple linear regression at half maximal intensity. All data and analysis scripts will be available for download at the time of publication.

**TIRF analysis of fluorescent labeled ACTA1.** A subset of rabbit ACTA1 was Alexa Fluor 647-labeled and another subset of rabbit ACTA1 was biotinylated as previously described (46). Similar to above, actin polymers were removed by spinning at 90,000xg for 1 h at 4°C and taking only the top 80% of the supernatant immediately after spinning. Slides were cleaned and prepared as previously described. Each flow chamber was prepared immediately before each reaction like prior: 3 min incubation in 1% HBSA (1% BSA in 10 mM Imidazole pH 7.4, 50 mM KCl), 30 s incubation in 4 mg/mL streptavidin, washed with 1% HBSA, and then equilibrated with TIRF buffer (20 mM Imidazole pH 7.4, 100 mM KCl, 0.4 mM ATP pH 7.0, 2 mM MgCl_2_, 2 mM EGTA, 20 mM DTT, 30 mM glucose, 0.5% methylcellulose 4000 cP). To initiate polymerization, actin was rapidly mixed to a final concentration of 2 µM with 10% Alexa Fluor 647-labeled and 0.25% biotinylated rabbit ACTA1 in TIRF buffer supplemented with 2 mg/mL catalase and 10 mg/mL glucose oxidase. The point of mixing was designated as the start time to the reaction. The polymerizing mixture was transferred to a flow chamber, and then imaged with a Nikon Ti2 inverted microscope equipped with through-the-objective TIRF illumination, a Nikon LU-4N 4-laser unit, and an Andor Technology iXon Ultra 888 EMCCD camera. Analysis was performed using the Fiji plugin “FilamentDetector” (47). Notable parameters include a “line width” setting of 6 to 7, a “high contrast” setting of 400 for WT and 700 for R256H reactions, “low contrast” setting of 1, and a “simplify tolerance distance” of 2 for WT and 10 for R256H reactions. Settings were varied based on manual inspection of filament tracking to minimize background noise while maximizing filament detection which varied from flow cell to flow cell. Data was exported and then analyzed using custom written Python scripts. Filament detection became erroneous once filaments started to overlap which was determined by a drop in the average filament per focal field. Data were truncated past this point, and then the number of filaments per frame were quantified. Nucleation rate determined by simple linear regression. All data and analysis scripts will be available for download at the time of publication.

***In vitro*** **motility assay.** Porcine cardiac myosin and actin purification, and preparation of recombinant human cardiac tropomyosin and troponin were performed as we have described previously (48). The *in vitro* motility assay was also similarly executed, but in brief: phalloidin-stabilized porcine cardiac F-actin and rhodamine-phalloidin-stabilized F-actin composed of recombinant human WT ACTA1 and R256H ACTA1 were prepared in KMg25 buffer (25 mM KCl, 2 mM EGTA, 60 mM MOPS, 1 mM DTT, and 4 mM MgCl_2_). Enzymatically inactive myosin was removed by ultracentrifugation of full-length porcine cardiac myosin in high salt buffer (KMg25 with approximately 300 mM KCl) with phalloidin-stabilized porcine cardiac actin. Nitrocellulose coated flow cells were loaded with one volume (35 µL) of 200 mg/mL myosin, one volume 1 mg/mL BSA, 1 volume 1 µM phalloidin-stabilized porcine cardiac F-actin, 4 volumes KMg25 + 1 mM MgATP, 2 volumes KMg25, 1 volume 40 nM rhodamine-phalloidin-stabilized F-actin composed of recombinant ACTA1, 1 volume 0.2 µM recombinant human cardiac tropomyosin and troponin complex in KMg25, and, finally, 1 volume activation buffer (25 mM KCl, 2 mM EGTA, 60 mM MOPS, 5 mM MgATP, 1 mg/mL glucose, 192 U/mL glucose oxidase, 48 µg/mL catalase, 0.2 µM recombinant human cardiac tropomyosin, and troponin complex in 0.5% methyl cellulose) at various pCa concentrations as previously described (49). Fluorescent thin filament translocation was recorded as 30 s videos at 30°C (heated above ambient temperature to minimize thermal fluctuations as microscope room warmed throughout the day) using a Bioptechs objective heater. Images were auto-balanced in Fiji (47), and then analyzed using FASTrack (50) using the following parameters: velocity averaged over 5 frames, minimum path length of 5 pixels, no filtering of fluctuating velocities, and maximum allowed distance of 2664.5 nm between each frame. Note that FASTrack detected very low speed even when filaments were non-motile. Thus, when calculating “absolute speeds” in Figure 3C, the average pCa 9 speed was subtracted from the total speed. Fitting to Hill plots was done with MATLAB as previously described (48). All data and analysis scripts will be available for download at the time of publication.

**Cryo-EM sample preparation and data processing**. WT and R256H ACTA1 were polymerized by mixing recombinant G-actin (initially in G-actin buffer [20 mM Tris-HCl pH 8, 0.2 mM CaCl_2_, 0.2 mM ATP]) to a final concentration of 0.25 mg/mL in KMg25 buffer with 10 mM DTT (25 mM KCl, 2 mM EGTA, 60 mM MOPS, and 4 mM MgCl_2_) and 12 µM phalloidin. Filaments were allowed to polymerize overnight at 4°C and kept at 4°C for remaining steps. Cryo-EM grids were prepared using a Vitrobot Mark IV (Thermo Fisher Scientific) with a chamber temperature of 4°C and humidity of 95%. Three mL of 0.15 mg/mL WT ACTA1 or 0.1 mg/mL R256H ACTA1 filaments were pipetted onto lacey carbon grids (Ted Pella), which had been freshly glow-discharged for 15 s with a GloQube glow-discharger. The grids were blotted for 2 s after a 20 s wait time, then plunge frozen into liquid ethane and stored under liquid nitrogen.

The samples were imaged using a Glacios 200kV microscope equipped with a Falcon 4 direct electron detector (ThermoFisher Scientific). Movies were recorded with a defocus range of -0.8 to -2.4 µm and at a nominal magnification of 120kx, corresponding to a calibrated pixel size of 1.184 Å. Each movie consisted of 50 frames over an 11.63 s exposure time and the total dose of the WT and R236H ACTA11 samples were 56.7 and 55.3 e/Å^2^, respectively. A total of 2,709 movies for the WT ACTA1 and 2,593 movies for R256H ACTA1 were recorded. We processed our data with single-particle methods similar to prior (51, 52). Briefly, we used CryoSPARC (53) for drift correction and electron-dose weighting (53). Actin filaments were automatically picked from the micrographs using template based 'Filament tracer' in CryoSPARC. The templates were a few 2D class averages from a small subset of actin filaments that were picked manually. Then the filaments were computationally cut into overlapping boxes (400x400 pixels) with an arbitrary 70 Å step size between adjacent boxes. Two-dimensional classification was used to remove junk particles and off-centered particles. The good particles were used to create the initial model using 'Ab-Initio Reconstruction' in CryoSPARC. In the next step, we used 'Helical Refinement' in CryoSPARC to obtain a 3D reconstruction, using the known helical symmetry of actin filaments (rise 28 Å, twist -168°) as the starting point. Next, we performed 'Local CTF refinement' and another round of 'Helical Refinement', resulting in a cryo-EM structure of actin filament at 3.4 Å resolution for the WT and 3.3 Å for the R256H mutant.

**Cryo-EM model building and refinement.** We utilized Coot v0.9.8.92 for all of our model building (54) and PHENIX v1.20.1-4487-000 for refinement (55) similar to prior (51). We used the PDB 6T1Y cryo-EM structure of phalloidin-stabilized F-actin as our starting structure for both WT and R256H (56). Initial model building and refinement was done on WT ACTA1 which was then used for R256H ACTA1. We initially fit a single actin subunit from the 6T1Y model into our maps using Chimera (57). We then performed real-space refinement of this single monomer with torsion, planar peptide, trans peptide, and Ramachandran restraints with an auto-calculated weight of 60. Next, we refined our single actin molecule with a resolution limit of 3.4 Å through several rounds which included interim manual refinement in Coot. We performed an interim validation with the built-in MolProbity module in PHENIX (58).

Following refinement of the monomer, we applied helical symmetry to add five additional actin subunits for a total of six subunits. Then, we performed additional refinement with PHENIX similar to above with additional manual refinement in Coot to ensure proper alignment of each subunit to each other. Finally, we restored ADP, Mg^2+^ ions, and phalloidin molecules in their original configuration as the 6T1Y starting structure which already had excellent fit of in the WT and R256H ACTA1 maps. We performed a final validation with MolProbity. PDB validation reports are attached as separate supplemental files. Images for our model and electron density maps were created in PyMol.

**hPSC line derivation, maintenance, and hPSC-CM differentiation.** *ACTA1^R256H/+^* hPSCs were derived by the Genome Engineering and iPSC Center (GEiC) at Washington University in St. Louis, cultured, and differentiated as detailed before (48). In brief, the parent human BJ fibroblast cell line (CRL-2522 American Type Culture Collection) was reprogrammed to hPSCs by GEiC which was previously validated by whole exome sequencing (48).Using CRISPR-Cas9, we cut our target site using the gRNA sequence of 5’-CAA CGA GCG CTT CCG CTG CCC GG-3’ chosen to minimize off target effects. Targeting was performed with a ssODN with the sequence of 5’- TCC TCC CTG GAA AAG AGC TAC GAG CTG CCA GAC GGG CAG GTC ATC ACC ATC GGC AAC GAG CAC TTT CGC TGC CCG GAG ACG CTC TTC CAG CCC TCC TTC ATC GGT GAG CCC CGC TCG CCC TCG CCC CGG C-3’ where the “A” denotes a G>A mutation to induce R256H, and the “T” denotes a silent mutation to block cutting of the ssODN by the gRNA. The presence of the heterozygous mutation was verified by next generation sequencing and verified to be mycoplasma free by the GEiC. Two separate lines were independently maintained and differentiated.

For culture, cells were maintained with feeder-free culture using Gibco StemFlex media. Cells were differentiated with small-molecule modifiers of WNT signaling (59, 60), enriched with metabolic selection (61), and then replated after > 28 days of differentiation for traction force microscopy and after > 35 days for calcium imaging.

**Traction force microscopy.** Traction force microscopy was performed as previously described (48, 62, 63). Briefly, hPSC-CMs were singularized and passaged on to Geltrex (Thermo Scientific) coated patterns on a 10 kPa hydrogel containing fluorescent beads. Cells were paced at 1 Hz. To minimize confounding force differences due to issues with electrical-calcium coupling, only cells that responded to pacing at 1 Hz were used for analysis. A spinning disk confocal microscope was used to record beating cells and fluorescent bead movement. These videos were inputted into the Contrax software (64, 65) which outputs the data displayed in our figure. Cell area was calculated within this software by manually tracing cell edges. All data and analysis scripts will be available for download at the time of publication.

**Measurement and analysis of calcium transients in live cells.** We performed calcium transient imaging as previously except with a different calcium indicator dye and used a different method for analysis with CalTrack (62, 66). Briefly, hPSC-CMs were singularized and seeded onto Geltrex-coated patterns on glass bottom plates (MatTek Life Sciences) and allowed to recover for 5-7 days. On the day of imaging, cells were loaded with 2.5 µM FLUOFORTE (Enzo Life Sciences) in RPMI-B27 with insulin for 20 minutes followed by washing twice in Tyrode’s solution (1.8 mM CaCl_2_, 135 mM NaCl, 4 mM KCl, 1 mM MgCl_2_, 5 mM glucose, and 10 mM HEPES, pH 7) and allowed to de-esterify for 15 minutes at 37°C. Cells were recorded in line scan mode using a 40X objective using 488 nm excitation and 515 nm emission collection for less than 35 minutes since cells began exhibiting toxicity past this point. With line scans, we recorded signal intensity every 1.9 ms (approximately 526 frames per second) for 20 seconds. Raw image files were opened in Fiji and processed into time-dependent pixel intensity using a custom Fiji macro. Next, a custom Python script was written to convert pixel intensity files into a format compatible with the CalTrack calcium transient analysis program and relevant data is shown (66). ΔF/F0 signal was integrated using a custom Python script. Statistical analysis was performed with GraphPad Prism. All data and analysis scripts will be available for download at the time of publication.

**Immunofluorescence staining and measurement of sarcomere alignment.** Immunostaining was performed similar to prior (48). SotaTool was used to calculate sarcomere alignment (67). Cells were seeded in the same manner as above, but after 5-7 days cells were fixed in 4% paraformaldehyde/phosphate buffered saline (PBS) followed by washing with PBS and storage at 4°C until immunostaining. On the day of immunostaining, cells were incubated in 0.1% Triton X-100 in PBS at room temperature for 15 minutes, washed in PBS, and then blocked in 5% donkey serum (Sigma). Cells were then incubated with primary antibody at 4°C. Primary antibodies include 1:400 rabbit anti-cardiac-TnT (ab45932 Abcam) and 1:100 mouse anti-ACTA1 (MUB0108P Nordic MUBio) in blocking solution. The following day, cells were washed thoroughly in PBS followed by 1:150 secondary antibodies and 1:1000 Alexa Fluor 647 phalloidin (Invitrogen). Following incubation in the dark at room temperature, nuclei were stained with 0.3 nM 4′,6-diamidino-2-phenylindole (DAPI) followed by thorough washing. Images were acquired with a Nikon A1Rsi confocal microscope with an Andor Zyla 4.2 Megapixel sCMOS camera as Z-stacks. Images were processed with Fiji which included extracting the channel with phalloidin staining, and then performing a maximum intensity projection. Alignment scores were calculated from the maximum intensity projections using SotaTool using default parameters. Negative alignment scores and failed alignments were given a score of “0”. Data were tabulated and statistical analysis was performed with GraphPad Prism. All data and analysis scripts will be available for download at the time of publication.

Supplemental Figures

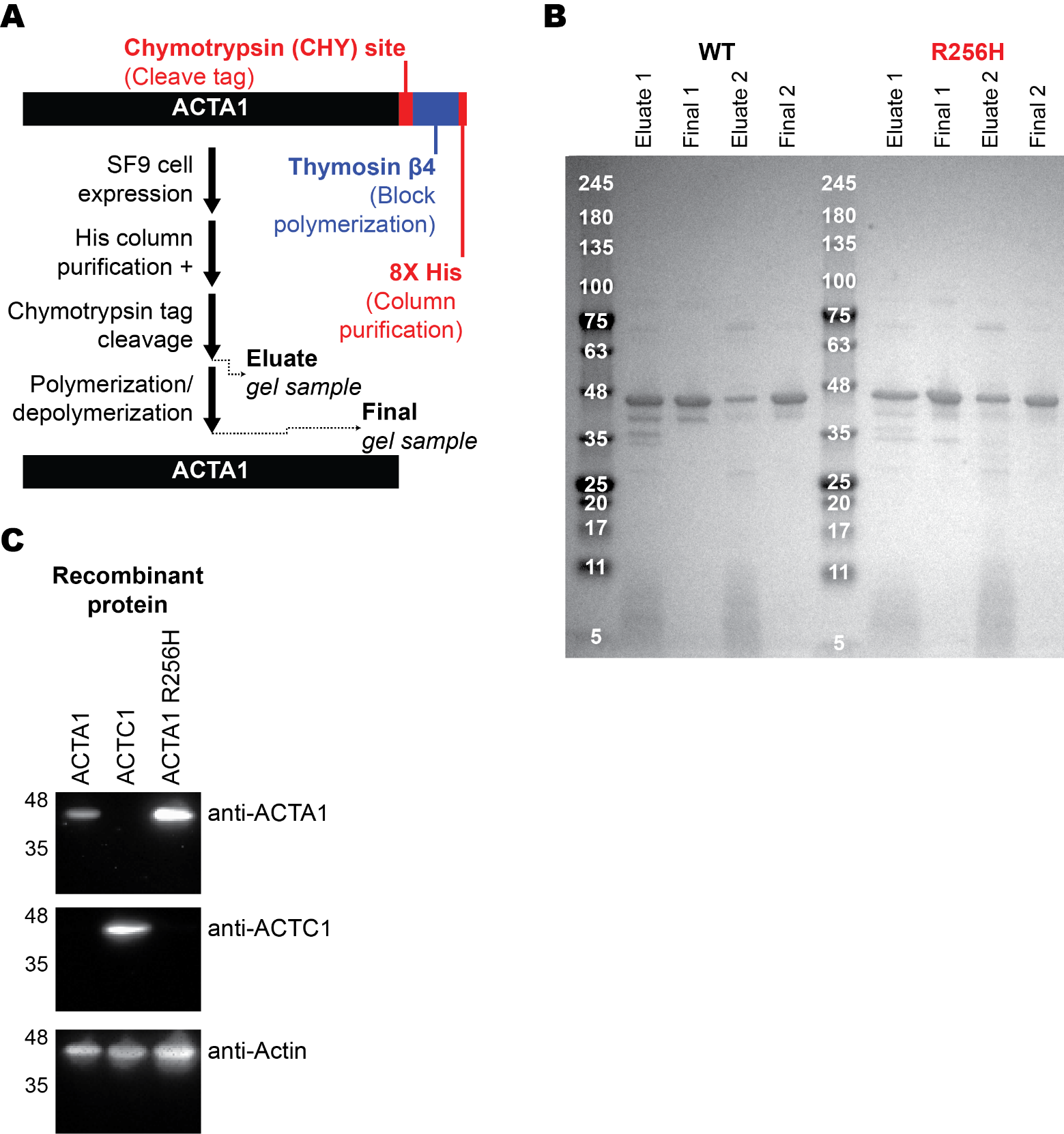

Supplementary Figure 1. Recombinant ACTA1 purification strategy. A) Overall purification strategy. ACTA1 is C-terminally fused to an epitope containing thymosin β4, which inhibits polymerization with endogenous SF9 actin, and an 8X His tag that allows for binding to the Ni^2+^ column. Once the protein is bound to the column, it is removed by chymotrypsin cleavage of the C-terminal tag which leaves no additional amino acid residues. Also noted are the two protein samples displayed in B) representative SDS-PAGE of protein at the indicated purification steps from two separate protein purifications. Initial eluate from the column has a dominant band of recombinant protein, but multiple other bands were present. We found that a single round of polymerization/depolymerization improved purity. C) Verification of recombinant protein by Western blot using ACTA1 specific antibodies. This figure also serves to validate our commercially purchased ACTA1 (Nordic MUBio MUB0108P) and ACTC1 (Nordic MUBio MUB0109P) antibodies. Note that the ACTC1 control protein was created recombinantly using the same protocol described here. Anti-actin antibody is the A2172 monoclonal anti-sarcomeric actin antibody from Sigma.

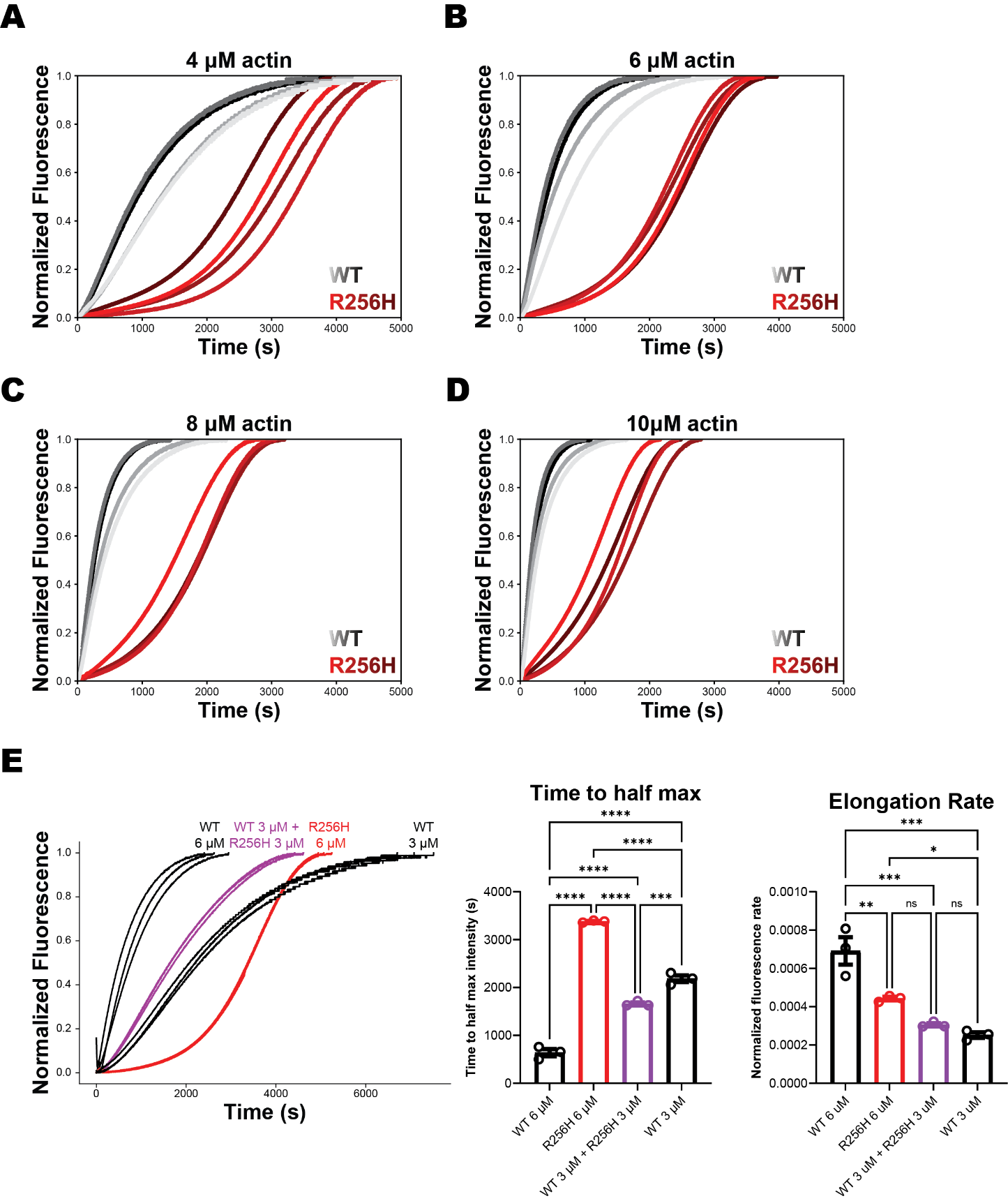

Supplementary Figure 2. A-D) Individual pyrene polymerization traces for ACTA1 WT and R256H used to generate the data in Figure 2C and 2D. WT traces are shown in shades of black while R256H traces are shown in shades of red. *N* = 4 for each condition and represent two polymerization experiments from two independent protein preparations. E) ACTA1 R256H does not inhibit ACTA1 WT polymerization. (Left) Individual pyrene polymerization traces with indicated WT and R256H mixtures. Each condition contains three polymerization replicates (*N*=3) from a single protein preparation. Note that the 6 μM data is used in Figure 2C. (Right) Quantification of kinetic parameters in the same manner as Figure 2C. Note that the time to half maximum intensity is significantly less for the mixed WT 3 μM + R256H 3 μM compared to WT 3 μM alone indicating that the mixed condition has faster kinetics than the WT only condition. The elongation rate is not significantly different for these two conditions. If R256H had a dominant inhibitory effect on polymerization, the WT + R256H mixed condition would be expected to have slower kinetics with a longer time to half maximum intensity and a lower elongation rate at the equivalent concentration of WT monomer which is not the case. * = p < 0.05, ** = p < 0.01, *** = p < 0.001, **** = p < 0.0001 as calculated by one-way ANOVA followed by Tukey multiple comparison correction.

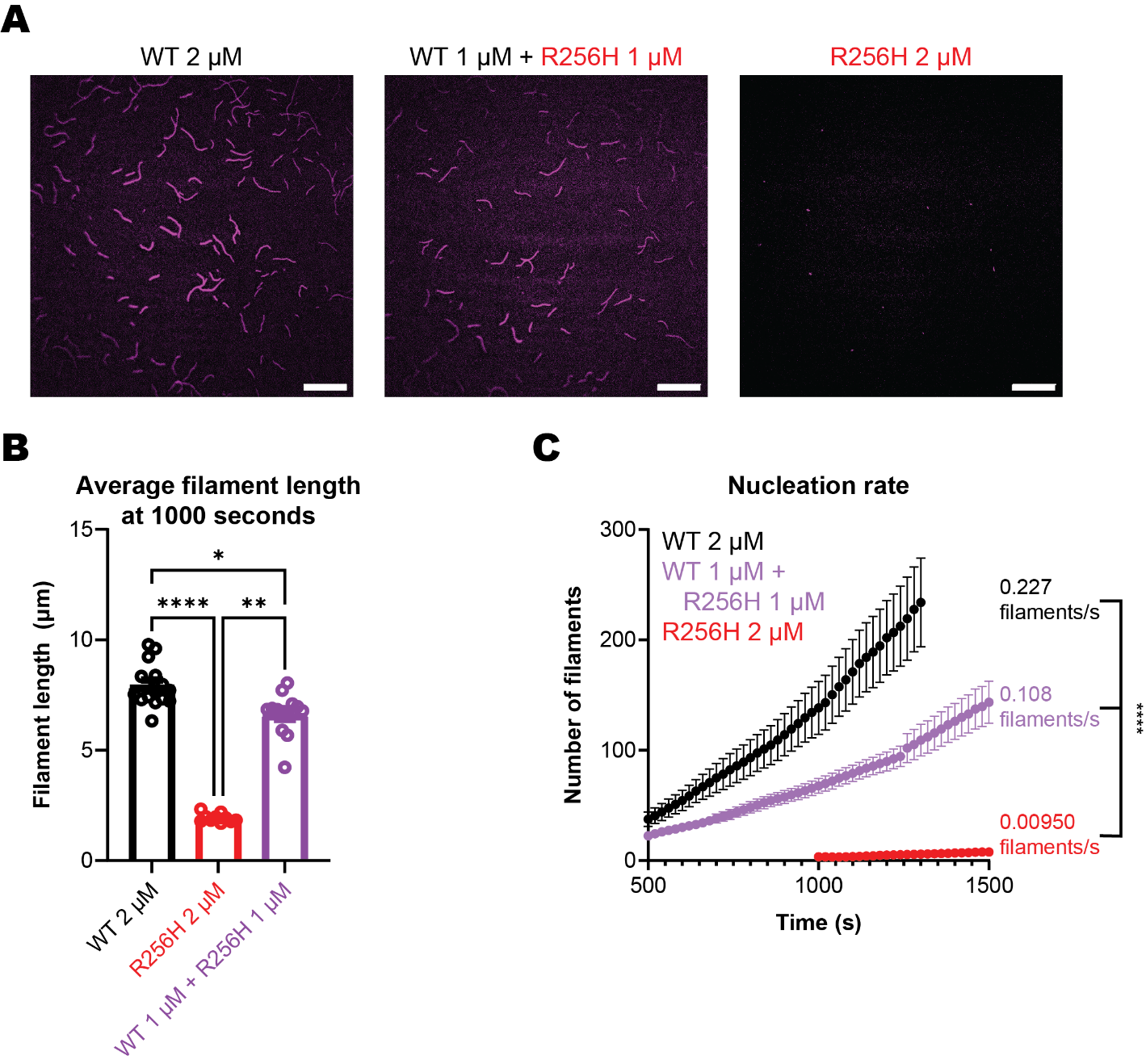

Supplementary Figure 3. Mixing WT and R256H ACTA1 results in intermediate filament length and nucleation rate. A) Representative image of a single frame at 1000 seconds from a fluorescent actin polymerization experiment using TIRF microscopy. Note that the WT and R256H images are a less-magnified version of the images seen in Figure 2D. Scale bar represents 20 μm. As mutant protein is substituted in the actin pool, the number of filaments and average filament length decreases. B) Average filament length is decreased significantly upon substitution of the total actin pool with R256H ACTA1. * = p < 0.05, ** = p < 0.01, **** = p < 0.0001 as calculated by one-way ANOVA with nonparametric Kruskal-Wallis test followed by Dunn correction. Data obtained from 10-15 different focal fields from 2-3 replicates per condition. C) Number of filaments over time using preceding data. Nucleation rates are significantly different between all groups with rates decreasing as ACTA1 R256H is substituted into the total actin pool. Nucleation rate was calculated by performing simple linear regression and significance of difference in slopes was calculated by GraphPad Prism. Note that WT and R256H data is reproduced from Figure 2D.

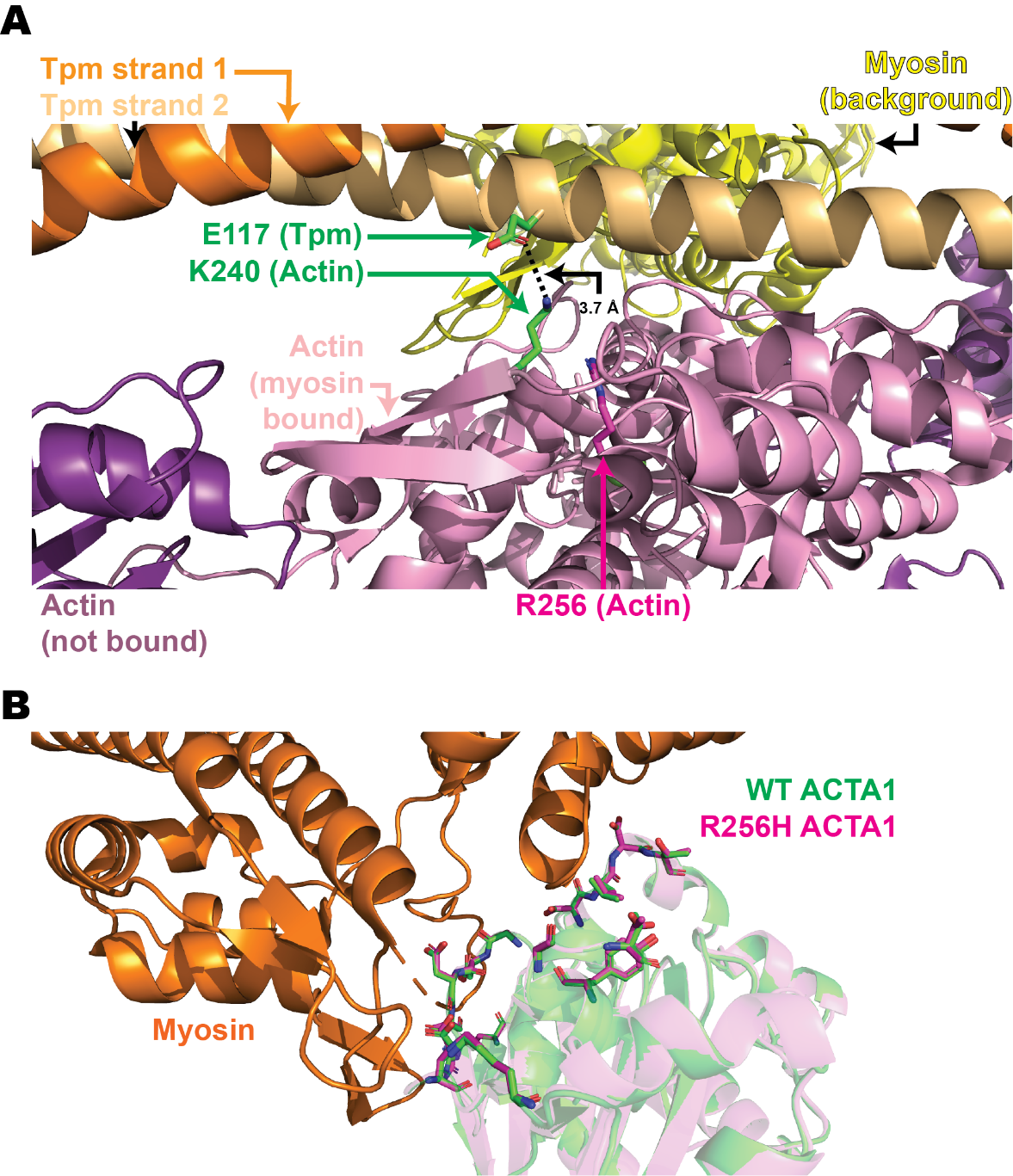

Supplementary Figure 4. A) Actin K240, near Actin R256, interacts with tropomyosin E117. Cartoon render of the published PDB 8EFI structure (68) which shows myosin bound to an actin filament in rigor in the presence of tropomyosin (Tpm). Individual proteins are labeled. Of note, the E117 residue on “Tpm strand 2” are shown as atomic sticks with a green carbon backbone and negatively charged oxygen shown in red. The K240 residue on the actin monomer (light purple) that is bound by myosin (yellow, in background) is also shown as atomic sticks with its positively charged nitrogen shown in blue. The ε-1 oxygen of Tpm E117 is approximately 3.7 Å from the ζ nitrogen actin K240 suggesting a salt bridge (shown as dashed black line). Notably, actin R256 is in close vicinity of K240 suggesting that this residue can influence the positioning of K240. B) R256H ACTA1 does not alter the actin-myosin interaction surface. Focused view of the WT and R256H ACTA1 side chains (shown as stick diagrams) that lay near the bound myosin. Note that there is little difference in the positions of these side chains. Myosin and WT and R256H ACTA1 residues that are not near myosin are rendered as a cartoon. Structure was obtained by aligning published 8EFI structure to our WT and R256H ACTA1 filaments.

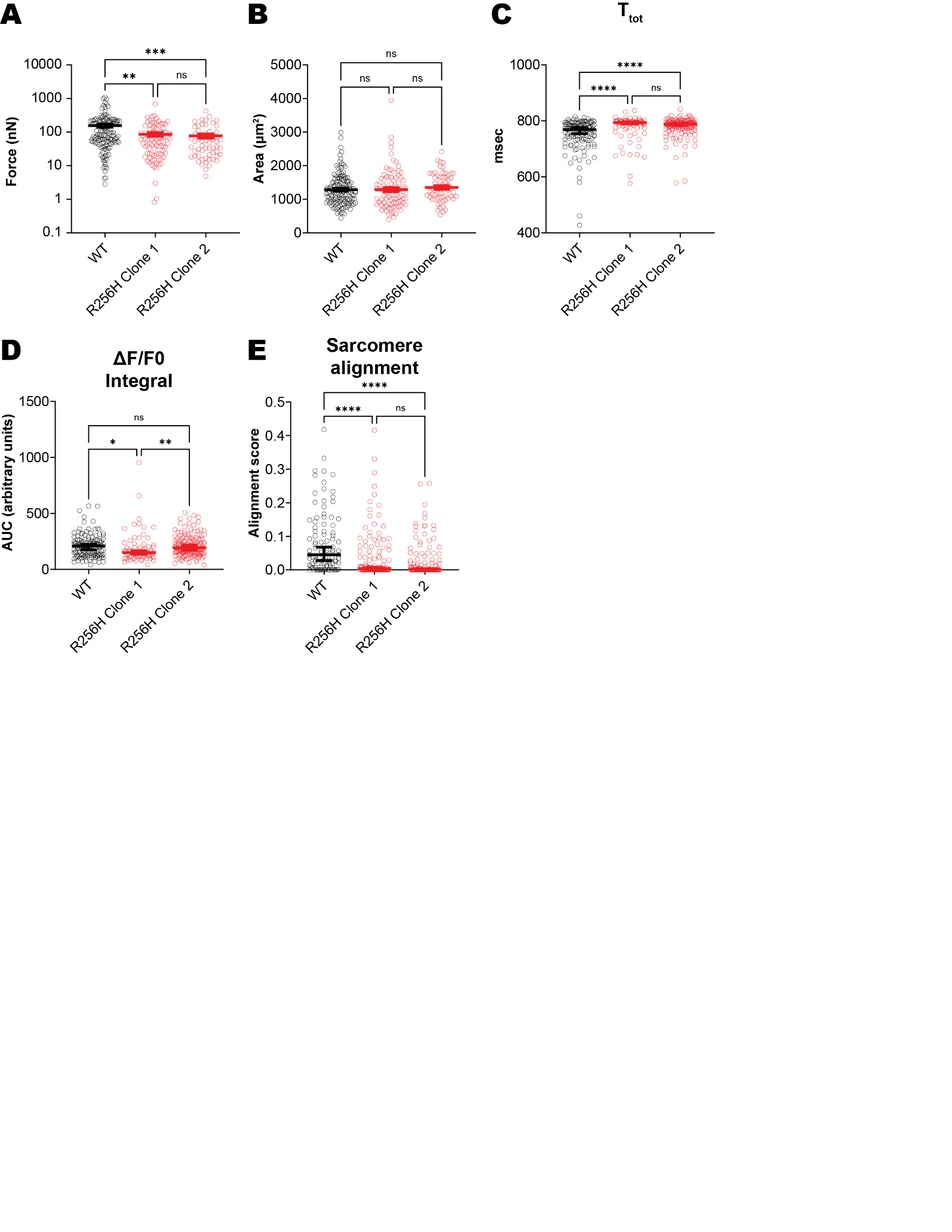

Supplementary Figure 5. *ACTA1^R256H/+^* hPSC-CMs separated data. A) Traction force data from Figure 5A and B) cell area data from Figure 5B with R256H clones separated. ** = p < 0.01 and *** = p < 0.001 as calculated by one-way ANOVA followed by non-parametric Kruskal-Wallis test with Dunn method for correction for multiple comparisons. WT *N=*139, R256H Clone 1 *N=*90, R256H Clone 2 *N=*59. C and D) Calcium transient data separated into two separate mutant lines from Figures 5E, 5F, and 5G. WT *N=*135, R256H Clone 1 *N=*74, R256H Clone 2 *N=*151. E) Sarcomere alignment scores from Figure 5I. WT *N=*99, R256H Clone 1 *N=*123, R256H Clone 2 *N=*135. * = p < 0.05, ** = p < 0.01, *** = p < 0.001, and **** = p < 0.0001 calculated by one-way ANOVA followed by Kruskal-Wallis analysis with multiple comparisons corrected by the Dunn method.

Dataset S1 (separate file). WT F-ACTA1 validation report generated by wwPDB Validation System.

Dataset S2 (separate file). R256H F-ACTA1 validation report generated by wwPDB Validation System.
