## Supplemental Dataset 1 for "Dilated cardiomyopathy-associated skeletal muscle actin (ACTA1) mutation R256H disrupts actin structure and function and causes cardiomyocyte hypocontractility"

### Preliminary Full wwPDB EM Validation Report (i)

Feb 22, 2024 – 11:43 AM EST

**This wwPDB validation report is NOT for manuscript review**

This is a Preliminary Full wwPDB EM Validation Report.

This report is produced by the standalone wwPDB validation server.
**The structure in question has not been deposited to the wwPDB.**
**This report should not be submitted to journals.**

We welcome your comments at **
A user guide is available at
https://www.wwpdb.org/validation/2017/EMValidationReportHelp
with specific help available everywhere you see the (i) symbol.

The types of validation reports are described at
http://www.wwpdb.org/validation/2017/FAQs#types.

The following versions of software and data (see references (i)) were used in the production of this report:

| | | |
|---|---|---|
| EMDB validation analysis | : | 0.0.1.dev70 |
| Mogul | : | 2022.3.0, CSD as543be (2022) |
| MolProbity | : | 4.02b-467 |
| buster-report | : | 1.1.7 (2018) |
| Percentile statistics | : | 20191225.v01 (using entries in the PDB archive December 25th 2019) |
| MapQ | : | 1.9.13 |
| Ideal geometry (proteins) | : | Engh & Huber (2001) |
| Ideal geometry (DNA, RNA) | : | Parkinson et al. (1996) |
| Validation Pipeline (wwPDB-VP) | : | 2.36 |

Not For Manuscript Review

### 1 Overall quality at a glance (i)

The following experimental techniques were used to determine the structure:
*ELECTRON MICROSCOPY*

The reported resolution of this entry is unknown.

Percentile scores (ranging between 0-100) for global validation metrics of the entry are shown in the following graphic. The table shows the number of entries on which the scores are based.

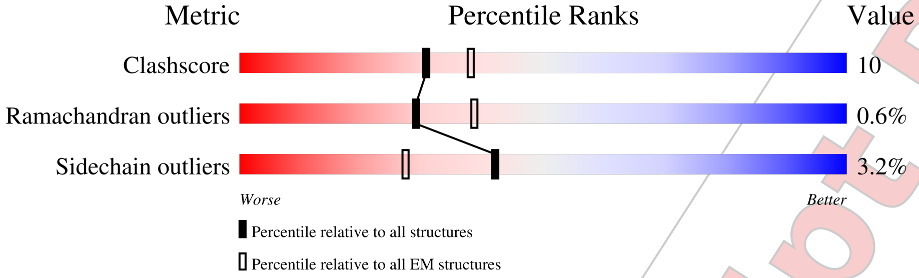


| Metric | Whole archive (#Entries) | EM structures (#Entries) |
|---|---|---|
| Clashscore | 158937 | 4297 |
| Ramachandran outliers | 154571 | 4023 |
| Sidechain outliers | 154315 | 3826 |

The table below summarises the geometric issues observed across the polymeric chains and their fit to the map. The red, orange, yellow and green segments of the bar indicate the fraction of residues that contain outliers for >=3, 2, 1 and 0 types of geometric quality criteria respectively. A grey segment represents the fraction of residues that are not modelled. The numeric value for each fraction is indicated below the corresponding segment, with a dot representing fractions <=5% The upper red bar (where present) indicates the fraction of residues that have poor fit to the EM map (all-atom inclusion < 40%). The numeric value is given above the bar.

| Mol | Chain | Length | Quality of chain |
|---|---|---|---|
| 1 | A | 371 | • / 80% 19% • |
| 1 | B | 371 | • / 78% 21% • |
| 1 | C | 371 | • / 79% 20% • |
| 1 | D | 371 | • / 78% 22% • |
| 1 | E | 371 | • / 80% 19% • |
| 1 | F | 371 | • / 80% 19% • |
| 2 | M | 7 | 14% 43% 29% 14% |
| 2 | N | 7 | 57% 29% 14% |

*Continued on next page...*

Not For Manuscript Review

*Continued from previous page...*

| Mol | Chain | Length | Quality of chain |
|---|---|---|---|
| 2 | O | 7 | 57% 29% 14% |
| 2 | P | 7 | 57% 29% 14% |
| 2 | Q | 7 | 57% 29% 14% |
| 2 | R | 7 | 14% 43% 29% 14% |
| 2 | S | 7 | 14% 43% 29% 14% |

The following table lists non-polymeric compounds, carbohydrate monomers and non-standard residues in protein, DNA, RNA chains that are outliers for geometric or electron-density-fit criteria:

| Mol | Type | Chain | Res | Chirality | Geometry | Clashes | Electron density |
|---|---|---|---|---|---|---|---|
| 2 | DTH | M | 4 | - | - | X | - |
| 2 | HYP | M | 6 | - | - | X | - |
| 2 | DTH | N | 4 | - | - | X | - |
| 2 | HYP | N | 6 | - | - | X | - |
| 2 | DTH | O | 4 | - | - | X | - |
| 2 | HYP | O | 6 | - | - | X | - |
| 2 | DTH | P | 4 | - | - | X | - |
| 2 | HYP | P | 6 | - | - | X | - |
| 2 | DTH | Q | 4 | - | - | X | - |
| 2 | HYP | Q | 6 | - | - | X | - |
| 2 | HYP | R | 6 | - | - | X | - |
| 2 | HYP | S | 6 | - | - | X | - |

Not For Manuscript Review

#### 2 Entry composition (i)

There are 4 unique types of molecules in this entry. The entry contains 17953 atoms, of which 0 are hydrogens and 0 are deuteriums.

In the tables below, the AltConf column contains the number of residues with at least one atom in alternate conformation and the Trace column contains the number of residues modelled with at most 2 atoms.

- Molecule 1 is a protein.

| Mol | Chain | Residues | Atoms | | | | | AltConf | Trace |
|---|---|---|---|---|---|---|---|---|---|
| 1 | A | 371 | Total 2900 | C 1837 | N 489 | O 553 | S 21 | 0 | 0 |
| 1 | B | 371 | Total 2900 | C 1837 | N 489 | O 553 | S 21 | 0 | 0 |
| 1 | C | 371 | Total 2900 | C 1837 | N 489 | O 553 | S 21 | 0 | 0 |
| 1 | D | 371 | Total 2900 | C 1837 | N 489 | O 553 | S 21 | 0 | 0 |
| 1 | E | 371 | Total 2900 | C 1837 | N 489 | O 553 | S 21 | 0 | 0 |
| 1 | F | 371 | Total 2900 | C 1837 | N 489 | O 553 | S 21 | 0 | 0 |

- Molecule 2 is a protein called TRP-EEP-ALA-DTH-CYS-HYP-ALA.

| Mol | Chain | Residues | Atoms | | | | | AltConf | Trace |
|---|---|---|---|---|---|---|---|---|---|
| 2 | M | 7 | Total 55 | C 35 | N 8 | O 11 | S 1 | 0 | 0 |
| 2 | N | 7 | Total 55 | C 35 | N 8 | O 11 | S 1 | 0 | 0 |
| 2 | O | 7 | Total 55 | C 35 | N 8 | O 11 | S 1 | 0 | 0 |
| 2 | P | 7 | Total 55 | C 35 | N 8 | O 11 | S 1 | 0 | 0 |
| 2 | Q | 7 | Total 55 | C 35 | N 8 | O 11 | S 1 | 0 | 0 |
| 2 | R | 7 | Total 55 | C 35 | N 8 | O 11 | S 1 | 0 | 0 |
| 2 | S | 7 | Total 55 | C 35 | N 8 | O 11 | S 1 | 0 | 0 |

- Molecule 3 is ADENOSINE-5'-DIPHOSPHATE (three-letter code: ADP) (formula: $C_{10}H_{15}N_5O_{10}P_2$).

Not For Manuscript Review

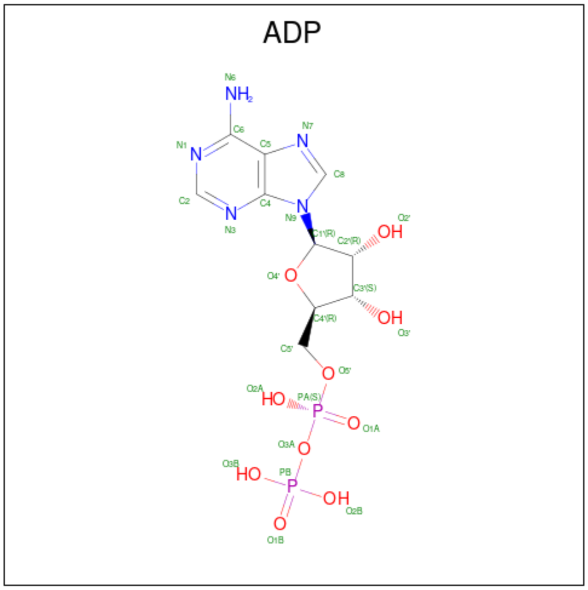


| Mol | Chain | Residues | Atoms | | | | | AltConf |
|---|---|---|---|---|---|---|---|---|
| 3 | G | 1 | Total 27 | C 10 | N 5 | O 10 | P 2 | 0 |
| 3 | H | 1 | Total 27 | C 10 | N 5 | O 10 | P 2 | 0 |
| 3 | I | 1 | Total 27 | C 10 | N 5 | O 10 | P 2 | 0 |
| 3 | J | 1 | Total 27 | C 10 | N 5 | O 10 | P 2 | 0 |
| 3 | K | 1 | Total 27 | C 10 | N 5 | O 10 | P 2 | 0 |
| 3 | L | 1 | Total 27 | C 10 | N 5 | O 10 | P 2 | 0 |

- Molecule 4 is MAGNESIUM ION (three-letter code: MG) (formula: Mg).

| Mol | Chain | Residues | Atoms | | AltConf |
|---|---|---|---|---|---|
| 4 | G | 1 | Total 1 | Mg 1 | 0 |
| 4 | H | 1 | Total 1 | Mg 1 | 0 |
| 4 | I | 1 | Total 1 | Mg 1 | 0 |
| 4 | J | 1 | Total 1 | Mg 1 | 0 |
| 4 | K | 1 | Total 1 | Mg 1 | 0 |
| 4 | L | 1 | Total 1 | Mg 1 | 0 |

Not For Manuscript Review

### 3 Residue-property plots (i)

These plots are drawn for all protein, RNA, DNA and oligosaccharide chains in the entry. The first graphic for a chain summarises the proportions of the various outlier classes displayed in the second graphic. The second graphic shows the sequence view annotated by issues in geometry and atom inclusion in map density. Residues are color-coded according to the number of geometric quality criteria for which they contain at least one outlier: green = 0, yellow = 1, orange = 2 and red = 3 or more. A red diamond above a residue indicates a poor fit to the EM map for this residue (all-atom inclusion < 40%). Stretches of 2 or more consecutive residues without any outlier are shown as a green connector. Residues present in the sample, but not in the model, are shown in grey.

- Molecule 1:

Chain A:

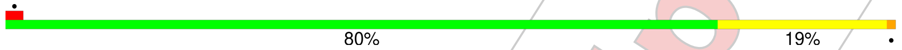


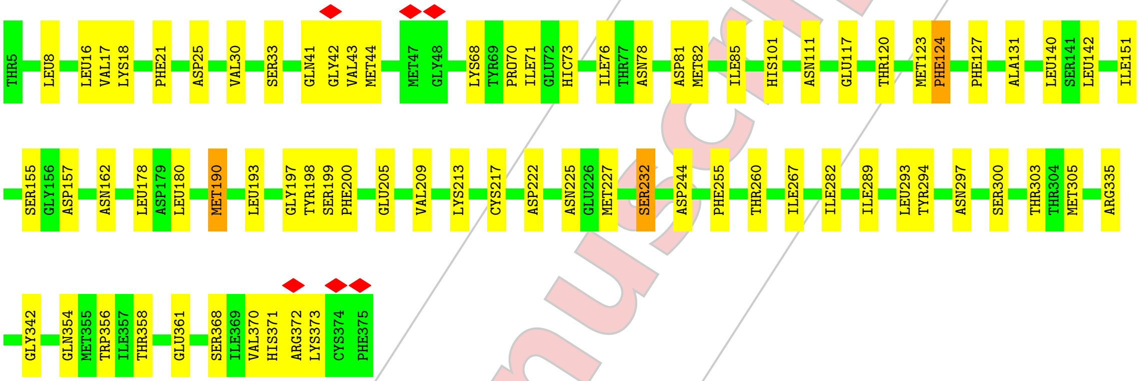


- Molecule 1:

Chain B:

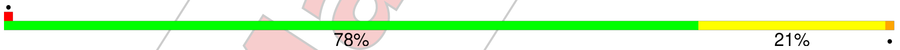


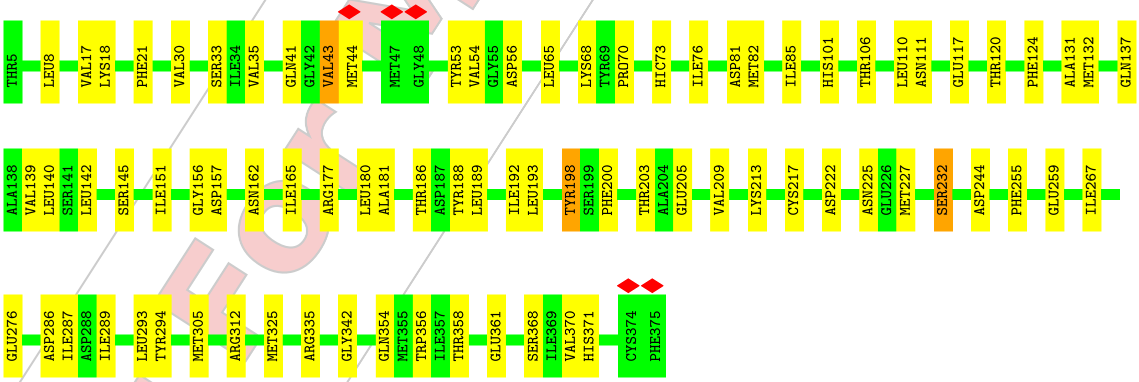


- Molecule 1:

Chain C:

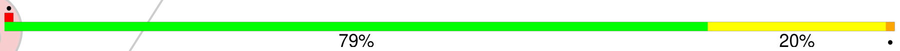


Not For Manuscript Review

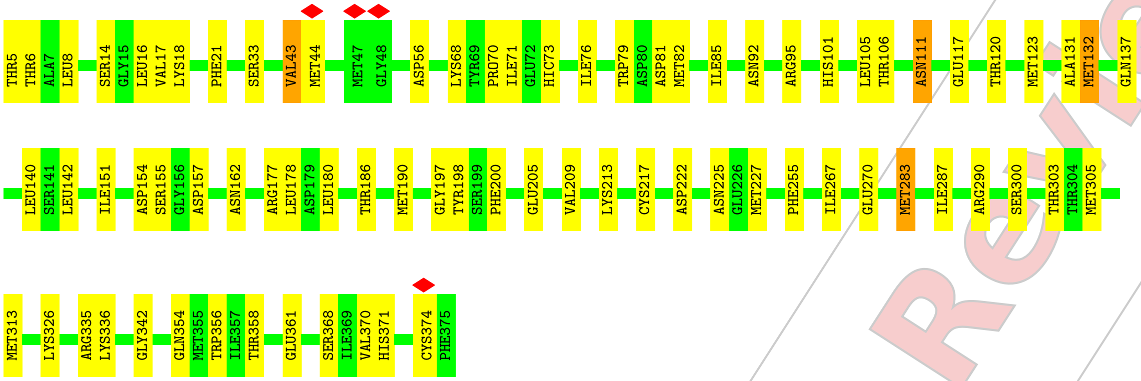


• Molecule 1:

Chain D:

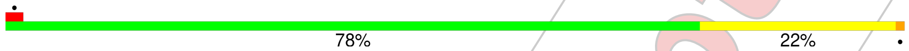


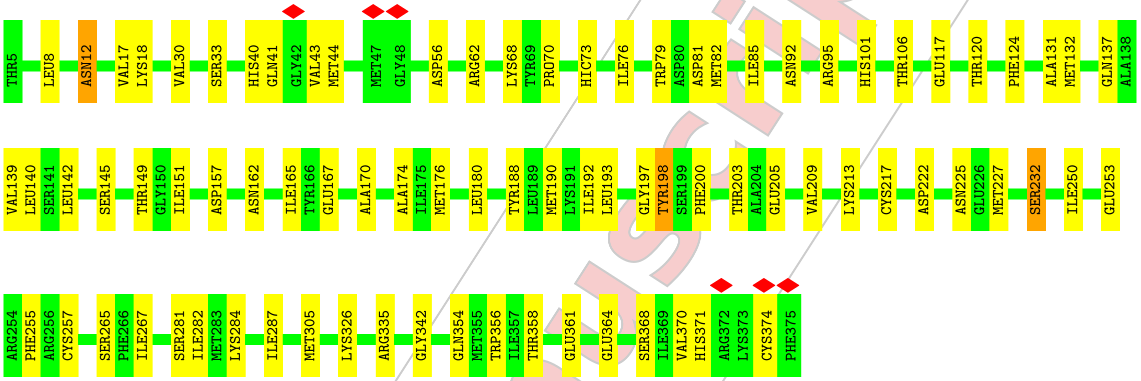


• Molecule 1:

Chain E:

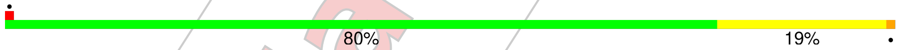


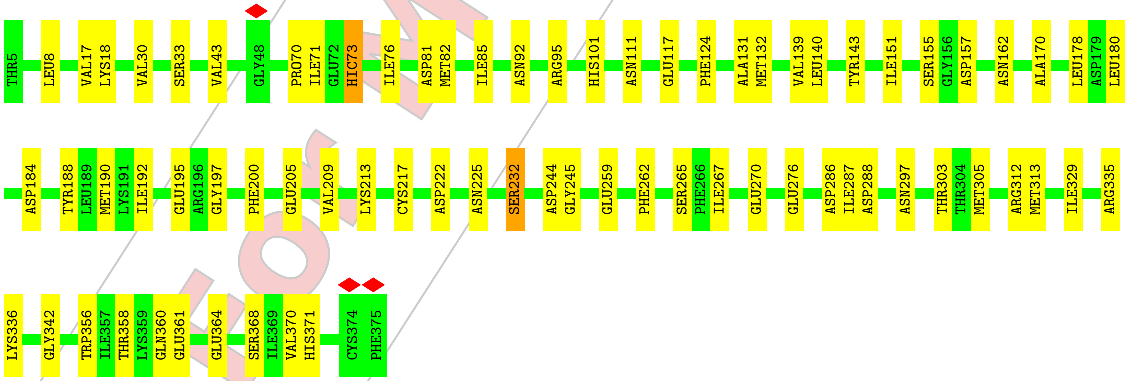


• Molecule 1:

Chain F:

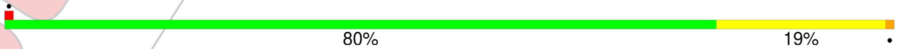


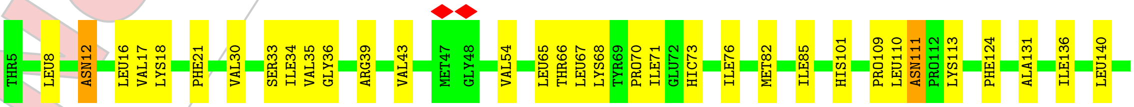


Not For Manuscript Review

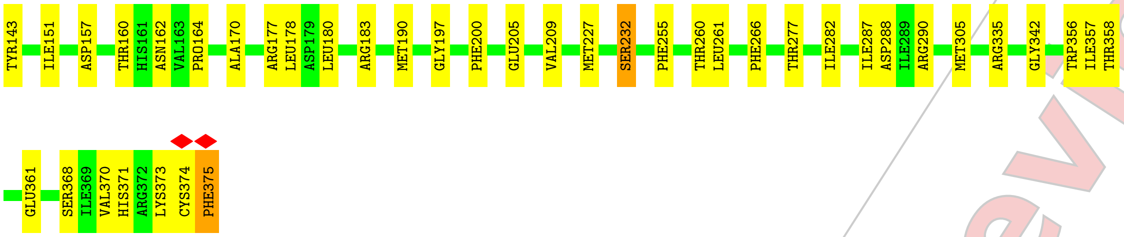


- Molecule 2: TRP-EEP-ALA-DTH-CYS-HYP-ALA

Chain M: 14% 43% 29% 14%

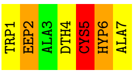


- Molecule 2: TRP-EEP-ALA-DTH-CYS-HYP-ALA

Chain N: 57% 29% 14%

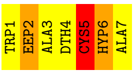


- Molecule 2: TRP-EEP-ALA-DTH-CYS-HYP-ALA

Chain O: 57% 29% 14%

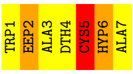


- Molecule 2: TRP-EEP-ALA-DTH-CYS-HYP-ALA

Chain P: 57% 29% 14%

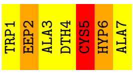


- Molecule 2: TRP-EEP-ALA-DTH-CYS-HYP-ALA

Chain Q: 57% 29% 14%

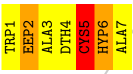


- Molecule 2: TRP-EEP-ALA-DTH-CYS-HYP-ALA

Chain R: 14% 43% 29% 14%

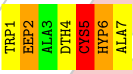


- Molecule 2: TRP-EEP-ALA-DTH-CYS-HYP-ALA

Chain S: 14% 43% 29% 14%

Not For Manuscript Review

TRP1 EEP2 ALA3 DTH4 CYS5 HYP6 ALA7

Not For Manuscript Review

### 4 Experimental information (i)

| Property | Value | Source |
|---|---|---|
| EM reconstruction method | HELICAL | Depositor |
| Imposed symmetry | POINT, Not provided | |
| Number of segments used | Not provided | |
| Resolution determination method | Not provided | |
| CTF correction method | Not provided | |
| Microscope | Not provided | |
| Voltage (kV) | Not provided | |
| Electron dose $(e^-/\text{Å}^2)$ | Not provided | |
| Minimum defocus (nm) | Not provided | |
| Maximum defocus (nm) | Not provided | |
| Magnification | Not provided | |
| Image detector | Not provided | |
| Maximum map value | 7.167 | Depositor |
| Minimum map value | -4.599 | Depositor |
| Average map value | 0.003 | Depositor |
| Map value standard deviation | 0.148 | Depositor |
| Recommended contour level | 0.9 | Depositor |
| Map size (Å) | 473.6, 473.6, 473.6 | wwPDB |
| Map dimensions | 400, 400, 400 | wwPDB |
| Map angles (°) | 90.0, 90.0, 90.0 | wwPDB |
| Pixel spacing (Å) | 1.184, 1.184, 1.184 | Depositor |

Not For Manuscript Review

### 5 Model quality (i)

#### 5.1 Standard geometry (i)

Bond lengths and bond angles in the following residue types are not validated in this section: DTH, HIC, HYP, ADP, EEP, MG

The Z score for a bond length (or angle) is the number of standard deviations the observed value is removed from the expected value. A bond length (or angle) with $|Z| > 5$ is considered an outlier worth inspection. RMSZ is the root-mean-square of all Z scores of the bond lengths (or angles).

| Mol | Chain | Bond lengths | | Bond angles | |
|---|---|---|---|---|---|
| | | RMSZ | #$\|Z\|$ >5 | RMSZ | #$\|Z\|$ >5 |
| 1 | A | 0.26 | 0/2950 | 0.48 | 0/3994 |
| 1 | B | 0.26 | 0/2950 | 0.47 | 0/3994 |
| 1 | C | 0.26 | 0/2950 | 0.47 | 0/3994 |
| 1 | D | 0.26 | 0/2950 | 0.48 | 0/3994 |
| 1 | E | 0.25 | 0/2950 | 0.47 | 0/3994 |
| 1 | F | 0.27 | 0/2950 | 0.47 | 0/3994 |
| 2 | M | 1.03 | 0/28 | 1.40 | 0/33 |
| 2 | N | 1.03 | 0/28 | 1.43 | 0/33 |
| 2 | O | 1.04 | 0/28 | 1.40 | 0/33 |
| 2 | P | 1.04 | 0/28 | 1.40 | 0/33 |
| 2 | Q | 1.03 | 0/28 | 1.40 | 0/33 |
| 2 | R | 1.04 | 0/28 | 1.40 | 0/33 |
| 2 | S | 1.03 | 0/28 | 1.43 | 0/33 |
| All | All | 0.28 | 0/17896 | 0.49 | 0/24195 |

There are no bond length outliers.

There are no bond angle outliers.

There are no chirality outliers.

There are no planarity outliers.

#### 5.2 Too-close contacts (i)

In the following table, the Non-H and H(model) columns list the number of non-hydrogen atoms and hydrogen atoms in the chain respectively. The H(added) column lists the number of hydrogen atoms added and optimized by MolProbity. The Clashes column lists the number of clashes within the asymmetric unit, whereas Symm-Clashes lists symmetry-related clashes.

| Mol | Chain | Non-H | H(model) | H(added) | Clashes | Symm-Clashes |
|---|---|---|---|---|---|---|
| 1 | A | 2900 | 0 | 2872 | 48 | 0 |

*Continued on next page...*

Not For Manuscript Review

*Continued from previous page...*

| Mol | Chain | Non-H | H(model) | H(added) | Clashes | Symm-Clashes |
|---|---|---|---|---|---|---|
| 1 | B | 2900 | 0 | 2872 | 53 | 0 |
| 1 | C | 2900 | 0 | 2872 | 55 | 0 |
| 1 | D | 2900 | 0 | 2872 | 57 | 0 |
| 1 | E | 2900 | 0 | 2872 | 46 | 0 |
| 1 | F | 2900 | 0 | 2872 | 50 | 0 |
| 2 | M | 55 | 0 | 37 | 11 | 0 |
| 2 | N | 55 | 0 | 37 | 12 | 0 |
| 2 | O | 55 | 0 | 37 | 16 | 0 |
| 2 | P | 55 | 0 | 37 | 15 | 0 |
| 2 | Q | 55 | 0 | 37 | 12 | 0 |
| 2 | R | 55 | 0 | 37 | 10 | 0 |
| 2 | S | 55 | 0 | 37 | 9 | 0 |
| 3 | G | 27 | 0 | 12 | 1 | 0 |
| 3 | H | 27 | 0 | 12 | 3 | 0 |
| 3 | I | 27 | 0 | 12 | 3 | 0 |
| 3 | J | 27 | 0 | 12 | 1 | 0 |
| 3 | K | 27 | 0 | 12 | 3 | 0 |
| 3 | L | 27 | 0 | 12 | 1 | 0 |
| 4 | G | 1 | 0 | 0 | 0 | 0 |
| 4 | H | 1 | 0 | 0 | 0 | 0 |
| 4 | I | 1 | 0 | 0 | 0 | 0 |
| 4 | J | 1 | 0 | 0 | 0 | 0 |
| 4 | K | 1 | 0 | 0 | 0 | 0 |
| 4 | L | 1 | 0 | 0 | 0 | 0 |
| All | All | 17953 | 0 | 17563 | 348 | 0 |

The all-atom clashscore is defined as the number of clashes found per 1000 atoms (including hydrogen atoms). The all-atom clashscore for this structure is 10.

All (348) close contacts within the same asymmetric unit are listed below, sorted by their clash magnitude.

| Atom-1 | Atom-2 | Interatomic distance (Å) | Clash overlap (Å) |
|---|---|---|---|
| 1:B:200:PHE:HB3 | 1:B:205:GLU:HB3 | 1.54 | 0.90 |
| 1:A:200:PHE:HB3 | 1:A:205:GLU:HB3 | 1.53 | 0.89 |
| 1:F:287:ILE:HD12 | 2:Q:4:DTH:CG2 | 2.05 | 0.87 |
| 2:Q:2:EEP:N | 2:Q:2:EEP:O1 | 2.07 | 0.87 |
| 2:S:2:EEP:O1 | 2:S:2:EEP:N | 2.07 | 0.85 |
| 1:D:200:PHE:HB3 | 1:D:205:GLU:HB3 | 1.58 | 0.84 |
| 2:O:2:EEP:N | 2:O:2:EEP:O1 | 2.07 | 0.84 |
| 1:D:198:TYR:HB3 | 2:Q:3:ALA:HB3 | 1.58 | 0.84 |
| 2:N:2:EEP:O1 | 2:N:2:EEP:N | 2.07 | 0.84 |

*Continued on next page...*

Not For Manuscript Review

Continued from previous page...

| Atom-1 | Atom-2 | Interatomic distance (Å) | Clash overlap (Å) |
|---|---|---|---|
| 2:P:2:EEP:N | 2:P:2:EEP:O1 | 2.07 | 0.84 |
| 2:R:2:EEP:N | 2:R:2:EEP:O1 | 2.07 | 0.84 |
| 1:F:287:ILE:HD12 | 2:Q:4:DTH:HG21 | 1.58 | 0.82 |
| 2:M:2:EEP:N | 2:M:2:EEP:O1 | 2.07 | 0.81 |
| 1:F:151:ILE:HD13 | 1:F:282:ILE:HG13 | 1.64 | 0.78 |
| 2:R:6:HYP:O | 2:R:7:ALA:C | 2.22 | 0.77 |
| 2:P:6:HYP:O | 2:P:7:ALA:C | 2.22 | 0.77 |
| 2:O:6:HYP:O | 2:O:7:ALA:C | 2.22 | 0.76 |
| 2:M:6:HYP:O | 2:M:7:ALA:C | 2.22 | 0.75 |
| 2:N:6:HYP:O | 2:N:7:ALA:C | 2.23 | 0.75 |
| 1:E:287:ILE:HD12 | 2:P:4:DTH:HG21 | 1.66 | 0.75 |
| 1:C:197:GLY:O | 2:P:1:TRP:CE3 | 2.40 | 0.74 |
| 2:S:6:HYP:O | 2:S:7:ALA:C | 2.22 | 0.74 |
| 2:Q:6:HYP:O | 2:Q:7:ALA:C | 2.22 | 0.74 |
| 1:D:287:ILE:CD1 | 2:O:4:DTH:HG21 | 2.20 | 0.72 |
| 1:B:198:TYR:HB3 | 2:O:3:ALA:HB3 | 1.72 | 0.71 |
| 1:A:180:LEU:HD11 | 1:A:267:ILE:HD11 | 1.72 | 0.71 |
| 2:M:4:DTH:O | 2:M:5:CYS:O | 2.09 | 0.71 |
| 2:O:4:DTH:O | 2:O:5:CYS:O | 2.09 | 0.71 |
| 2:P:4:DTH:O | 2:P:5:CYS:O | 2.09 | 0.70 |
| 2:Q:4:DTH:O | 2:Q:5:CYS:O | 2.09 | 0.70 |
| 2:R:4:DTH:O | 2:R:5:CYS:O | 2.09 | 0.70 |
| 1:C:197:GLY:O | 2:P:1:TRP:HE3 | 1.75 | 0.70 |
| 2:N:4:DTH:O | 2:N:5:CYS:O | 2.09 | 0.70 |
| 2:S:4:DTH:O | 2:S:5:CYS:O | 2.09 | 0.69 |
| 1:A:17:VAL:HG23 | 1:A:33:SER:HB2 | 1.75 | 0.68 |
| 1:B:17:VAL:HG23 | 1:B:33:SER:HB2 | 1.77 | 0.67 |
| 1:A:76:ILE:HD13 | 1:A:82:MET:HG2 | 1.76 | 0.66 |
| 1:F:287:ILE:HA | 1:F:290:ARG:HE | 1.61 | 0.66 |
| 1:F:305:MET:HA | 1:F:335:ARG:HH21 | 1.60 | 0.66 |
| 1:C:283:MET:SD | 1:C:290:ARG:NH1 | 2.69 | 0.65 |
| 1:D:190:MET:HG2 | 1:D:209:VAL:HG21 | 1.78 | 0.65 |
| 1:E:197:GLY:O | 2:R:1:TRP:CE3 | 2.50 | 0.65 |
| 1:D:287:ILE:CD1 | 2:O:4:DTH:CG2 | 2.74 | 0.65 |
| 1:D:17:VAL:HG23 | 1:D:33:SER:HB2 | 1.79 | 0.65 |
| 1:F:70:PRO:HG2 | 1:F:85:ILE:HD12 | 1.79 | 0.64 |
| 1:E:287:ILE:HD12 | 2:P:4:DTH:CG2 | 2.26 | 0.64 |
| 2:O:1:TRP:N | 2:O:6:HYP:O | 2.31 | 0.64 |
| 2:Q:1:TRP:N | 2:Q:6:HYP:O | 2.31 | 0.64 |
| 2:P:1:TRP:N | 2:P:6:HYP:O | 2.31 | 0.64 |
| 2:S:1:TRP:N | 2:S:6:HYP:O | 2.31 | 0.64 |

Continued on next page...

Not For Manuscript Review

*Continued from previous page...*

| Atom-1 | Atom-2 | Interatomic distance (Å) | Clash overlap (Å) |
|---|---|---|---|
| 1:D:287:ILE:HD13 | 2:O:4:DTH:CG2 | 2.28 | 0.64 |
| 1:D:287:ILE:HD13 | 2:O:4:DTH:HG21 | 1.79 | 0.64 |
| 2:M:1:TRP:N | 2:M:6:HYP:O | 2.31 | 0.64 |
| 2:R:1:TRP:N | 2:R:6:HYP:O | 2.31 | 0.64 |
| 1:F:110:LEU:O | 1:F:177:ARG:NH1 | 2.31 | 0.63 |
| 1:B:41:GLN:NE2 | 1:D:374:CYS:O | 2.25 | 0.63 |
| 2:N:1:TRP:N | 2:N:6:HYP:O | 2.31 | 0.63 |
| 1:C:17:VAL:HG23 | 1:C:33:SER:HB2 | 1.80 | 0.63 |
| 1:C:200:PHE:HB3 | 1:C:205:GLU:HB3 | 1.81 | 0.63 |
| 1:D:70:PRO:HG2 | 1:D:85:ILE:HD12 | 1.80 | 0.63 |
| 1:C:43:VAL:HG21 | 1:E:170:ALA:H | 1.64 | 0.63 |
| 1:A:227:MET:HG2 | 1:A:255:PHE:HE2 | 1.63 | 0.62 |
| 1:E:180:LEU:HD11 | 1:E:267:ILE:HD11 | 1.81 | 0.62 |
| 1:C:180:LEU:HD21 | 1:C:267:ILE:HD11 | 1.82 | 0.62 |
| 1:C:305:MET:HA | 1:C:335:ARG:HH21 | 1.65 | 0.62 |
| 1:F:180:LEU:HB3 | 1:F:261:LEU:HD23 | 1.82 | 0.61 |
| 1:A:305:MET:HA | 1:A:335:ARG:HH21 | 1.66 | 0.61 |
| 1:B:44:MET:HB3 | 1:D:139:VAL:HG11 | 1.83 | 0.61 |
| 1:E:17:VAL:HG23 | 1:E:33:SER:HB2 | 1.81 | 0.61 |
| 1:B:305:MET:HA | 1:B:335:ARG:HH21 | 1.65 | 0.61 |
| 1:E:287:ILE:CD1 | 2:P:4:DTH:HG21 | 2.30 | 0.60 |
| 1:F:17:VAL:HG23 | 1:F:33:SER:HB2 | 1.82 | 0.60 |
| 1:E:336:LYS:HE3 | 3:K:376:ADP:H8 | 1.67 | 0.60 |
| 1:A:131:ALA:HB1 | 1:A:356:TRP:HB3 | 1.82 | 0.60 |
| 1:B:287:ILE:HD12 | 2:M:4:DTH:CG2 | 2.32 | 0.60 |
| 1:C:120:THR:HG23 | 1:C:132:MET:HE1 | 1.84 | 0.59 |
| 1:D:151:ILE:HD11 | 1:D:162:ASN:HD21 | 1.67 | 0.59 |
| 1:B:227:MET:HG2 | 1:B:255:PHE:HE2 | 1.68 | 0.59 |
| 1:C:44:MET:HB3 | 1:E:139:VAL:HG11 | 1.84 | 0.58 |
| 1:E:200:PHE:HB3 | 1:E:205:GLU:HB3 | 1.84 | 0.58 |
| 1:E:131:ALA:HB1 | 1:E:356:TRP:HB3 | 1.86 | 0.58 |
| 1:D:305:MET:HA | 1:D:335:ARG:HH21 | 1.68 | 0.58 |
| 1:C:131:ALA:HB1 | 1:C:356:TRP:HB3 | 1.85 | 0.58 |
| 1:B:43:VAL:HG21 | 1:D:170:ALA:N | 2.19 | 0.58 |
| 1:D:287:ILE:HD12 | 2:O:4:DTH:HG21 | 1.85 | 0.58 |
| 1:C:190:MET:HG3 | 1:C:209:VAL:HG11 | 1.86 | 0.57 |
| 1:E:190:MET:HG2 | 1:E:209:VAL:HG11 | 1.84 | 0.57 |
| 1:D:176:MET:HG3 | 1:D:281:SER:HB2 | 1.87 | 0.57 |
| 1:B:368:SER:O | 1:B:368:SER:OG | 2.22 | 0.57 |
| 1:D:151:ILE:HD13 | 1:D:282:ILE:HG13 | 1.87 | 0.57 |
| 1:B:232:SER:O | 1:B:232:SER:OG | 2.23 | 0.57 |

*Continued on next page...*

Not For Manuscript Review

*Continued from previous page...*

| Atom-1 | Atom-2 | Interatomic distance (Å) | Clash overlap (Å) |
|---|---|---|---|
| 1:F:136:ILE:HG13 | 1:F:375:PHE:HB3 | 1.86 | 0.57 |
| 1:E:70:PRO:HG2 | 1:E:85:ILE:HD12 | 1.87 | 0.57 |
| 1:F:200:PHE:HB3 | 1:F:205:GLU:HB3 | 1.85 | 0.57 |
| 1:F:227:MET:HG2 | 1:F:255:PHE:HE1 | 1.70 | 0.56 |
| 1:D:76:ILE:HD13 | 1:D:82:MET:HG2 | 1.86 | 0.56 |
| 1:E:244:ASP:OD1 | 1:E:245:GLY:N | 2.37 | 0.56 |
| 1:A:41:GLN:NE2 | 1:C:374:CYS:O | 2.37 | 0.56 |
| 1:D:117:GLU:OE2 | 1:D:371:HIS:NE2 | 2.36 | 0.56 |
| 1:B:180:LEU:HD21 | 1:B:267:ILE:HD11 | 1.87 | 0.56 |
| 1:A:81:ASP:O | 1:A:85:ILE:HD12 | 2.05 | 0.56 |
| 1:C:43:VAL:HG21 | 1:E:170:ALA:N | 2.21 | 0.56 |
| 1:C:287:ILE:HD12 | 2:N:4:DTH:CG2 | 2.37 | 0.55 |
| 1:E:336:LYS:HE3 | 3:K:376:ADP:C8 | 2.41 | 0.55 |
| 1:A:71:ILE:HG12 | 1:A:76:ILE:HG12 | 1.88 | 0.55 |
| 1:C:370:VAL:O | 1:C:371:HIS:ND1 | 2.40 | 0.55 |
| 1:C:222:ASP:OD2 | 1:C:225:ASN:ND2 | 2.38 | 0.54 |
| 1:C:151:ILE:HD11 | 1:C:162:ASN:HD21 | 1.71 | 0.54 |
| 1:D:180:LEU:HD21 | 1:D:267:ILE:HD11 | 1.89 | 0.54 |
| 1:D:203:THR:OG1 | 1:E:270:GLU:OE2 | 2.22 | 0.54 |
| 1:F:370:VAL:O | 1:F:371:HIS:ND1 | 2.41 | 0.54 |
| 2:O:4:DTH:C | 2:O:5:CYS:O | 2.56 | 0.54 |
| 1:D:370:VAL:O | 1:D:371:HIS:ND1 | 2.41 | 0.54 |
| 1:A:190:MET:HG2 | 1:A:209:VAL:HG11 | 1.90 | 0.54 |
| 1:F:197:GLY:O | 2:S:1:TRP:HE3 | 1.89 | 0.54 |
| 2:Q:4:DTH:C | 2:Q:5:CYS:O | 2.56 | 0.54 |
| 2:S:4:DTH:C | 2:S:5:CYS:O | 2.56 | 0.53 |
| 2:R:4:DTH:C | 2:R:5:CYS:O | 2.56 | 0.53 |
| 2:N:4:DTH:C | 2:N:5:CYS:O | 2.56 | 0.53 |
| 1:C:70:PRO:HG2 | 1:C:85:ILE:HD12 | 1.90 | 0.53 |
| 1:C:76:ILE:HG23 | 1:C:82:MET:HG3 | 1.91 | 0.53 |
| 1:D:41:GLN:HB3 | 1:F:170:ALA:HB3 | 1.91 | 0.53 |
| 1:D:62:ARG:NE | 1:F:288:ASP:OD2 | 2.41 | 0.53 |
| 1:E:222:ASP:OD2 | 1:E:225:ASN:ND2 | 2.38 | 0.53 |
| 1:B:370:VAL:O | 1:B:371:HIS:ND1 | 2.42 | 0.53 |
| 1:E:287:ILE:CD1 | 2:P:4:DTH:CG2 | 2.87 | 0.53 |
| 2:M:4:DTH:C | 2:M:5:CYS:O | 2.56 | 0.53 |
| 1:E:232:SER:O | 1:E:232:SER:OG | 2.23 | 0.53 |
| 1:E:305:MET:HA | 1:E:335:ARG:HH21 | 1.74 | 0.53 |
| 1:F:131:ALA:HB1 | 1:F:356:TRP:HB3 | 1.91 | 0.53 |
| 1:F:305:MET:SD | 1:F:335:ARG:NH2 | 2.82 | 0.52 |
| 1:C:287:ILE:HD12 | 2:N:4:DTH:HG21 | 1.90 | 0.52 |

*Continued on next page...*

Not For Manuscript Review

Continued from previous page...

| Atom-1 | Atom-2 | Interatomic distance (Å) | Clash overlap (Å) |
|---|---|---|---|
| 2:P:4:DTH:C | 2:P:5:CYS:O | 2.56 | 0.52 |
| 1:A:222:ASP:OD2 | 1:A:225:ASN:ND2 | 2.39 | 0.52 |
| 1:D:131:ALA:HB1 | 1:D:356:TRP:HB3 | 1.91 | 0.52 |
| 1:B:53:TYR:HD2 | 1:B:65:LEU:HD21 | 1.74 | 0.52 |
| 1:C:140:LEU:O | 1:C:342:GLY:HA3 | 2.09 | 0.52 |
| 1:B:222:ASP:OD2 | 1:B:225:ASN:ND2 | 2.37 | 0.52 |
| 1:B:140:LEU:O | 1:B:342:GLY:HA3 | 2.10 | 0.52 |
| 1:B:193:LEU:HB3 | 1:B:198:TYR:O | 2.10 | 0.52 |
| 1:A:358:THR:OG1 | 1:A:361:GLU:OE1 | 2.27 | 0.52 |
| 1:D:232:SER:O | 1:D:232:SER:OG | 2.23 | 0.51 |
| 1:D:287:ILE:HD12 | 2:O:4:DTH:CG2 | 2.39 | 0.51 |
| 1:C:368:SER:O | 1:C:368:SER:OG | 2.26 | 0.51 |
| 1:A:370:VAL:HG13 | 1:A:373:LYS:HG2 | 1.92 | 0.51 |
| 1:F:12:ASN:ND2 | 1:F:12:ASN:O | 2.42 | 0.51 |
| 1:B:43:VAL:HG21 | 1:D:170:ALA:H | 1.74 | 0.51 |
| 1:A:198:TYR:HB3 | 2:N:3:ALA:HB3 | 1.91 | 0.51 |
| 1:A:140:LEU:O | 1:A:342:GLY:HA3 | 2.10 | 0.51 |
| 1:B:131:ALA:HB1 | 1:B:356:TRP:HB3 | 1.92 | 0.51 |
| 1:B:203:THR:OG1 | 1:C:270:GLU:OE2 | 2.20 | 0.51 |
| 1:B:287:ILE:HD12 | 2:M:4:DTH:HG21 | 1.93 | 0.51 |
| 1:C:198:TYR:HB3 | 2:P:3:ALA:HB3 | 1.93 | 0.51 |
| 1:D:188:TYR:O | 1:D:192:ILE:HG23 | 2.11 | 0.51 |
| 1:A:155:SER:HG | 1:A:303:THR:HG1 | 1.57 | 0.50 |
| 1:F:12:ASN:C | 1:F:12:ASN:HD22 | 2.13 | 0.50 |
| 1:A:8:LEU:HD23 | 1:A:101:HIS:HB2 | 1.94 | 0.50 |
| 1:A:370:VAL:O | 1:A:371:HIS:ND1 | 2.44 | 0.50 |
| 1:D:174:ALA:HA | 1:D:284:LYS:HD2 | 1.93 | 0.50 |
| 2:Q:1:TRP:O | 2:Q:6:HYP:O | 2.30 | 0.50 |
| 2:S:1:TRP:O | 2:S:6:HYP:O | 2.30 | 0.50 |
| 1:E:151:ILE:HD11 | 1:E:162:ASN:HD21 | 1.75 | 0.50 |
| 1:E:370:VAL:O | 1:E:371:HIS:ND1 | 2.44 | 0.50 |
| 1:A:232:SER:OG | 1:A:232:SER:O | 2.23 | 0.50 |
| 2:N:1:TRP:O | 2:N:6:HYP:O | 2.30 | 0.50 |
| 2:O:1:TRP:O | 2:O:6:HYP:O | 2.30 | 0.50 |
| 2:P:1:TRP:O | 2:P:6:HYP:O | 2.30 | 0.50 |
| 1:D:358:THR:OG1 | 1:D:361:GLU:OE1 | 2.29 | 0.50 |
| 1:E:18:LYS:HG2 | 1:E:30:VAL:HG22 | 1.94 | 0.50 |
| 1:C:358:THR:OG1 | 1:C:361:GLU:OE1 | 2.27 | 0.50 |
| 1:F:35:VAL:HG22 | 1:F:54:VAL:HG13 | 1.93 | 0.50 |
| 2:R:1:TRP:O | 2:R:6:HYP:O | 2.30 | 0.50 |
| 1:D:222:ASP:OD2 | 1:D:225:ASN:ND2 | 2.39 | 0.49 |

Continued on next page...

Not For Manuscript Review

*Continued from previous page...*

| Atom-1 | Atom-2 | Interatomic distance (Å) | Clash overlap (Å) |
|---|---|---|---|
| 2:M:1:TRP:O | 2:M:6:HYP:O | 2.30 | 0.49 |
| 1:A:373:LYS:N | 1:A:373:LYS:HE2 | 2.27 | 0.49 |
| 1:C:154:ASP:OD1 | 1:C:300:SER:OG | 2.30 | 0.49 |
| 1:D:92:ASN:O | 1:D:95:ARG:NH2 | 2.45 | 0.49 |
| 1:E:155:SER:OG | 1:E:303:THR:OG1 | 2.30 | 0.49 |
| 1:E:157:ASP:HB2 | 3:K:376:ADP:H5'1 | 1.93 | 0.49 |
| 1:E:305:MET:SD | 1:E:335:ARG:NH2 | 2.86 | 0.49 |
| 1:D:8:LEU:HD23 | 1:D:101:HIS:HB2 | 1.94 | 0.49 |
| 1:F:16:LEU:O | 1:F:18:LYS:NZ | 2.45 | 0.49 |
| 1:F:18:LYS:HG2 | 1:F:30:VAL:HG22 | 1.94 | 0.49 |
| 1:F:109:PRO:HB3 | 1:F:136:ILE:HD13 | 1.94 | 0.49 |
| 1:F:39:ARG:HB2 | 1:F:66:THR:HG23 | 1.94 | 0.49 |
| 2:S:5:CYS:SG | 2:S:6:HYP:N | 2.85 | 0.49 |
| 1:C:106:THR:OG1 | 1:C:137:GLN:NE2 | 2.35 | 0.49 |
| 1:B:188:TYR:O | 1:B:192:ILE:HG23 | 2.13 | 0.48 |
| 1:C:336:LYS:HE3 | 3:I:376:ADP:C8 | 2.47 | 0.48 |
| 1:F:151:ILE:HD12 | 1:F:164:PRO:HB3 | 1.95 | 0.48 |
| 2:Q:5:CYS:SG | 2:Q:6:HYP:N | 2.85 | 0.48 |
| 1:D:70:PRO:HG3 | 1:D:81:ASP:HB3 | 1.94 | 0.48 |
| 2:O:5:CYS:SG | 2:O:6:HYP:N | 2.85 | 0.48 |
| 1:B:151:ILE:HD11 | 1:B:162:ASN:HD21 | 1.79 | 0.48 |
| 1:C:8:LEU:HD23 | 1:C:101:HIS:HB2 | 1.95 | 0.48 |
| 1:F:190:MET:HG3 | 1:F:209:VAL:HG11 | 1.95 | 0.48 |
| 2:M:5:CYS:SG | 2:M:6:HYP:N | 2.85 | 0.48 |
| 1:B:120:THR:HA | 1:B:132:MET:HE1 | 1.95 | 0.48 |
| 1:C:68:LYS:HB2 | 1:C:68:LYS:HE2 | 1.67 | 0.48 |
| 1:D:250:ILE:HG23 | 1:D:253:GLU:HB2 | 1.96 | 0.48 |
| 1:E:197:GLY:O | 2:R:1:TRP:HE3 | 1.94 | 0.48 |
| 1:C:16:LEU:O | 1:C:18:LYS:NZ | 2.46 | 0.48 |
| 1:D:44:MET:SD | 1:F:170:ALA:HB2 | 2.53 | 0.48 |
| 1:D:157:ASP:N | 3:J:376:ADP:O2B | 2.42 | 0.48 |
| 1:B:35:VAL:HG22 | 1:B:54:VAL:HG13 | 1.96 | 0.47 |
| 1:B:70:PRO:HG2 | 1:B:85:ILE:HD12 | 1.96 | 0.47 |
| 1:B:76:ILE:HG21 | 1:B:82:MET:HB2 | 1.95 | 0.47 |
| 1:E:195:GLU:HB3 | 1:F:113:LYS:HD3 | 1.97 | 0.47 |
| 1:F:260:THR:HG23 | 1:F:266:PHE:HB2 | 1.96 | 0.47 |
| 1:B:189:LEU:HD22 | 1:B:213:LYS:HB2 | 1.97 | 0.47 |
| 1:D:368:SER:O | 1:D:368:SER:OG | 2.27 | 0.47 |
| 1:B:186:THR:HG23 | 1:B:209:VAL:HG12 | 1.96 | 0.47 |
| 1:B:305:MET:SD | 1:B:335:ARG:NH2 | 2.88 | 0.47 |
| 1:C:336:LYS:HE3 | 3:I:376:ADP:H8 | 1.78 | 0.47 |

*Continued on next page...*

Not For Manuscript Review

*Continued from previous page...*

| Atom-1 | Atom-2 | Interatomic distance (Å) | Clash overlap (Å) |
|---|---|---|---|
| 1:D:140:LEU:O | 1:D:342:GLY:HA3 | 2.15 | 0.47 |
| 1:E:71:ILE:HG12 | 1:E:76:ILE:HG12 | 1.95 | 0.47 |
| 1:E:92:ASN:O | 1:E:95:ARG:NH2 | 2.48 | 0.47 |
| 1:F:109:PRO:HD3 | 1:F:136:ILE:HG23 | 1.97 | 0.47 |
| 1:F:111:ASN:OD1 | 1:F:177:ARG:NH1 | 2.42 | 0.47 |
| 1:A:305:MET:SD | 1:A:335:ARG:NH2 | 2.88 | 0.47 |
| 1:F:34:ILE:HD13 | 1:F:67:LEU:HD13 | 1.97 | 0.47 |
| 1:F:357:ILE:HD11 | 1:F:373:LYS:HG3 | 1.97 | 0.47 |
| 1:B:287:ILE:HD12 | 2:M:4:DTH:HG23 | 1.97 | 0.47 |
| 1:F:232:SER:O | 1:F:232:SER:OG | 2.23 | 0.47 |
| 1:F:358:THR:OG1 | 1:F:361:GLU:OE1 | 2.28 | 0.47 |
| 1:A:180:LEU:HD21 | 1:A:260:THR:HG22 | 1.98 | 0.46 |
| 2:N:5:CYS:SG | 2:N:6:HYP:N | 2.85 | 0.46 |
| 1:B:358:THR:OG1 | 1:B:361:GLU:OE1 | 2.28 | 0.46 |
| 1:C:227:MET:HG2 | 1:C:255:PHE:HE2 | 1.80 | 0.46 |
| 1:A:41:GLN:HG2 | 1:A:42:GLY:H | 1.80 | 0.46 |
| 1:B:157:ASP:N | 3:H:376:ADP:O2B | 2.42 | 0.46 |
| 1:E:8:LEU:HD23 | 1:E:101:HIS:HB2 | 1.98 | 0.46 |
| 1:C:92:ASN:O | 1:C:95:ARG:NH2 | 2.48 | 0.46 |
| 1:F:76:ILE:HG12 | 1:F:82:MET:HG2 | 1.97 | 0.46 |
| 1:A:123:MET:HA | 1:A:127:PHE:HD2 | 1.81 | 0.46 |
| 1:C:305:MET:SD | 1:C:335:ARG:NH2 | 2.88 | 0.46 |
| 1:B:259:GLU:OE2 | 1:B:312:ARG:NH1 | 2.44 | 0.46 |
| 2:P:5:CYS:SG | 2:P:6:HYP:N | 2.85 | 0.46 |
| 1:A:117:GLU:OE2 | 1:A:371:HIS:NE2 | 2.47 | 0.45 |
| 1:D:149:THR:HB | 1:D:167:GLU:H | 1.81 | 0.45 |
| 1:A:289:ILE:HG22 | 1:A:293:LEU:HG | 1.98 | 0.45 |
| 1:B:106:THR:OG1 | 1:B:137:GLN:NE2 | 2.48 | 0.45 |
| 1:D:151:ILE:CD1 | 1:D:282:ILE:HG13 | 2.46 | 0.45 |
| 1:A:193:LEU:HB3 | 1:A:198:TYR:O | 2.16 | 0.45 |
| 1:A:244:ASP:HB3 | 1:C:287:ILE:HG13 | 1.98 | 0.45 |
| 1:B:117:GLU:OE2 | 1:B:371:HIS:NE2 | 2.45 | 0.45 |
| 1:F:140:LEU:O | 1:F:342:GLY:HA3 | 2.15 | 0.45 |
| 1:C:105:LEU:HD11 | 1:C:123:MET:SD | 2.56 | 0.45 |
| 1:F:8:LEU:HD23 | 1:F:101:HIS:HB2 | 1.98 | 0.45 |
| 2:R:1:TRP:HB2 | 2:R:4:DTH:O | 2.17 | 0.45 |
| 1:E:73:HIC:HE1 | 1:E:184:ASP:OD1 | 2.16 | 0.45 |
| 2:R:5:CYS:SG | 2:R:6:HYP:N | 2.85 | 0.45 |
| 1:D:120:THR:HG23 | 1:D:132:MET:HE1 | 1.99 | 0.45 |
| 2:N:1:TRP:HB2 | 2:N:4:DTH:O | 2.17 | 0.45 |
| 1:A:70:PRO:HG2 | 1:A:85:ILE:HD13 | 1.98 | 0.45 |

*Continued on next page...*

Not For Manuscript Review

Continued from previous page...

| Atom-1 | Atom-2 | Interatomic distance (Å) | Clash overlap (Å) |
|---|---|---|---|
| 1:A:151:ILE:HD11 | 1:A:162:ASN:HD21 | 1.81 | 0.45 |
| 1:E:360:GLN:NE2 | 1:E:364:GLU:OE2 | 2.50 | 0.45 |
| 1:F:68:LYS:HB2 | 1:F:68:LYS:HE2 | 1.67 | 0.45 |
| 1:A:44:MET:N | 1:A:44:MET:SD | 2.91 | 0.44 |
| 1:D:40:HIS:HB3 | 1:F:170:ALA:O | 2.17 | 0.44 |
| 1:F:157:ASP:N | 3:L:376:ADP:O2B | 2.45 | 0.44 |
| 1:A:120:THR:O | 1:A:124:PHE:HB2 | 2.18 | 0.44 |
| 2:M:1:TRP:HB2 | 2:M:4:DTH:O | 2.17 | 0.44 |
| 2:P:1:TRP:HB2 | 2:P:4:DTH:O | 2.17 | 0.44 |
| 1:E:140:LEU:O | 1:E:342:GLY:HA3 | 2.17 | 0.44 |
| 1:F:178:LEU:HD13 | 1:F:277:THR:HG21 | 2.00 | 0.44 |
| 2:O:1:TRP:HB2 | 2:O:4:DTH:O | 2.17 | 0.44 |
| 2:S:1:TRP:HB2 | 2:S:4:DTH:O | 2.17 | 0.44 |
| 1:D:142:LEU:HD23 | 1:D:165:ILE:HD12 | 1.99 | 0.44 |
| 1:D:193:LEU:HB3 | 1:D:198:TYR:O | 2.17 | 0.44 |
| 1:D:197:GLY:O | 2:Q:1:TRP:CE3 | 2.71 | 0.44 |
| 2:Q:1:TRP:HB2 | 2:Q:4:DTH:O | 2.17 | 0.44 |
| 1:B:70:PRO:HG3 | 1:B:81:ASP:HB3 | 1.99 | 0.44 |
| 1:E:188:TYR:O | 1:E:192:ILE:HG23 | 2.18 | 0.44 |
| 1:A:78:ASN:ND2 | 1:A:81:ASP:OD2 | 2.51 | 0.43 |
| 1:D:18:LYS:HG2 | 1:D:30:VAL:HG22 | 2.00 | 0.43 |
| 1:A:180:LEU:HD12 | 1:A:180:LEU:HA | 1.87 | 0.43 |
| 1:B:110:LEU:O | 1:B:177:ARG:NH1 | 2.51 | 0.43 |
| 1:C:8:LEU:HA | 1:C:21:PHE:HA | 2.00 | 0.43 |
| 1:B:68:LYS:HB2 | 1:B:68:LYS:HE2 | 1.65 | 0.43 |
| 1:C:157:ASP:N | 3:I:376:ADP:O2B | 2.47 | 0.43 |
| 1:A:142:LEU:HD12 | 1:A:142:LEU:HA | 1.83 | 0.43 |
| 1:A:157:ASP:N | 3:G:376:ADP:O2B | 2.46 | 0.43 |
| 1:A:178:LEU:HD23 | 1:A:178:LEU:HA | 1.86 | 0.43 |
| 1:C:142:LEU:HD12 | 1:C:142:LEU:HA | 1.85 | 0.43 |
| 1:B:8:LEU:HD23 | 1:B:101:HIS:HB2 | 2.00 | 0.43 |
| 1:B:18:LYS:HG2 | 1:B:30:VAL:HG22 | 2.01 | 0.43 |
| 1:B:53:TYR:CD2 | 1:B:65:LEU:HD21 | 2.53 | 0.43 |
| 1:D:68:LYS:HB2 | 1:D:68:LYS:HE2 | 1.66 | 0.43 |
| 1:A:197:GLY:O | 2:N:1:TRP:CE3 | 2.72 | 0.43 |
| 1:B:157:ASP:OD2 | 3:H:376:ADP:O3' | 2.24 | 0.43 |
| 1:B:244:ASP:HB2 | 1:D:287:ILE:HG13 | 2.01 | 0.43 |
| 1:D:12:ASN:O | 1:D:12:ASN:ND2 | 2.43 | 0.43 |
| 1:E:297:ASN:HB3 | 1:E:329:ILE:HG12 | 2.01 | 0.43 |
| 1:B:142:LEU:HD12 | 1:B:142:LEU:HA | 1.83 | 0.42 |
| 1:C:186:THR:HG23 | 1:C:209:VAL:HG12 | 2.01 | 0.42 |

Continued on next page...

Not For Manuscript Review

Continued from previous page...

| Atom-1 | Atom-2 | Interatomic distance (Å) | Clash overlap (Å) |
|---|---|---|---|
| 1:E:180:LEU:HD12 | 1:E:180:LEU:HA | 1.78 | 0.42 |
| 1:E:358:THR:OG1 | 1:E:361:GLU:OE1 | 2.32 | 0.42 |
| 1:E:368:SER:O | 1:E:368:SER:OG | 2.27 | 0.42 |
| 1:B:294:TYR:CD2 | 1:B:325:MET:HG2 | 2.55 | 0.42 |
| 1:C:111:ASN:OD1 | 1:C:177:ARG:NH1 | 2.42 | 0.42 |
| 1:F:8:LEU:HA | 1:F:21:PHE:HA | 2.01 | 0.42 |
| 1:A:68:LYS:HB2 | 1:A:68:LYS:HE2 | 1.64 | 0.42 |
| 1:C:5:THR:HB | 1:C:6:THR:H | 1.70 | 0.42 |
| 1:E:76:ILE:HD13 | 1:E:82:MET:HG2 | 2.01 | 0.42 |
| 1:B:157:ASP:HB2 | 3:H:376:ADP:H5'1 | 2.02 | 0.42 |
| 1:E:70:PRO:HG3 | 1:E:81:ASP:HB3 | 2.01 | 0.42 |
| 1:E:262:PHE:HE1 | 1:E:313:MET:HG2 | 1.84 | 0.42 |
| 1:F:151:ILE:HD11 | 1:F:162:ASN:OD1 | 2.19 | 0.42 |
| 1:F:373:LYS:HB3 | 1:F:373:LYS:HE3 | 1.87 | 0.42 |
| 1:A:16:LEU:O | 1:A:18:LYS:NZ | 2.52 | 0.42 |
| 1:C:213:LYS:O | 1:C:217:CYS:HB2 | 2.19 | 0.42 |
| 1:F:160:THR:OG1 | 1:F:180:LEU:O | 2.28 | 0.42 |
| 1:B:289:ILE:HG22 | 1:B:293:LEU:HG | 2.02 | 0.42 |
| 1:C:76:ILE:HD12 | 1:C:79:TRP:CH2 | 2.54 | 0.42 |
| 1:D:151:ILE:HD11 | 1:D:162:ASN:ND2 | 2.35 | 0.42 |
| 1:A:155:SER:OG | 1:A:303:THR:OG1 | 2.29 | 0.42 |
| 1:E:213:LYS:O | 1:E:217:CYS:HB2 | 2.20 | 0.42 |
| 1:A:8:LEU:HA | 1:A:21:PHE:HA | 2.02 | 0.41 |
| 1:A:213:LYS:O | 1:A:217:CYS:HB2 | 2.20 | 0.41 |
| 1:C:14:SER:HB3 | 1:C:71:ILE:HG22 | 2.02 | 0.41 |
| 1:D:213:LYS:O | 1:D:217:CYS:HB2 | 2.20 | 0.41 |
| 1:D:106:THR:OG1 | 1:D:137:GLN:NE2 | 2.52 | 0.41 |
| 1:D:287:ILE:CD1 | 2:O:4:DTH:HG23 | 2.48 | 0.41 |
| 1:E:259:GLU:OE2 | 1:E:312:ARG:NH1 | 2.46 | 0.41 |
| 1:B:294:TYR:HD2 | 1:B:325:MET:HG2 | 1.85 | 0.41 |
| 1:C:155:SER:HG | 1:C:303:THR:HG1 | 1.58 | 0.41 |
| 1:C:290:ARG:H | 1:C:290:ARG:HG3 | 1.51 | 0.41 |
| 1:F:36:GLY:HA3 | 1:F:65:LEU:HD23 | 2.02 | 0.41 |
| 1:F:71:ILE:O | 1:F:183:ARG:NH2 | 2.50 | 0.41 |
| 1:C:180:LEU:HD23 | 1:C:180:LEU:HA | 1.90 | 0.41 |
| 1:A:368:SER:O | 1:A:368:SER:OG | 2.32 | 0.41 |
| 1:C:178:LEU:HD23 | 1:C:178:LEU:HA | 1.93 | 0.41 |
| 1:C:117:GLU:OE2 | 1:C:371:HIS:NE2 | 2.48 | 0.41 |
| 1:D:76:ILE:HD12 | 1:D:79:TRP:CH2 | 2.55 | 0.41 |
| 1:A:372:ARG:HG2 | 1:A:373:LYS:HE2 | 2.02 | 0.41 |
| 1:C:16:LEU:HD12 | 1:C:16:LEU:HA | 1.91 | 0.41 |

Continued on next page...

Not For Manuscript Review

*Continued from previous page...*

| Atom-1 | Atom-2 | Interatomic distance (Å) | Clash overlap (Å) |
|---|---|---|---|
| 1:D:227:MET:HG2 | 1:D:255:PHE:HE1 | 1.86 | 0.41 |
| 1:A:18:LYS:HG2 | 1:A:30:VAL:HG22 | 2.03 | 0.40 |
| 1:B:156:GLY:O | 1:B:181:ALA:HB1 | 2.21 | 0.40 |
| 1:B:213:LYS:O | 1:B:217:CYS:HB2 | 2.22 | 0.40 |
| 1:E:117:GLU:OE2 | 1:E:371:HIS:NE2 | 2.54 | 0.40 |
| 1:A:282:ILE:HG21 | 1:A:294:TYR:CE1 | 2.56 | 0.40 |
| 1:C:287:ILE:HA | 1:C:290:ARG:HE | 1.85 | 0.40 |
| 1:F:368:SER:O | 1:F:368:SER:OG | 2.26 | 0.40 |
| 1:B:8:LEU:HA | 1:B:21:PHE:HA | 2.03 | 0.40 |
| 1:B:139:VAL:HG22 | 1:B:165:ILE:HD11 | 2.02 | 0.40 |
| 1:C:70:PRO:HG3 | 1:C:81:ASP:HB3 | 2.03 | 0.40 |

There are no symmetry-related clashes.

#### 5.3 Torsion angles (i)

##### 5.3.1 Protein backbone (i)

In the following table, the Percentiles column shows the percent Ramachandran outliers of the chain as a percentile score with respect to all PDB entries followed by that with respect to all EM entries.

The Analysed column shows the number of residues for which the backbone conformation was analysed, and the total number of residues.

| Mol | Chain | Analysed | Favoured | Allowed | Outliers | Percentiles | |
|---|---|---|---|---|---|---|---|
| 1 | A | 368/371 (99%) | 346 (94%) | 21 (6%) | 1 (0%) | 41 | 41 |
| 1 | B | 368/371 (99%) | 353 (96%) | 14 (4%) | 1 (0%) | 41 | 41 |
| 1 | C | 368/371 (99%) | 354 (96%) | 13 (4%) | 1 (0%) | 41 | 41 |
| 1 | D | 368/371 (99%) | 350 (95%) | 17 (5%) | 1 (0%) | 41 | 41 |
| 1 | E | 368/371 (99%) | 350 (95%) | 17 (5%) | 1 (0%) | 41 | 41 |
| 1 | F | 368/371 (99%) | 353 (96%) | 14 (4%) | 1 (0%) | 41 | 41 |
| 2 | M | 2/7 (29%) | 1 (50%) | 0 | 1 (50%) | 0 | 0 |
| 2 | N | 2/7 (29%) | 1 (50%) | 0 | 1 (50%) | 0 | 0 |
| 2 | O | 2/7 (29%) | 1 (50%) | 0 | 1 (50%) | 0 | 0 |
| 2 | P | 2/7 (29%) | 1 (50%) | 0 | 1 (50%) | 0 | 0 |
| 2 | Q | 2/7 (29%) | 1 (50%) | 0 | 1 (50%) | 0 | 0 |

*Continued on next page...*

Not For Manuscript Review

*Continued from previous page...*

| Mol | Chain | Analysed | Favoured | Allowed | Outliers | Percentiles | |
|---|---|---|---|---|---|---|---|
| 2 | R | 2/7 (29%) | 1 (50%) | 0 | 1 (50%) | 0 | 0 |
| 2 | S | 2/7 (29%) | 1 (50%) | 0 | 1 (50%) | 0 | 0 |
| All | All | 2222/2275 (98%) | 2113 (95%) | 96 (4%) | 13 (1%) | 29 | 25 |

All (13) Ramachandran outliers are listed below:

| Mol | Chain | Res | Type |
|---|---|---|---|
| 2 | M | 5 | CYS |
| 2 | N | 5 | CYS |
| 2 | O | 5 | CYS |
| 2 | P | 5 | CYS |
| 2 | Q | 5 | CYS |
| 2 | R | 5 | CYS |
| 2 | S | 5 | CYS |
| 1 | B | 43 | VAL |
| 1 | C | 43 | VAL |
| 1 | D | 43 | VAL |
| 1 | E | 43 | VAL |
| 1 | F | 43 | VAL |
| 1 | A | 43 | VAL |

##### 5.3.2 Protein sidechains (i)

In the following table, the Percentiles column shows the percent sidechain outliers of the chain as a percentile score with respect to all PDB entries followed by that with respect to all EM entries.

The Analysed column shows the number of residues for which the sidechain conformation was analysed, and the total number of residues.

| Mol | Chain | Analysed | Rotameric | Outliers | Percentiles | |
|---|---|---|---|---|---|---|
| 1 | A | 313/313 (100%) | 304 (97%) | 9 (3%) | 42 | 42 |
| 1 | B | 313/313 (100%) | 304 (97%) | 9 (3%) | 42 | 42 |
| 1 | C | 313/313 (100%) | 306 (98%) | 7 (2%) | 52 | 52 |
| 1 | D | 313/313 (100%) | 302 (96%) | 11 (4%) | 36 | 36 |
| 1 | E | 313/313 (100%) | 303 (97%) | 10 (3%) | 39 | 39 |
| 1 | F | 313/313 (100%) | 306 (98%) | 7 (2%) | 52 | 52 |
| 2 | M | 2/2 (100%) | 1 (50%) | 1 (50%) | 0 | 0 |

*Continued on next page...*

Not For Manuscript Review

*Continued from previous page...*

| Mol | Chain | Analysed | Rotameric | Outliers | Percentiles | |
|---|---|---|---|---|---|---|
| 2 | N | 2/2 (100%) | 1 (50%) | 1 (50%) | 0 | 0 |
| 2 | O | 2/2 (100%) | 1 (50%) | 1 (50%) | 0 | 0 |
| 2 | P | 2/2 (100%) | 1 (50%) | 1 (50%) | 0 | 0 |
| 2 | Q | 2/2 (100%) | 1 (50%) | 1 (50%) | 0 | 0 |
| 2 | R | 2/2 (100%) | 1 (50%) | 1 (50%) | 0 | 0 |
| 2 | S | 2/2 (100%) | 1 (50%) | 1 (50%) | 0 | 0 |
| All | All | 1892/1892 (100%) | 1832 (97%) | 60 (3%) | 42 | 39 |

All (60) residues with a non-rotameric sidechain are listed below:

| Mol | Chain | Res | Type |
|---|---|---|---|
| 1 | A | 25 | ASP |
| 1 | A | 111 | ASN |
| 1 | A | 124 | PHE |
| 1 | A | 190 | MET |
| 1 | A | 199 | SER |
| 1 | A | 232 | SER |
| 1 | A | 297 | ASN |
| 1 | A | 300 | SER |
| 1 | A | 354 | GLN |
| 1 | B | 56 | ASP |
| 1 | B | 111 | ASN |
| 1 | B | 124 | PHE |
| 1 | B | 145 | SER |
| 1 | B | 198 | TYR |
| 1 | B | 232 | SER |
| 1 | B | 276 | GLU |
| 1 | B | 286 | ASP |
| 1 | B | 354 | GLN |
| 1 | C | 56 | ASP |
| 1 | C | 111 | ASN |
| 1 | C | 132 | MET |
| 1 | C | 283 | MET |
| 1 | C | 313 | MET |
| 1 | C | 326 | LYS |
| 1 | C | 354 | GLN |
| 1 | D | 12 | ASN |
| 1 | D | 56 | ASP |
| 1 | D | 124 | PHE |
| 1 | D | 145 | SER |

*Continued on next page...*

Not For Manuscript Review

*Continued from previous page...*

| Mol | Chain | Res | Type |
|---|---|---|---|
| 1 | D | 198 | TYR |
| 1 | D | 232 | SER |
| 1 | D | 257 | CYS |
| 1 | D | 265 | SER |
| 1 | D | 326 | LYS |
| 1 | D | 354 | GLN |
| 1 | D | 364 | GLU |
| 1 | E | 111 | ASN |
| 1 | E | 124 | PHE |
| 1 | E | 132 | MET |
| 1 | E | 143 | TYR |
| 1 | E | 178 | LEU |
| 1 | E | 232 | SER |
| 1 | E | 265 | SER |
| 1 | E | 276 | GLU |
| 1 | E | 286 | ASP |
| 1 | E | 288 | ASP |
| 1 | F | 12 | ASN |
| 1 | F | 111 | ASN |
| 1 | F | 124 | PHE |
| 1 | F | 143 | TYR |
| 1 | F | 232 | SER |
| 1 | F | 374 | CYS |
| 1 | F | 375 | PHE |
| 2 | M | 5 | CYS |
| 2 | N | 5 | CYS |
| 2 | O | 5 | CYS |
| 2 | P | 5 | CYS |
| 2 | Q | 5 | CYS |
| 2 | R | 5 | CYS |
| 2 | S | 5 | CYS |

Sometimes sidechains can be flipped to improve hydrogen bonding and reduce clashes. All (1) such sidechains are listed below:

| Mol | Chain | Res | Type |
|---|---|---|---|
| 1 | F | 12 | ASN |

##### 5.3.3 RNA ⓘ

There are no RNA molecules in this entry.

Not For Manuscript Review

#### 5.4 Non-standard residues in protein, DNA, RNA chains (i)

27 non-standard protein/DNA/RNA residues are modelled in this entry.

In the following table, the Counts columns list the number of bonds (or angles) for which Mogul statistics could be retrieved, the number of bonds (or angles) that are observed in the model and the number of bonds (or angles) that are defined in the Chemical Component Dictionary. The Link column lists molecule types, if any, to which the group is linked. The Z score for a bond length (or angle) is the number of standard deviations the observed value is removed from the expected value. A bond length (or angle) with $|Z| > 2$ is considered an outlier worth inspection. RMSZ is the root-mean-square of all Z scores of the bond lengths (or angles).

| Mol | Type | Chain | Res | Link | Bond lengths | | | Bond angles | | |
|---|---|---|---|---|---|---|---|---|---|---|
| | | | | | Counts | RMSZ | $\#\|Z\| > 2$ | Counts | RMSZ | $\#\|Z\| > 2$ |
| 2 | EEP | O | 2 | 2 | 7,9,10 | 1.14 | 1 (14%) | 3,12,14 | 1.02 | 0 |
| 2 | EEP | Q | 2 | 2 | 7,9,10 | 1.15 | 1 (14%) | 3,12,14 | 1.01 | 0 |
| 1 | HIC | A | 73 | 1 | 8,11,12 | 1.64 | 2 (25%) | 5,14,16 | 0.93 | 0 |
| 2 | EEP | R | 2 | 2 | 7,9,10 | 1.17 | 1 (14%) | 3,12,14 | 1.01 | 0 |
| 2 | EEP | S | 2 | 2 | 7,9,10 | 1.14 | 1 (14%) | 3,12,14 | 1.00 | 0 |
| 2 | HYP | Q | 6 | 2 | 7,8,9 | 0.56 | 0 | 5,10,12 | 1.35 | 1 (20%) |
| 1 | HIC | C | 73 | 1 | 8,11,12 | 1.66 | 2 (25%) | 5,14,16 | 0.99 | 0 |
| 1 | HIC | F | 73 | 1 | 8,11,12 | 1.65 | 2 (25%) | 5,14,16 | 0.90 | 0 |
| 2 | EEP | N | 2 | 2 | 7,9,10 | 1.15 | 1 (14%) | 3,12,14 | 1.02 | 0 |
| 2 | HYP | M | 6 | 2 | 7,8,9 | 0.56 | 0 | 5,10,12 | 1.36 | 1 (20%) |
| 2 | HYP | N | 6 | 2 | 7,8,9 | 0.57 | 0 | 5,10,12 | 1.34 | 1 (20%) |
| 2 | EEP | M | 2 | 2 | 7,9,10 | 1.15 | 1 (14%) | 3,12,14 | 1.01 | 0 |
| 1 | HIC | E | 73 | 1 | 8,11,12 | 1.65 | 2 (25%) | 5,14,16 | 0.97 | 0 |
| 1 | HIC | B | 73 | 1 | 8,11,12 | 1.65 | 2 (25%) | 5,14,16 | 0.89 | 0 |
| 2 | HYP | P | 6 | 2 | 7,8,9 | 0.56 | 0 | 5,10,12 | 1.35 | 1 (20%) |
| 2 | HYP | O | 6 | 2 | 7,8,9 | 0.56 | 0 | 5,10,12 | 1.35 | 1 (20%) |
| 2 | HYP | S | 6 | 2 | 7,8,9 | 0.58 | 0 | 5,10,12 | 1.34 | 1 (20%) |
| 2 | HYP | R | 6 | 2 | 7,8,9 | 0.57 | 0 | 5,10,12 | 1.35 | 1 (20%) |
| 1 | HIC | D | 73 | 1 | 8,11,12 | 1.65 | 2 (25%) | 5,14,16 | 0.96 | 0 |
| 2 | EEP | P | 2 | 2 | 7,9,10 | 1.16 | 1 (14%) | 3,12,14 | 1.01 | 0 |

In the following table, the Chirals column lists the number of chiral outliers, the number of chiral centers analysed, the number of these observed in the model and the number defined in the Chemical Component Dictionary. Similar counts are reported in the Torsion and Rings columns. '-' means no outliers of that kind were identified.

Not For Manuscript Review

| Mol | Type | Chain | Res | Link | Chirals | Torsions | Rings |
|---|---|---|---|---|---|---|---|
| 2 | EEP | O | 2 | 2 | - | 4/9/10/12 | - |
| 2 | EEP | Q | 2 | 2 | - | 4/9/10/12 | - |
| 1 | HIC | A | 73 | 1 | - | 2/5/6/8 | 0/1/1/1 |
| 2 | EEP | R | 2 | 2 | - | 4/9/10/12 | - |
| 2 | EEP | S | 2 | 2 | - | 4/9/10/12 | - |
| 2 | HYP | Q | 6 | 2 | - | 0/0/11/13 | 0/1/1/1 |
| 1 | HIC | C | 73 | 1 | - | 2/5/6/8 | 0/1/1/1 |
| 1 | HIC | F | 73 | 1 | - | 2/5/6/8 | 0/1/1/1 |
| 2 | EEP | N | 2 | 2 | - | 4/9/10/12 | - |
| 2 | HYP | M | 6 | 2 | - | 0/0/11/13 | 0/1/1/1 |
| 2 | HYP | N | 6 | 2 | - | 0/0/11/13 | 0/1/1/1 |
| 2 | EEP | M | 2 | 2 | - | 4/9/10/12 | - |
| 1 | HIC | E | 73 | 1 | - | 2/5/6/8 | 0/1/1/1 |
| 1 | HIC | B | 73 | 1 | - | 2/5/6/8 | 0/1/1/1 |
| 2 | HYP | P | 6 | 2 | - | 0/0/11/13 | 0/1/1/1 |
| 2 | HYP | O | 6 | 2 | - | 0/0/11/13 | 0/1/1/1 |
| 2 | HYP | S | 6 | 2 | - | 0/0/11/13 | 0/1/1/1 |
| 2 | HYP | R | 6 | 2 | - | 0/0/11/13 | 0/1/1/1 |
| 1 | HIC | D | 73 | 1 | - | 4/5/6/8 | 0/1/1/1 |
| 2 | EEP | P | 2 | 2 | - | 4/9/10/12 | - |

All (19) bond length outliers are listed below:

| Mol | Chain | Res | Type | Atoms | Z | Observed(Å) | Ideal(Å) |
|---|---|---|---|---|---|---|---|
| 1 | C | 73 | HIC | CD2-CG | 3.68 | 1.41 | 1.36 |
| 1 | F | 73 | HIC | CD2-CG | 3.60 | 1.41 | 1.36 |
| 1 | B | 73 | HIC | CD2-CG | 3.59 | 1.41 | 1.36 |
| 1 | E | 73 | HIC | CD2-CG | 3.59 | 1.41 | 1.36 |
| 1 | A | 73 | HIC | CD2-CG | 3.56 | 1.41 | 1.36 |
| 1 | D | 73 | HIC | CD2-CG | 3.55 | 1.41 | 1.36 |
| 2 | R | 2 | EEP | CB-CA | -2.62 | 1.50 | 1.54 |
| 2 | P | 2 | EEP | CB-CA | -2.61 | 1.50 | 1.54 |
| 2 | N | 2 | EEP | CB-CA | -2.60 | 1.50 | 1.54 |
| 2 | Q | 2 | EEP | CB-CA | -2.59 | 1.50 | 1.54 |
| 2 | O | 2 | EEP | CB-CA | -2.55 | 1.50 | 1.54 |
| 2 | M | 2 | EEP | CB-CA | -2.55 | 1.50 | 1.54 |
| 2 | S | 2 | EEP | CB-CA | -2.55 | 1.50 | 1.54 |
| 1 | D | 73 | HIC | CZ-NE2 | -2.05 | 1.42 | 1.48 |
| 1 | E | 73 | HIC | CZ-NE2 | -2.04 | 1.42 | 1.48 |
| 1 | B | 73 | HIC | CZ-NE2 | -2.04 | 1.42 | 1.48 |

*Continued on next page...*

Not For Manuscript Review

*Continued from previous page...*

| Mol | Chain | Res | Type | Atoms | Z | Observed(Å) | Ideal(Å) |
|---|---|---|---|---|---|---|---|
| 1 | C | 73 | HIC | CZ-NE2 | -2.04 | 1.42 | 1.48 |
| 1 | F | 73 | HIC | CZ-NE2 | -2.03 | 1.42 | 1.48 |
| 1 | A | 73 | HIC | CZ-NE2 | -2.03 | 1.42 | 1.48 |

All (7) bond angle outliers are listed below:

| Mol | Chain | Res | Type | Atoms | Z | Observed($^o$) | Ideal($^o$) |
|---|---|---|---|---|---|---|---|
| 2 | P | 6 | HYP | O-C-CA | -2.35 | 118.72 | 124.77 |
| 2 | M | 6 | HYP | O-C-CA | -2.35 | 118.72 | 124.77 |
| 2 | Q | 6 | HYP | O-C-CA | -2.34 | 118.74 | 124.77 |
| 2 | N | 6 | HYP | O-C-CA | -2.33 | 118.77 | 124.77 |
| 2 | R | 6 | HYP | O-C-CA | -2.33 | 118.78 | 124.77 |
| 2 | O | 6 | HYP | O-C-CA | -2.31 | 118.82 | 124.77 |
| 2 | S | 6 | HYP | O-C-CA | -2.30 | 118.85 | 124.77 |

There are no chirality outliers.

All (42) torsion outliers are listed below:

| Mol | Chain | Res | Type | Atoms |
|---|---|---|---|---|
| 1 | A | 73 | HIC | CA-CB-CG-ND1 |
| 1 | A | 73 | HIC | CA-CB-CG-CD2 |
| 1 | B | 73 | HIC | CA-CB-CG-ND1 |
| 1 | B | 73 | HIC | CA-CB-CG-CD2 |
| 1 | C | 73 | HIC | CA-CB-CG-ND1 |
| 1 | C | 73 | HIC | CA-CB-CG-CD2 |
| 1 | D | 73 | HIC | C-CA-CB-CG |
| 1 | D | 73 | HIC | CA-CB-CG-ND1 |
| 1 | D | 73 | HIC | CA-CB-CG-CD2 |
| 1 | E | 73 | HIC | CA-CB-CG-ND1 |
| 1 | E | 73 | HIC | CA-CB-CG-CD2 |
| 1 | F | 73 | HIC | CA-CB-CG-ND1 |
| 1 | F | 73 | HIC | CA-CB-CG-CD2 |
| 2 | M | 2 | EEP | O2-CD1-CG-CB |
| 2 | M | 2 | EEP | O2-CD1-CG-O1 |
| 2 | M | 2 | EEP | O2-CD1-CG-CD2 |
| 2 | N | 2 | EEP | O2-CD1-CG-CB |
| 2 | N | 2 | EEP | O2-CD1-CG-O1 |
| 2 | N | 2 | EEP | O2-CD1-CG-CD2 |
| 2 | O | 2 | EEP | O2-CD1-CG-CB |
| 2 | O | 2 | EEP | O2-CD1-CG-O1 |
| 2 | O | 2 | EEP | O2-CD1-CG-CD2 |
| 2 | P | 2 | EEP | O2-CD1-CG-CB |

*Continued on next page...*

Not For Manuscript Review

*Continued from previous page...*

| Mol | Chain | Res | Type | Atoms |
|---|---|---|---|---|
| 2 | P | 2 | EEP | O2-CD1-CG-O1 |
| 2 | P | 2 | EEP | O2-CD1-CG-CD2 |
| 2 | Q | 2 | EEP | O2-CD1-CG-CB |
| 2 | Q | 2 | EEP | O2-CD1-CG-O1 |
| 2 | Q | 2 | EEP | O2-CD1-CG-CD2 |
| 2 | R | 2 | EEP | O2-CD1-CG-CB |
| 2 | R | 2 | EEP | O2-CD1-CG-O1 |
| 2 | R | 2 | EEP | O2-CD1-CG-CD2 |
| 2 | S | 2 | EEP | O2-CD1-CG-CB |
| 2 | S | 2 | EEP | O2-CD1-CG-O1 |
| 2 | S | 2 | EEP | O2-CD1-CG-CD2 |
| 1 | D | 73 | HIC | N-CA-CB-CG |
| 2 | M | 2 | EEP | C-CA-CB-CG |
| 2 | N | 2 | EEP | C-CA-CB-CG |
| 2 | O | 2 | EEP | C-CA-CB-CG |
| 2 | P | 2 | EEP | C-CA-CB-CG |
| 2 | Q | 2 | EEP | C-CA-CB-CG |
| 2 | R | 2 | EEP | C-CA-CB-CG |
| 2 | S | 2 | EEP | C-CA-CB-CG |

There are no ring outliers.

15 monomers are involved in 36 short contacts:

| Mol | Chain | Res | Type | Clashes | Symm-Clashes |
|---|---|---|---|---|---|
| 2 | O | 2 | EEP | 1 | 0 |
| 2 | Q | 2 | EEP | 1 | 0 |
| 2 | R | 2 | EEP | 1 | 0 |
| 2 | S | 2 | EEP | 1 | 0 |
| 2 | Q | 6 | HYP | 4 | 0 |
| 2 | N | 2 | EEP | 1 | 0 |
| 2 | M | 6 | HYP | 4 | 0 |
| 2 | N | 6 | HYP | 4 | 0 |
| 2 | M | 2 | EEP | 1 | 0 |
| 1 | E | 73 | HIC | 1 | 0 |
| 2 | P | 6 | HYP | 4 | 0 |
| 2 | O | 6 | HYP | 4 | 0 |
| 2 | S | 6 | HYP | 4 | 0 |
| 2 | R | 6 | HYP | 4 | 0 |
| 2 | P | 2 | EEP | 1 | 0 |

Not For Manuscript Review

#### 5.5 Carbohydrates (i)

There are no monosaccharides in this entry.

#### 5.6 Ligand geometry (i)

Of 12 ligands modelled in this entry, 6 are monoatomic - leaving 6 for Mogul analysis.

In the following table, the Counts columns list the number of bonds (or angles) for which Mogul statistics could be retrieved, the number of bonds (or angles) that are observed in the model and the number of bonds (or angles) that are defined in the Chemical Component Dictionary. The Link column lists molecule types, if any, to which the group is linked. The Z score for a bond length (or angle) is the number of standard deviations the observed value is removed from the expected value. A bond length (or angle) with $|Z| > 2$ is considered an outlier worth inspection. RMSZ is the root-mean-square of all Z scores of the bond lengths (or angles).

| Mol | Type | Chain | Res | Link | Bond lengths | | | Bond angles | | |
|---|---|---|---|---|---|---|---|---|---|---|
| | | | | | Counts | RMSZ | $\#\|Z\| > 2$ | Counts | RMSZ | $\#\|Z\| > 2$ |
| 3 | ADP | G | 376 | 4 | 24,29,29 | 0.89 | 0 | 29,45,45 | 1.18 | 2 (6%) |
| 3 | ADP | I | 376 | 4 | 24,29,29 | 0.90 | 0 | 29,45,45 | 1.14 | 2 (6%) |
| 3 | ADP | L | 376 | 4 | 24,29,29 | 0.89 | 0 | 29,45,45 | 1.16 | 2 (6%) |
| 3 | ADP | K | 376 | 4 | 24,29,29 | 0.87 | 0 | 29,45,45 | 1.17 | 2 (6%) |
| 3 | ADP | H | 376 | 4 | 24,29,29 | 0.88 | 0 | 29,45,45 | 1.16 | 2 (6%) |
| 3 | ADP | J | 376 | 4 | 24,29,29 | 0.89 | 0 | 29,45,45 | 1.17 | 2 (6%) |

In the following table, the Chirals column lists the number of chiral outliers, the number of chiral centers analysed, the number of these observed in the model and the number defined in the Chemical Component Dictionary. Similar counts are reported in the Torsion and Rings columns. '-' means no outliers of that kind were identified.

| Mol | Type | Chain | Res | Link | Chirals | Torsions | Rings |
|---|---|---|---|---|---|---|---|
| 3 | ADP | G | 376 | 4 | - | 6/12/32/32 | 0/3/3/3 |
| 3 | ADP | I | 376 | 4 | - | 7/12/32/32 | 0/3/3/3 |
| 3 | ADP | L | 376 | 4 | - | 5/12/32/32 | 0/3/3/3 |
| 3 | ADP | K | 376 | 4 | - | 7/12/32/32 | 0/3/3/3 |
| 3 | ADP | H | 376 | 4 | - | 7/12/32/32 | 0/3/3/3 |
| 3 | ADP | J | 376 | 4 | - | 7/12/32/32 | 0/3/3/3 |

There are no bond length outliers.

All (12) bond angle outliers are listed below:

Not For Manuscript Review

| Mol | Chain | Res | Type | Atoms | Z | Observed(°) | Ideal(°) |
|---|---|---|---|---|---|---|---|
| 3 | G | 376 | ADP | N3-C2-N1 | -3.59 | 123.81 | 128.67 |
| 3 | J | 376 | ADP | N3-C2-N1 | -3.56 | 123.84 | 128.67 |
| 3 | K | 376 | ADP | N3-C2-N1 | -3.55 | 123.86 | 128.67 |
| 3 | H | 376 | ADP | N3-C2-N1 | -3.54 | 123.87 | 128.67 |
| 3 | L | 376 | ADP | N3-C2-N1 | -3.50 | 123.93 | 128.67 |
| 3 | I | 376 | ADP | N3-C2-N1 | -3.47 | 123.96 | 128.67 |
| 3 | K | 376 | ADP | C4-C5-N7 | -2.62 | 106.56 | 109.34 |
| 3 | J | 376 | ADP | C4-C5-N7 | -2.60 | 106.58 | 109.34 |
| 3 | H | 376 | ADP | C4-C5-N7 | -2.58 | 106.62 | 109.34 |
| 3 | G | 376 | ADP | C4-C5-N7 | -2.56 | 106.63 | 109.34 |
| 3 | I | 376 | ADP | C4-C5-N7 | -2.56 | 106.63 | 109.34 |
| 3 | L | 376 | ADP | C4-C5-N7 | -2.55 | 106.65 | 109.34 |

There are no chirality outliers.

All (39) torsion outliers are listed below:

| Mol | Chain | Res | Type | Atoms |
|---|---|---|---|---|
| 3 | G | 376 | ADP | C5'-O5'-PA-O3A |
| 3 | G | 376 | ADP | O4'-C4'-C5'-O5' |
| 3 | H | 376 | ADP | C5'-O5'-PA-O1A |
| 3 | H | 376 | ADP | C5'-O5'-PA-O2A |
| 3 | H | 376 | ADP | O4'-C4'-C5'-O5' |
| 3 | H | 376 | ADP | C3'-C4'-C5'-O5' |
| 3 | I | 376 | ADP | C5'-O5'-PA-O2A |
| 3 | I | 376 | ADP | O4'-C4'-C5'-O5' |
| 3 | J | 376 | ADP | C5'-O5'-PA-O2A |
| 3 | J | 376 | ADP | O4'-C4'-C5'-O5' |
| 3 | K | 376 | ADP | C5'-O5'-PA-O2A |
| 3 | L | 376 | ADP | O4'-C4'-C5'-O5' |
| 3 | J | 376 | ADP | C3'-C4'-C5'-O5' |
| 3 | I | 376 | ADP | C3'-C4'-C5'-O5' |
| 3 | G | 376 | ADP | C3'-C4'-C5'-O5' |
| 3 | L | 376 | ADP | C3'-C4'-C5'-O5' |
| 3 | G | 376 | ADP | PB-O3A-PA-O2A |
| 3 | I | 376 | ADP | PB-O3A-PA-O2A |
| 3 | J | 376 | ADP | PB-O3A-PA-O2A |
| 3 | K | 376 | ADP | PB-O3A-PA-O1A |
| 3 | L | 376 | ADP | PB-O3A-PA-O2A |
| 3 | G | 376 | ADP | C5'-O5'-PA-O1A |
| 3 | H | 376 | ADP | C5'-O5'-PA-O3A |
| 3 | I | 376 | ADP | C5'-O5'-PA-O1A |
| 3 | I | 376 | ADP | C5'-O5'-PA-O3A |
| 3 | J | 376 | ADP | C5'-O5'-PA-O1A |

*Continued on next page...*

Not For Manuscript Review

*Continued from previous page...*

| Mol | Chain | Res | Type | Atoms |
|---|---|---|---|---|
| 3 | J | 376 | ADP | C5'-O5'-PA-O3A |
| 3 | K | 376 | ADP | C5'-O5'-PA-O1A |
| 3 | K | 376 | ADP | C5'-O5'-PA-O3A |
| 3 | L | 376 | ADP | C5'-O5'-PA-O3A |
| 3 | H | 376 | ADP | PB-O3A-PA-O1A |
| 3 | H | 376 | ADP | PB-O3A-PA-O2A |
| 3 | K | 376 | ADP | PB-O3A-PA-O2A |
| 3 | K | 376 | ADP | O4'-C4'-C5'-O5' |
| 3 | I | 376 | ADP | PB-O3A-PA-O1A |
| 3 | K | 376 | ADP | C3'-C4'-C5'-O5' |
| 3 | J | 376 | ADP | PB-O3A-PA-O1A |
| 3 | G | 376 | ADP | PB-O3A-PA-O1A |
| 3 | L | 376 | ADP | PB-O3A-PA-O1A |

There are no ring outliers.

6 monomers are involved in 12 short contacts:

| Mol | Chain | Res | Type | Clashes | Symm-Clashes |
|---|---|---|---|---|---|
| 3 | G | 376 | ADP | 1 | 0 |
| 3 | I | 376 | ADP | 3 | 0 |
| 3 | L | 376 | ADP | 1 | 0 |
| 3 | K | 376 | ADP | 3 | 0 |
| 3 | H | 376 | ADP | 3 | 0 |
| 3 | J | 376 | ADP | 1 | 0 |

The following is a two-dimensional graphical depiction of Mogul quality analysis of bond lengths, bond angles, torsion angles, and ring geometry for all instances of the Ligand of Interest. In addition, ligands with molecular weight > 250 and outliers as shown on the validation Tables will also be included. For torsion angles, if less then 5% of the Mogul distribution of torsion angles is within 10 degrees of the torsion angle in question, then that torsion angle is considered an outlier. Any bond that is central to one or more torsion angles identified as an outlier by Mogul will be highlighted in the graph. For rings, the root-mean-square deviation (RMSD) between the ring in question and similar rings identified by Mogul is calculated over all ring torsion angles. If the average RMSD is greater than 60 degrees and the minimal RMSD between the ring in question and any Mogul-identified rings is also greater than 60 degrees, then that ring is considered an outlier. The outliers are highlighted in purple. The color gray indicates Mogul did not find sufficient equivalents in the CSD to analyse the geometry.

Not For Manuscript Review

##### Ligand ADP G 376

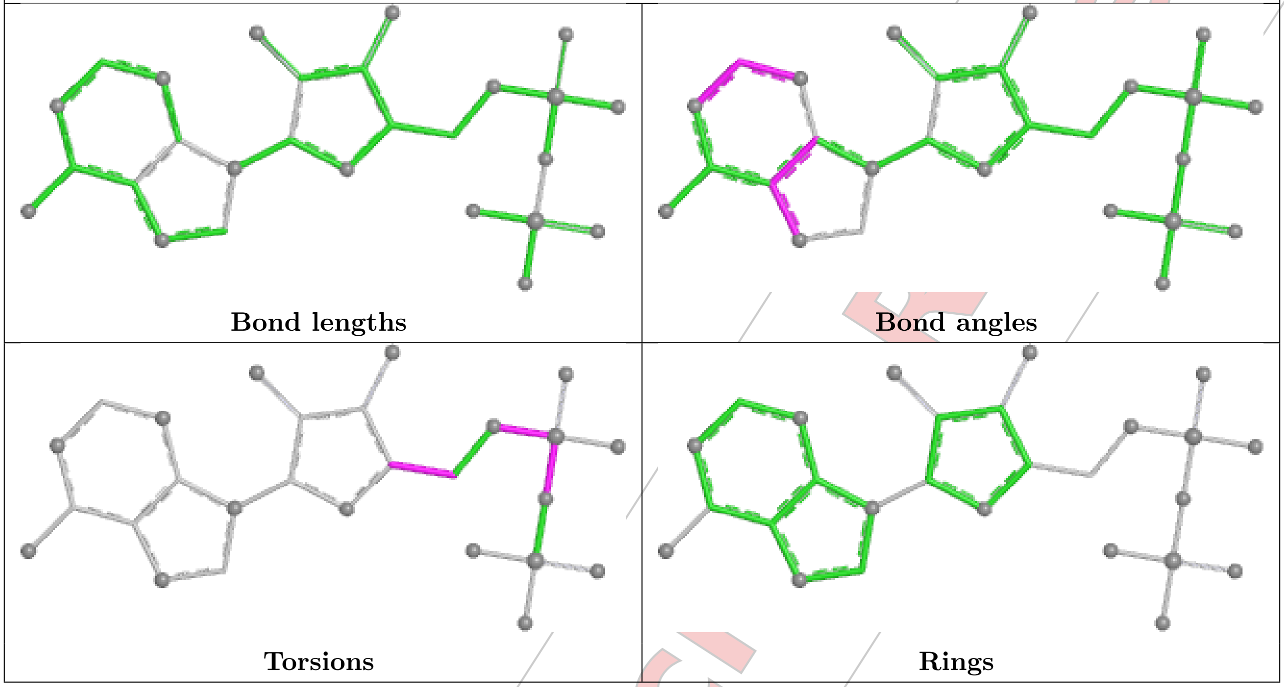


##### Ligand ADP I 376

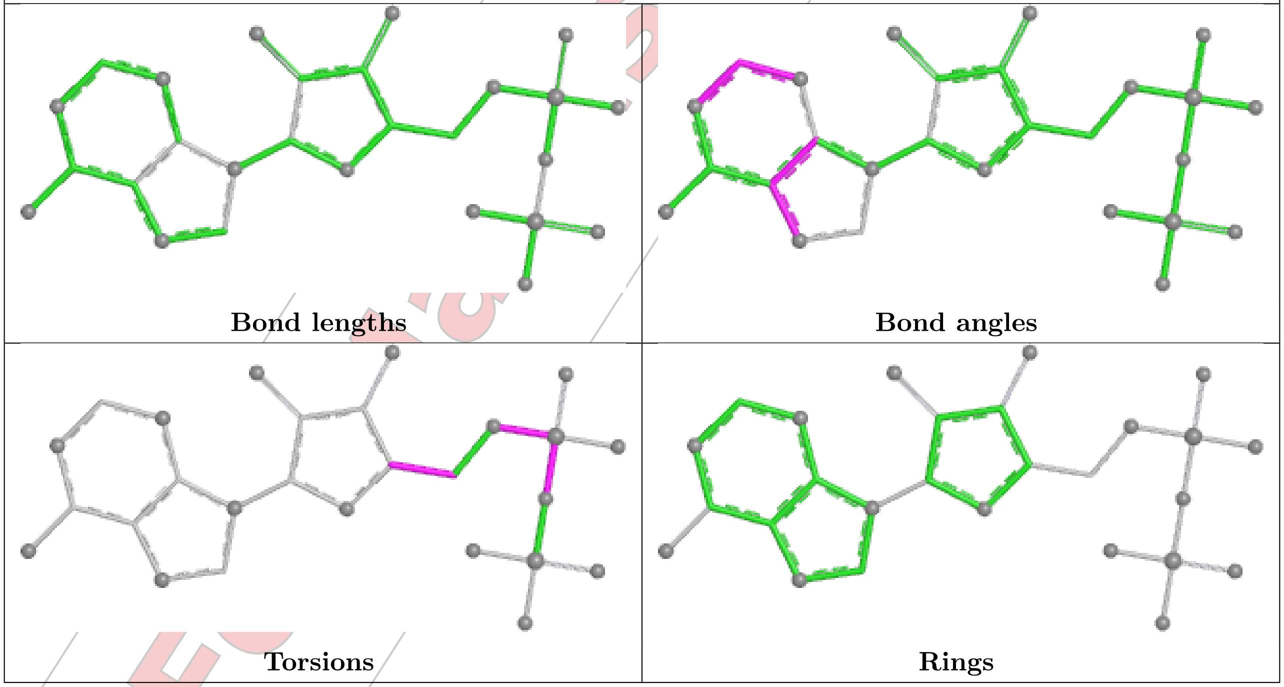


Not For Manuscript Review

##### Ligand ADP L 376

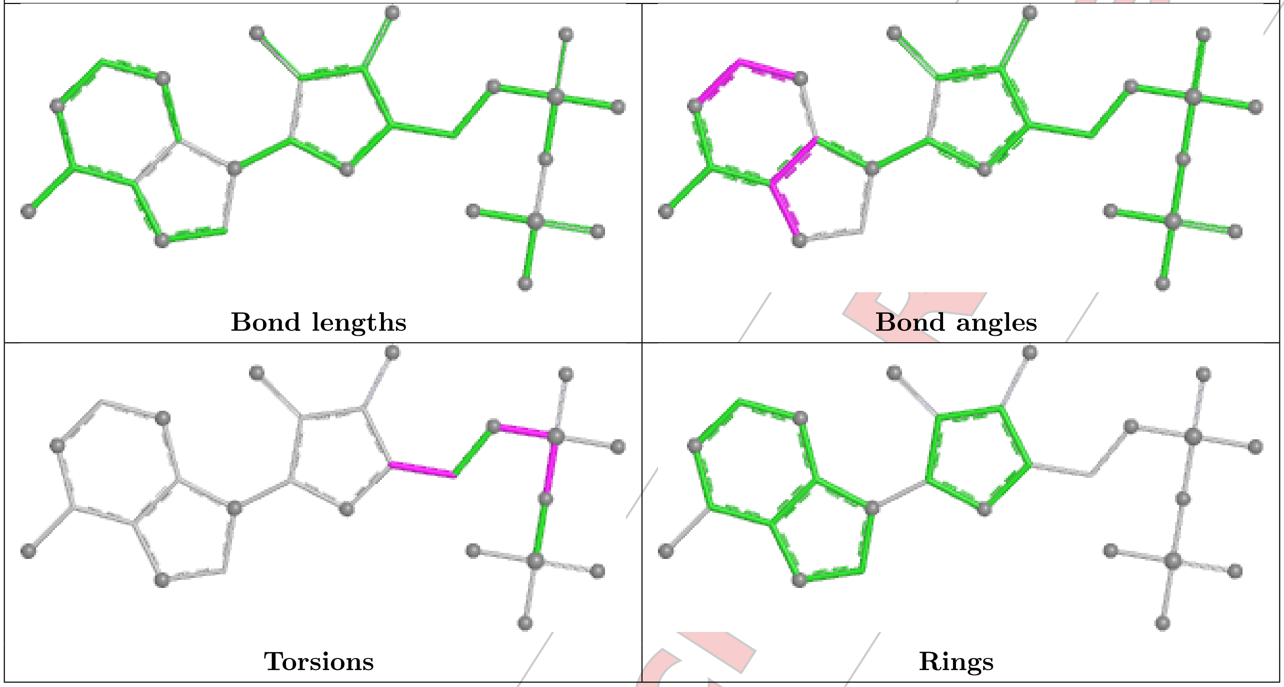


##### Ligand ADP K 376

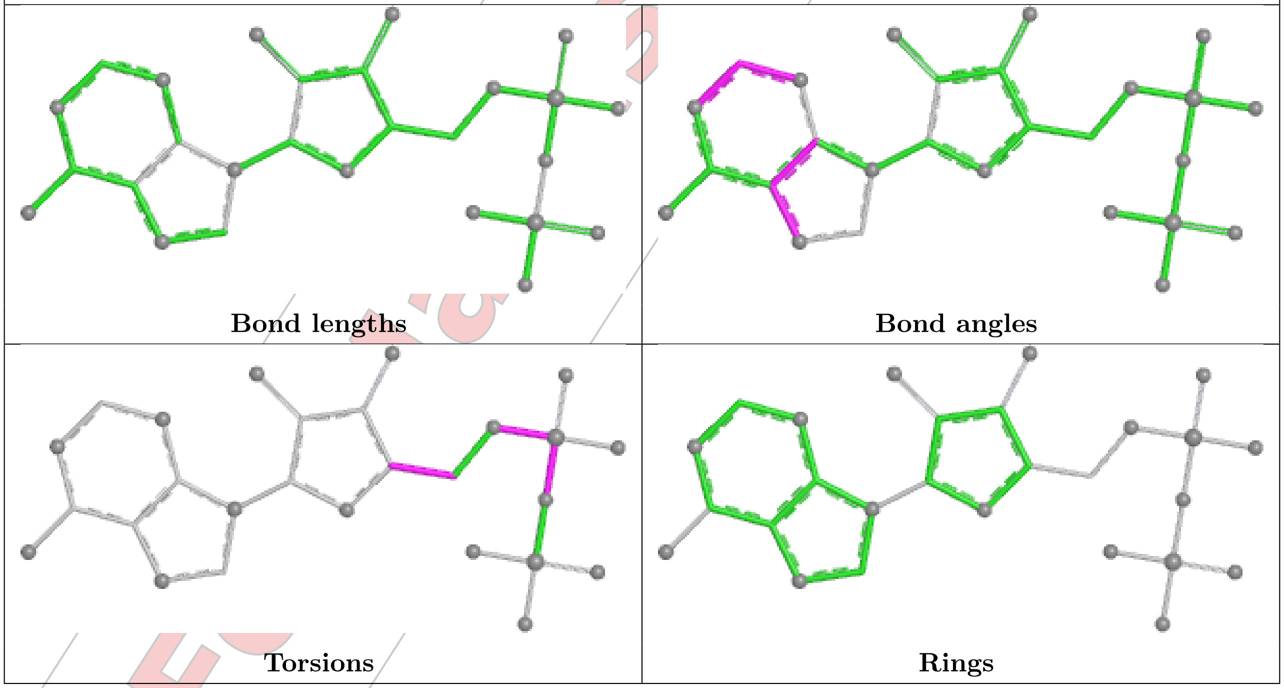


Not For Manuscript Review

**Ligand ADP H 376**

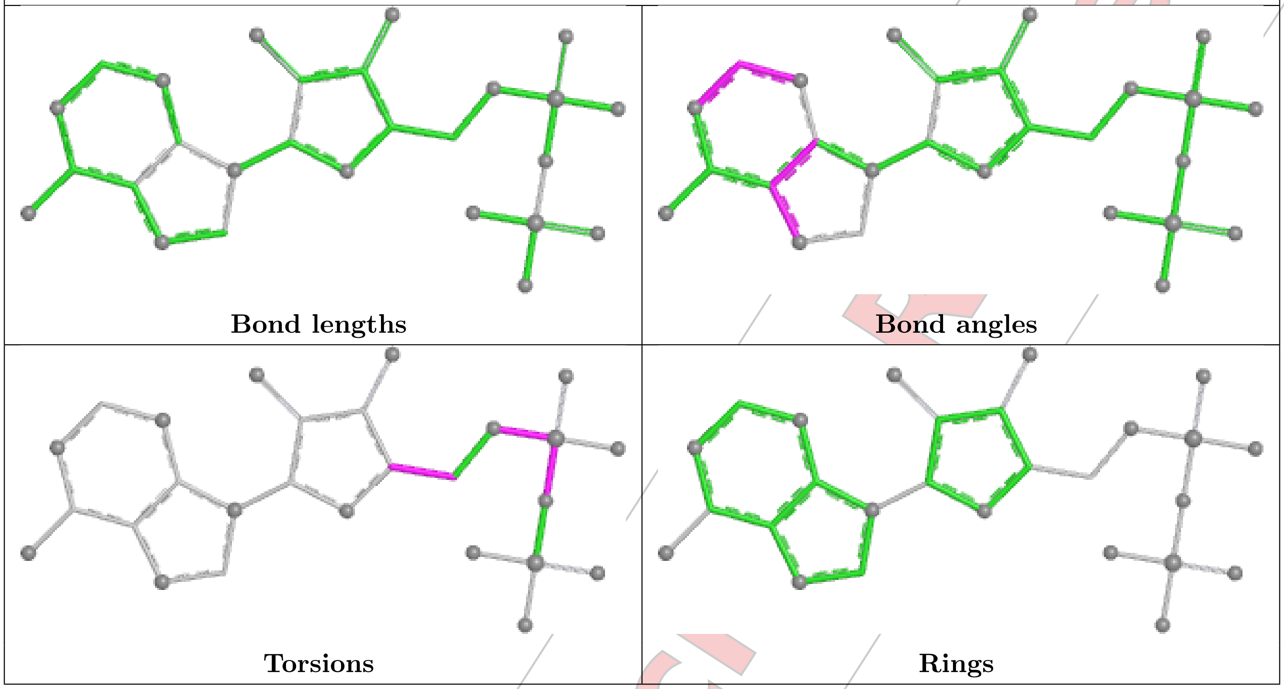


**Ligand ADP J 376**

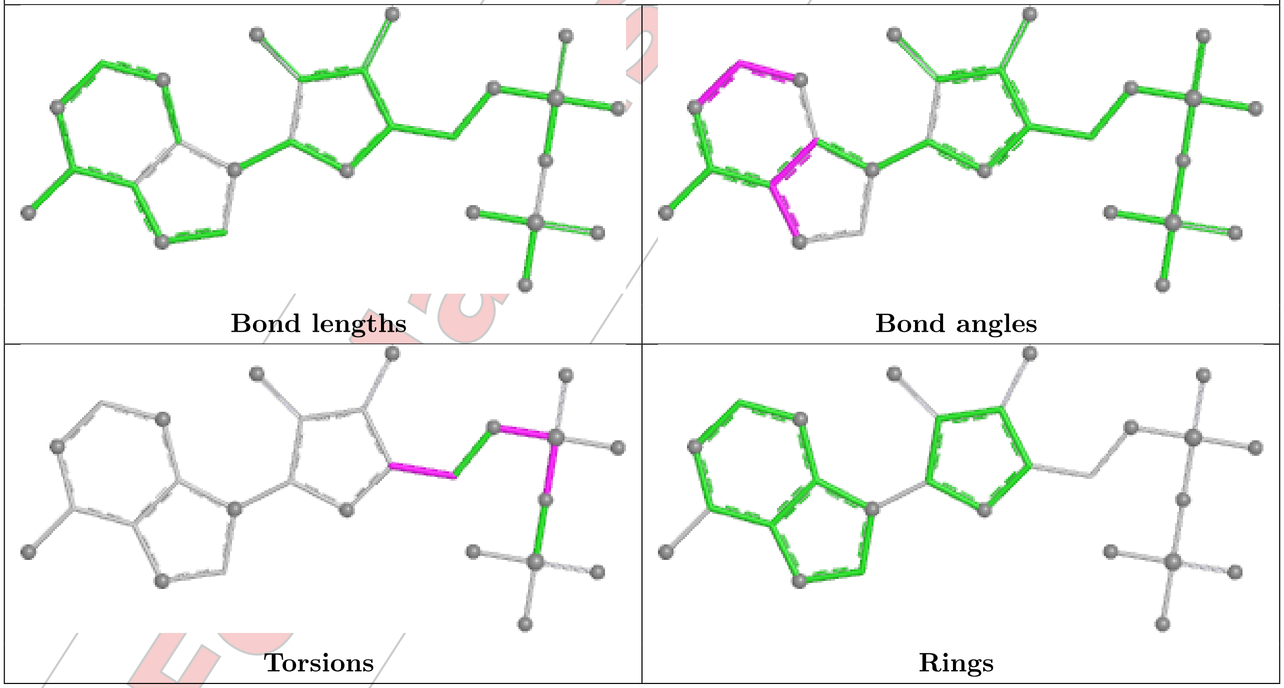


#### 5.7 Other polymers (i)

There are no such residues in this entry.

Not For Manuscript Review

#### 5.8 Polymer linkage issues (i)

There are no chain breaks in this entry.

Not For Manuscript Review

### 6 Map visualisation (i)

This section contains visualisations of the EMDB entry D_9100090603. These allow visual inspection of the internal detail of the map and identification of artifacts.

No raw map or half-maps were deposited for this entry and therefore no images, graphs, etc. pertaining to the raw map can be shown.

#### 6.1 Orthogonal projections (i)

##### 6.1.1 Primary map

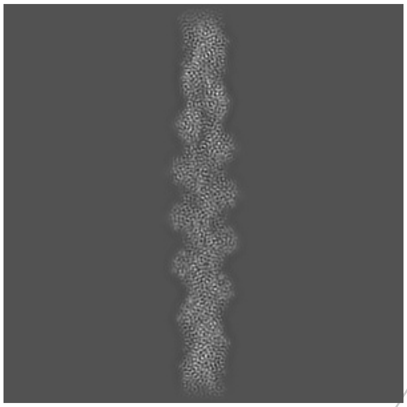

X

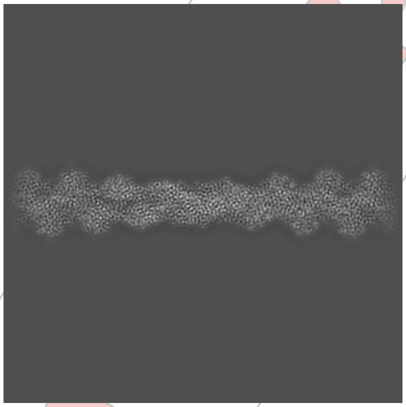

Y

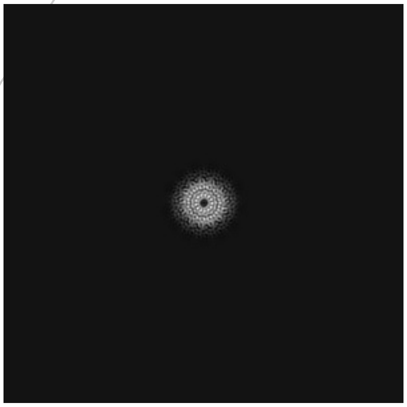

Z

The images above show the map projected in three orthogonal directions.

#### 6.2 Central slices (i)

##### 6.2.1 Primary map

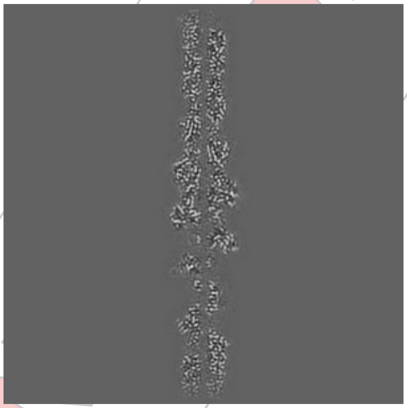

X Index: 200

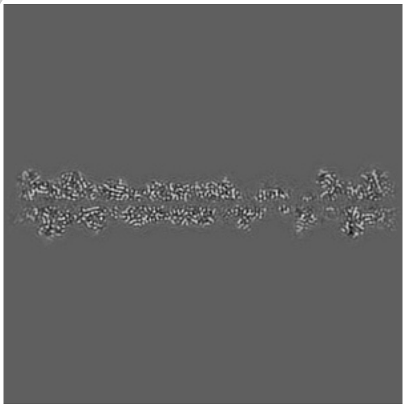

Y Index: 200

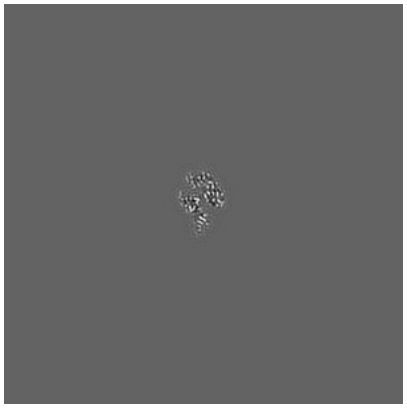

Z Index: 200

Not For Manuscript Review

The images above show central slices of the map in three orthogonal directions.

#### 6.3 Largest variance slices (i)

##### 6.3.1 Primary map

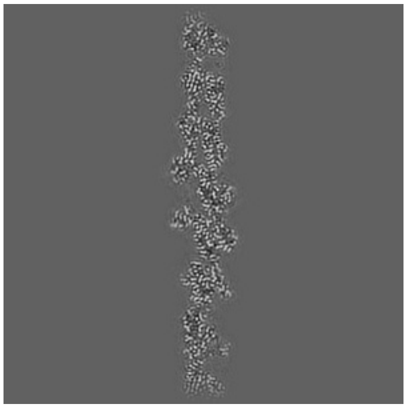

X Index: 207

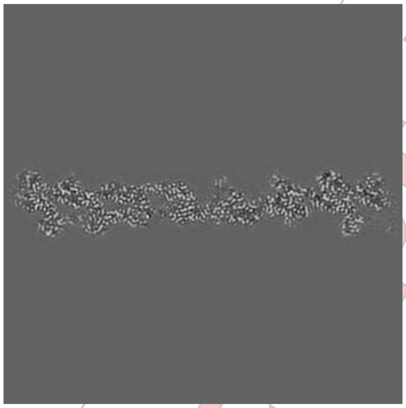

Y Index: 193

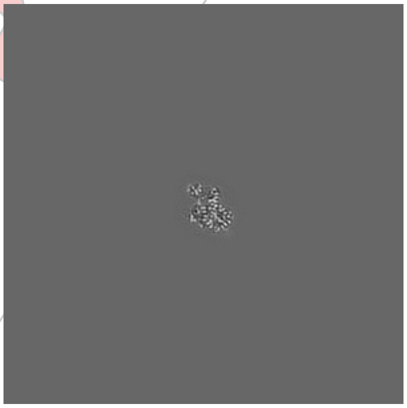

Z Index: 277

The images above show the largest variance slices of the map in three orthogonal directions.

#### 6.4 Orthogonal standard-deviation projections (False-color) (i)

##### 6.4.1 Primary map

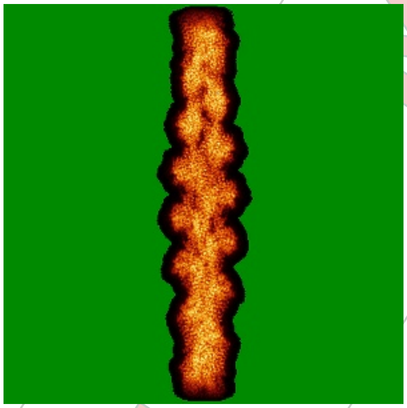

X

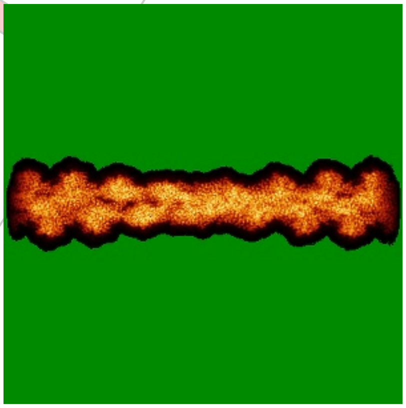

Y

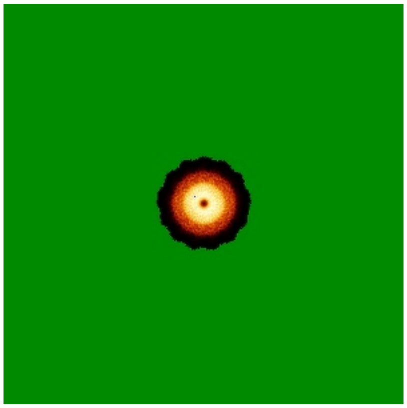

Z

The images above show the map standard deviation projections with false color in three orthogonal directions. Minimum values are shown in green, max in blue, and dark to light orange shades represent small to large values respectively.

Not For Manuscript Review

#### 6.5 Orthogonal surface views (i)

##### 6.5.1 Primary map

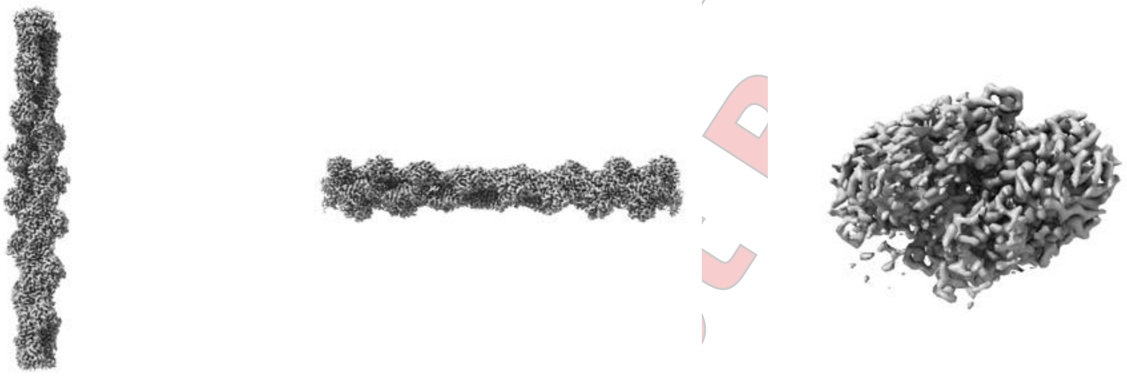

X Y Z

The images above show the 3D surface view of the map at the recommended contour level 0.9. These images, in conjunction with the slice images, may facilitate assessment of whether an appropriate contour level has been provided.

#### 6.6 Mask visualisation (i)

This section was not generated. No masks/segmentation were deposited.

Not For Manuscript Review

### 7 Map analysis (i)

This section contains the results of statistical analysis of the map.

#### 7.1 Map-value distribution (i)

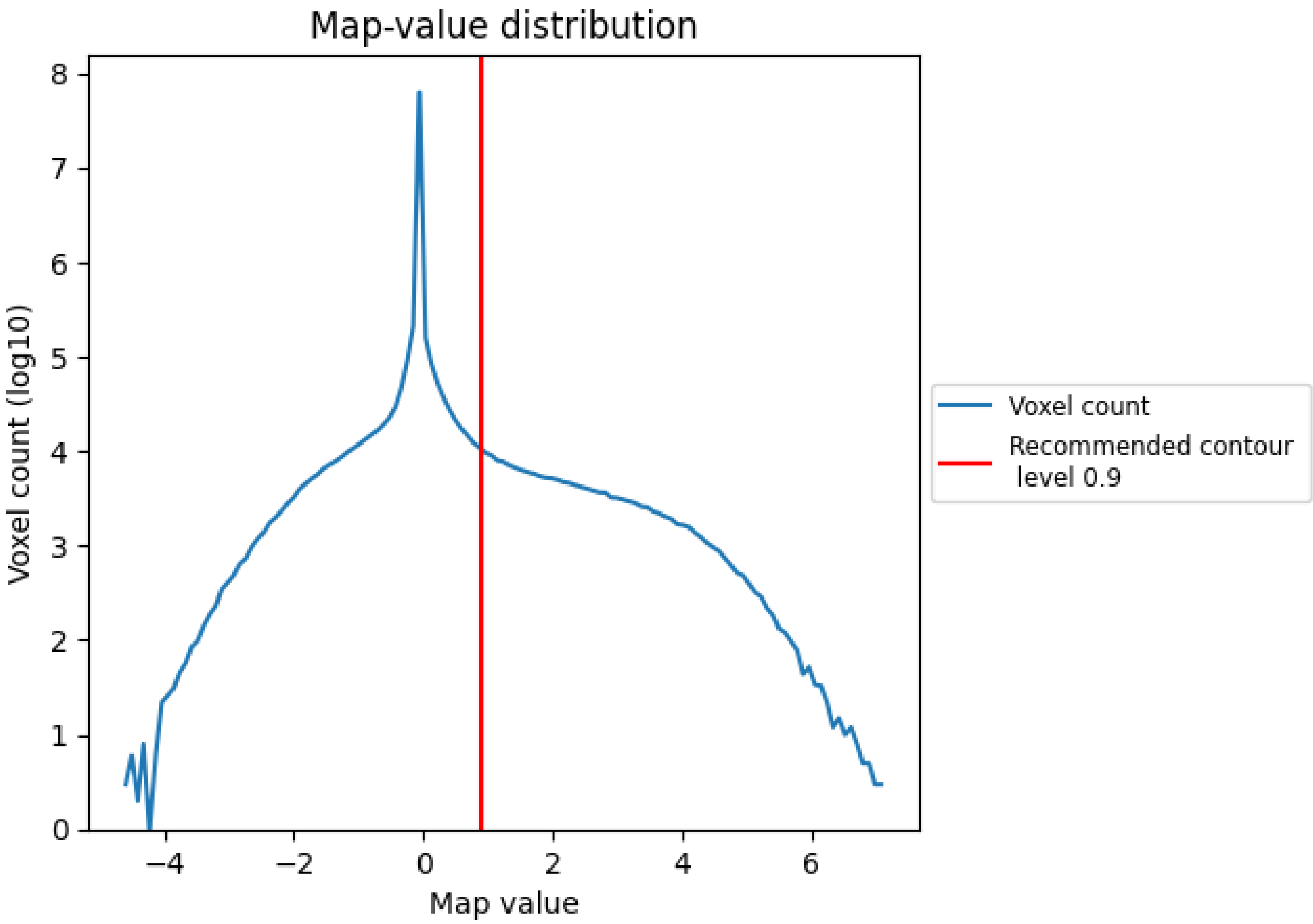


The map-value distribution is plotted in 128 intervals along the x-axis. The y-axis is logarithmic. A spike in this graph at zero usually indicates that the volume has been masked.

Not For Manuscript Review

#### 7.2 Volume estimate (i)

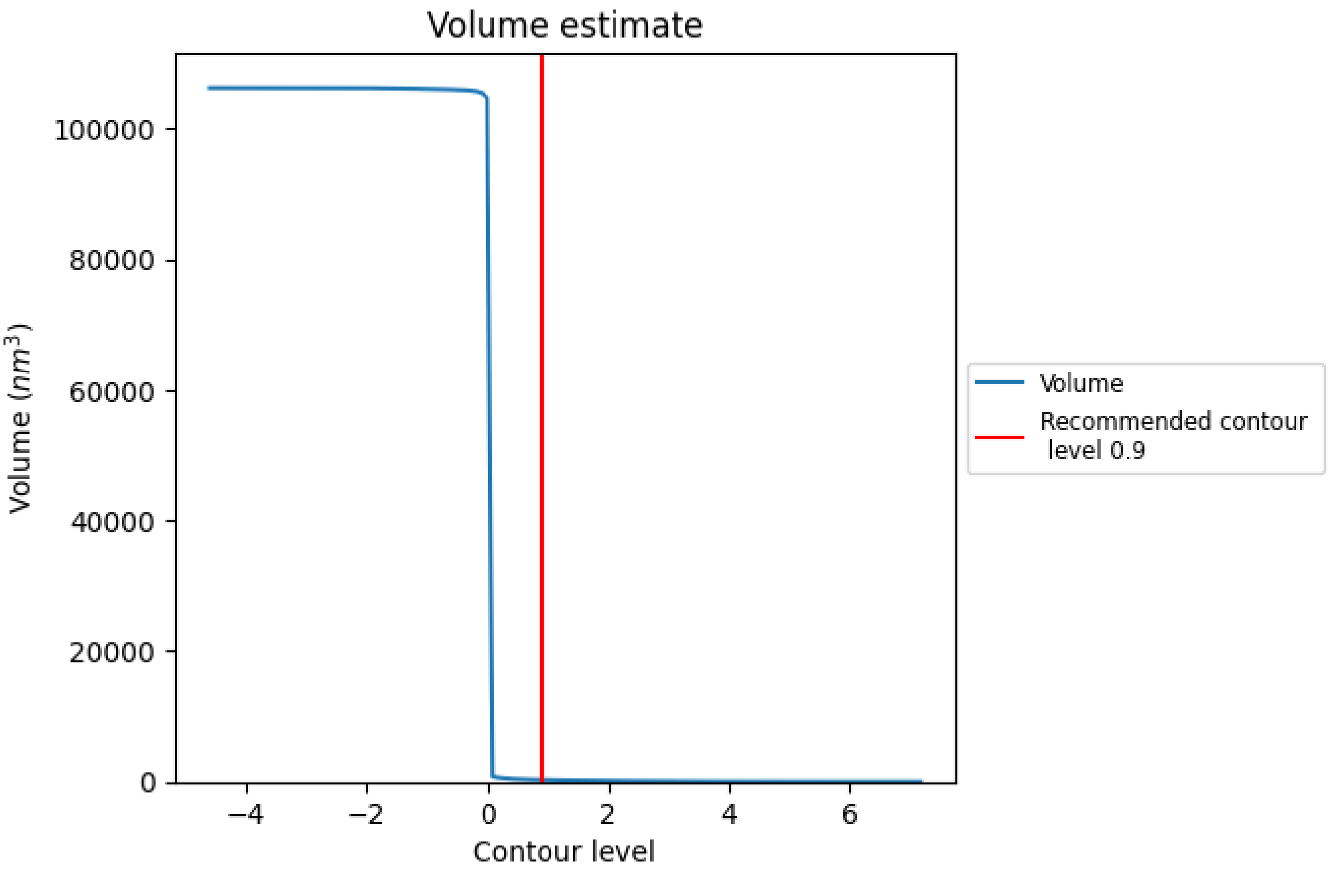


The volume at the recommended contour level is 284 $nm^3$; this corresponds to an approximate mass of 256 kDa.

The volume estimate graph shows how the enclosed volume varies with the contour level. The recommended contour level is shown as a vertical line and the intersection between the line and the curve gives the volume of the enclosed surface at the given level.

Not For Manuscript Review

#### 7.3 Rotationally averaged power spectrum (i)

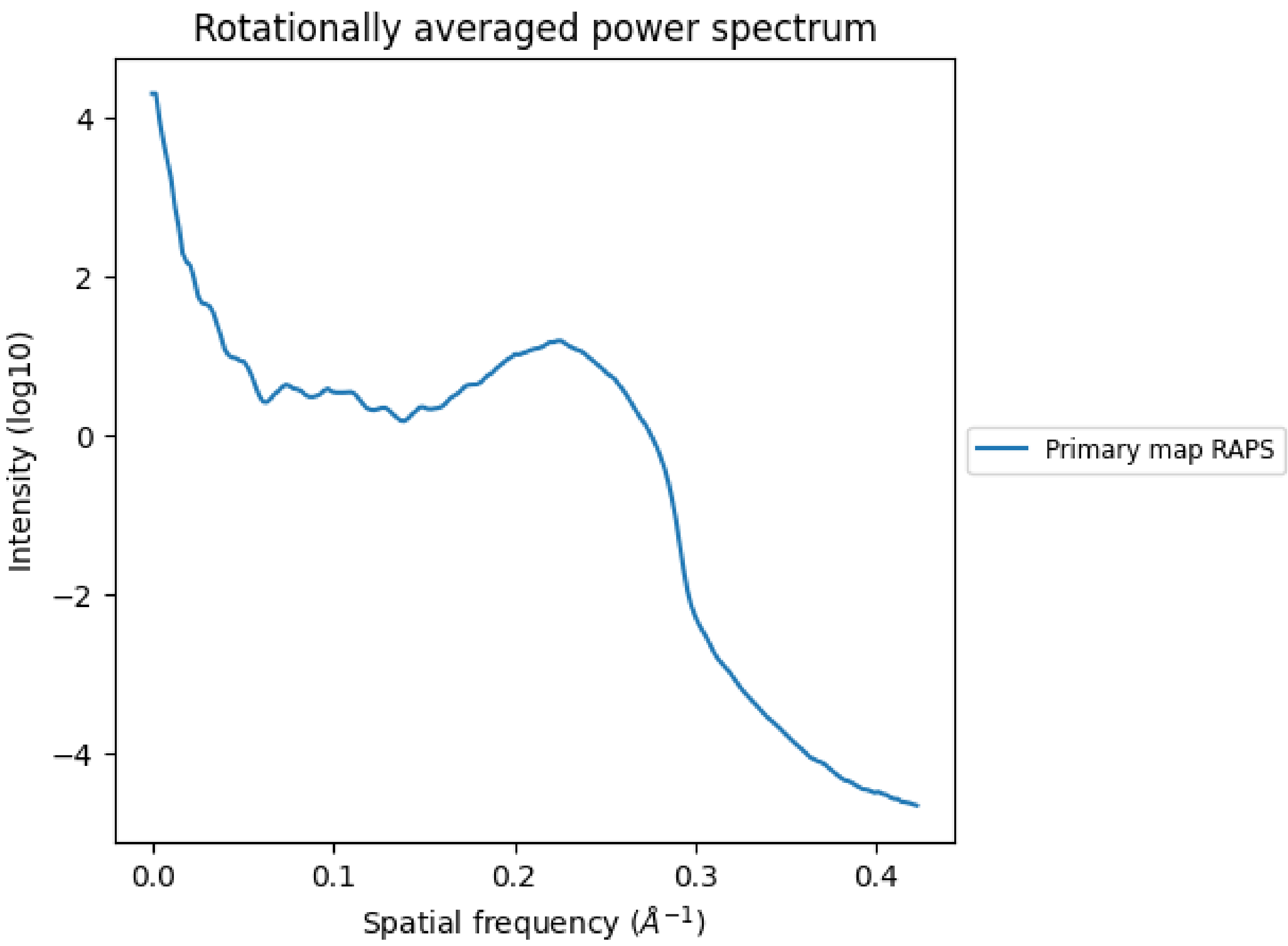


Not For Manuscript Review

### 8 Fourier-Shell correlation (i)

This section was not generated. No FSC curve or half-maps provided.

Not For Manuscript Review

### 9 Map-model fit (i)

This section contains information regarding the fit between EMDB map D_9100090603 and PDB model D_9100090603. Per-residue inclusion information can be found in section 3 on page 6.

#### 9.1 Map-model overlay (i)

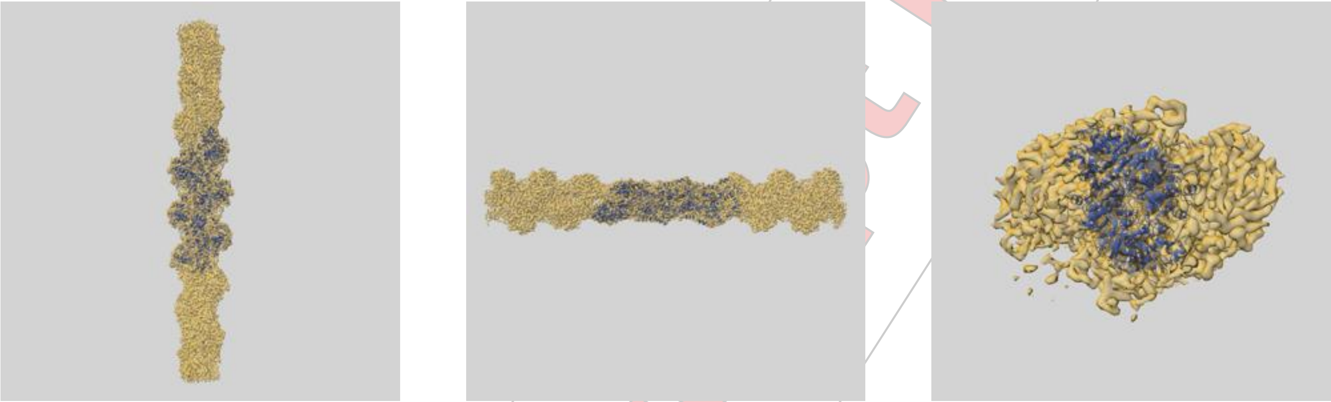

X Y Z

The images above show the 3D surface view of the map at the recommended contour level 0.9 at 50% transparency in yellow overlaid with a ribbon representation of the model coloured in blue. These images allow for the visual assessment of the quality of fit between the atomic model and the map.

Not For Manuscript Review

#### 9.2 Q-score mapped to coordinate model (i)

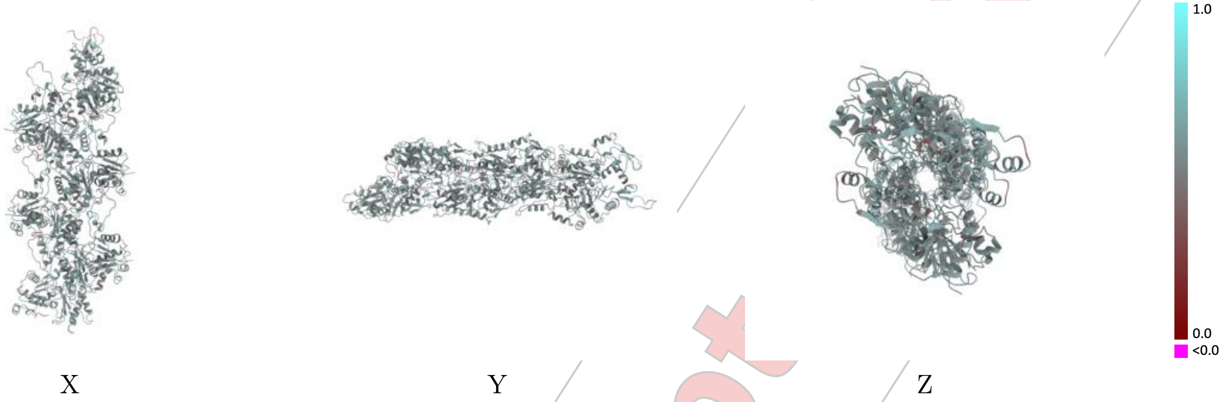


The images above show the model with each residue coloured according its Q-score. This shows their resolvability in the map with higher Q-score values reflecting better resolvability. Please note: Q-score is calculating the resolvability of atoms, and thus high values are only expected at resolutions at which atoms can be resolved. Low Q-score values may therefore be expected for many entries.

#### 9.3 Atom inclusion mapped to coordinate model (i)

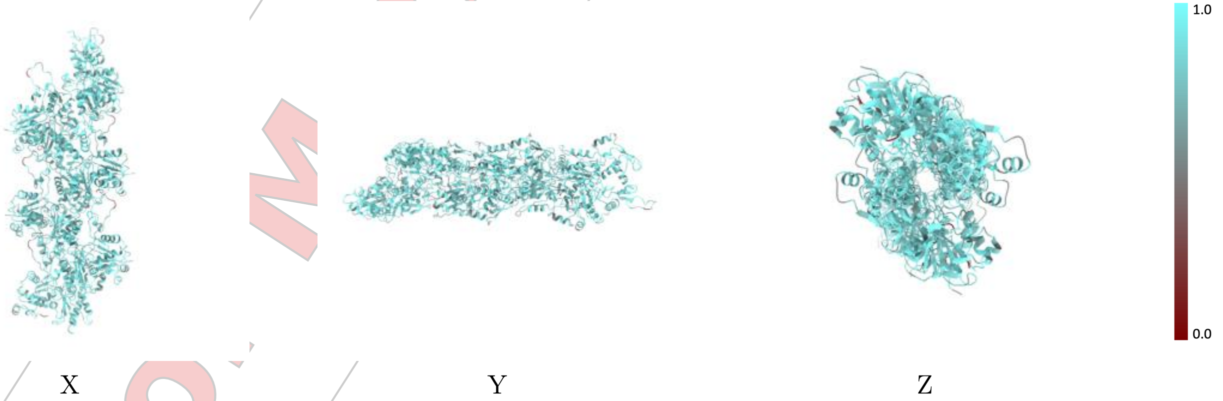


The images above show the model with each residue coloured according to its atom inclusion. This shows to what extent they are inside the map at the recommended contour level (0.9).

Not For Manuscript Review

#### 9.4 Atom inclusion (i)

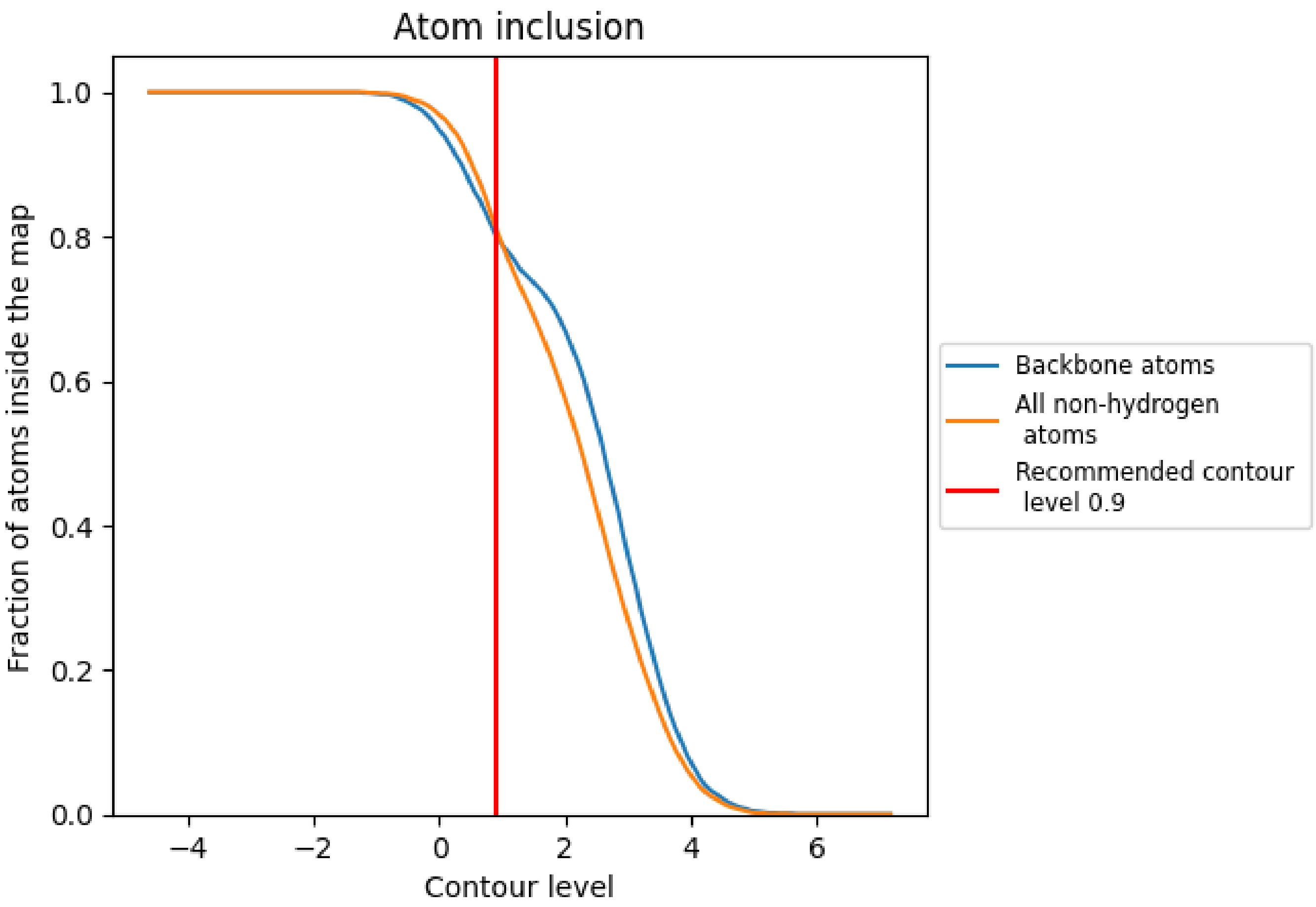


At the recommended contour level, 80% of all backbone atoms, 81% of all non-hydrogen atoms, are inside the map.

Not For Manuscript Review

#### 9.5 Map-model fit summary (i)

The table lists the average atom inclusion at the recommended contour level (0.9) and Q-score for the entire model and for each chain.

| Chain | Atom inclusion | Q-score |
|---|---|---|
| All | 0.8130 | 0.5310 |
| A | 0.8150 | 0.5320 |
| B | 0.8160 | 0.5320 |
| C | 0.8080 | 0.5290 |
| D | 0.8120 | 0.5300 |
| E | 0.8100 | 0.5310 |
| F | 0.8130 | 0.5340 |
| G | 0.9290 | 0.5460 |
| H | 0.9640 | 0.5560 |
| I | 0.9290 | 0.5720 |
| J | 0.9290 | 0.5310 |
| K | 0.8930 | 0.5410 |
| L | 0.8930 | 0.5630 |
| M | 0.7780 | 0.4790 |
| N | 0.7960 | 0.4940 |
| O | 0.7960 | 0.5010 |
| P | 0.7780 | 0.4950 |
| Q | 0.7780 | 0.4900 |
| R | 0.7960 | 0.4930 |
| S | 0.8150 | 0.5000 |

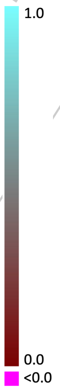


Not For Manuscript Review
